# Improving deep models of protein-coding potential with a Fourier-transform architecture and machine translation task

**DOI:** 10.1101/2023.04.03.535488

**Authors:** Joseph D. Valencia, David A. Hendrix

## Abstract

Ribosomes are information-processing macromolecular machines that integrate complex sequence patterns in messenger RNA (mRNA) transcripts to synthesize proteins. Studies of the sequence features that distinguish mRNAs from long noncoding RNAs (lncRNAs) may yield insight into the information that directs and regulates translation. Computational methods for calculating protein-coding potential are important for distinguishing mRNAs from lncRNAs during genome annotation, but most machine learning methods for this task rely on previously known rules to define features. Sequence-to-sequence (seq2seq) models, particularly ones using transformer networks, have proven capable of learning complex grammatical relationships between words to perform natural language translation. Seeking to leverage these advancements in the biological domain, we present a seq2seq formulation for predicting protein-coding potential with deep neural networks and demonstrate that simultaneously learning translation from RNA to protein improves classification performance relative to a classification-only training objective. Inspired by classical signal processing methods for gene discovery and Fourier-based image-processing neural networks, we introduce LocalFilterNet (LFNet). LFNet is a network architecture with an inductive bias for modeling the three-nucleotide periodicity apparent in coding sequences. We incorporate LFNet within an encoder-decoder framework to test whether the translation task improves the classification of transcripts and the interpretation of their sequence features. We use the resulting model to compute nucleotide-resolution importance scores, revealing sequence patterns that could assist the cellular machinery in distinguishing mRNAs and lncRNAs. Finally, we develop a novel approach for estimating mutation effects from Integrated Gradients, a backpropagation-based feature attribution, and characterize the difficulty of efficient approximations in this setting.

## 1 Introduction

The flow of genetic information from DNA to RNA to protein is a fundamental life process in which messenger RNAs (mRNAs) act as the information-carrying intermediaries. High-throughput sequencing has revealed the abundance of another class of RNA called long noncoding RNAs (lncRNAs), which share important biochemical features such as 5’ capping and polyadenylation with protein-coding mRNAs (Iyer et al. 2015). Long noncoding RNAs are differentiated from smaller noncoding RNAs like tRNAs and microRNAs based on their greater length of at least 200 nucleotide (nt), and from mRNAs based on limited evidence of lncRNA protein expression and sequence conservation (Derrien et al. 2012). LncRNAs make up more than 68% of the human transcriptome and play important regulatory roles, particularly during development (Statello et al. 2021; Ransohoff et al. 2018). They are implicated in numerous diseases including cancer and cardiovascular disease (Sallam et al. 2018).

The protein-coding potential of many transcripts is unresolved, and many transcripts previously or currently annotated as lncRNAs are mislabeled and in fact possess small open reading frames (sORFs) that encode micropeptides (Choi et al. 2019). Ribosome profiling (Ribo-Seq) shows that ribosomes bind readily to lncRNAs (Ingolia, Lareau, et al. 2011), though the ribosome does not interact with lncRNA ORFs in the same way as mRNAs, lacking a distinctive drop-off of Ribo-Seq coverage at ORF end (Guttman et al. 2013). Ribo-Seq protocols accounting for the 3-nt periodicity of ribosome footprint density (Guo et al. 2010) have identified some genuine sORF translation (Ingolia, Brar, et al. 2014; Ji et al. 2015). Only a small fraction of the possible set of micropeptides encoded by transcripts currently annotated as lncRNAs have been directly detected via mass spectrometry, leaving the vast majority as presumptively nonfunctional or rapidly degraded (Housman and Ulitsky 2016; Bánfai et al. 2012; Verheggen et al. 2017). Still, hundreds of lncRNAs have been confirmed to be misannotated, and these transcripts do encode micropeptides, for example, myoregulin, a 46-aa. regulator of Ca^2+^ activity that contributes to muscle cell performance (Anderson et al. 2015). Micropeptides are also involved in metabolism, red blood cell development, cardiomyocyte hypertrophy (Yan et al. 2021), inflammation, tumorigenesis and tumor suppression (Othoum et al. 2020; Wu et al. 2020), and more (Hartford and Lal 2020).

Such uncertainty as to the intrinsic protein-coding potential of ORFs raises the question of how cells distinguish true coding regions, with the translational machinery likely to play a critical role. Recent results suggest that general sequence features governing the kinetics of protein synthesis also separate mRNA and untranslated lncRNA ORFs more broadly (Patraquim et al. 2022). The Kozak consensus sequence is well-characterized as the optimal context for translation initiation, and ribosomes can skip unfavorable AUGs through leaky scanning (Kozak 1987; Kozak 2002). Initiation can be affected by cis-regulatory features such as 5’ UTR secondary structure (JJ Li et al. 2019) and upstream ORFs (Johnstone et al. 2016), and by trans-acting factors such as microRNAs (Guo et al. 2010) and RNA-binding proteins (Szostak and Gebauer 2013). Codon usage biases in the 5’-most region of the CDS are particularly known to affect the elongation rate during protein synthesis (Tuller et al. 2010; Verma et al. 2019; Subramanian et al. 2021).

Distinguishing between mRNAs and lncRNAs is an important step in annotating newly sequenced genomes, and a variety of statistical and computational methods have been developed for this task. Codon Adaptive Index (CAI) (Sharp and WH Li 1987) discriminates coding nucleic acids according to biases in the synonymous codons that code for each amino acid and Fickett scores (Fickett 1982) by the nucleotides present in the three codon positions. Early computational approaches used Fourier or wavelet analysis to identify coding sequences (CDS) from their characteristic periodicity of nucleotide identity induced by codon usage bias (Tiwari et al. 1997; Anastassiou 2000; Deng et al. 2010; Hassani Saadi et al. 2017). Machine learning methods have been designed around features such as the absolute length of ORFs, ORF length relative to the transcript, codon and hexamer frequencies including Coding Potential Assessment Tool (CPAT) (Wang et al. 2013) and coding potential calculator (Kong et al. 2007), and others (A Li et al. 2014; Wucher et al. 2017).

Although many prior machine learning methods achieve high classification performance, they typically rely on transcript-level summary features. Deep learning approaches can operate directly on sequences without such intermediate features and have proven effective in predicting properties of biological sequences, including a wide variety of functional genomics assays (Avsec, Agarwal, et al. 2021; Tareen et al. 2022), RNA splicing (Zeng and YI Li 2022) and degradation (Agarwal and Kelley 2022), and protein structure (Jumper et al. 2021). A recent method called RNAsamba uses a convolutional neural network variant to achieve high performance from both nucleotide and amino acid sequence, but also relies on pre-defined features such as the longest ORF (Camargo et al. 2020). A critical limitation in the development of intelligent systems for classifying transcripts as protein-coding vs noncoding is the bias of using the translation and length of the longest ORF in machine learning approaches. Our group previously developed mRNN, the first recurrent neural network classifier of coding RNA from primary sequences alone (Hill et al. 2018). There is a need for more flexible neural networks capable of learning sequence-specific rules that promote translation to better understand what drives translational efficiency. The advantage of these approaches is that they do not require feature engineering, and are capable of learning new biological rules that are encapsulated in the weights of the neural network. Interpretation of these deep neural networks can lead to the identification of new sequence features that are informative for the evaluation of biological sequences and understanding the regulation of translation. Interpreting deep models is challenging, but a significant literature in explainable artificial intelligence (xAI) has arisen in regulatory genomics, with notable successes in uncovering transcription factor binding logic (Avsec, Weilert, et al. 2021; Novakovsky et al. 2022). Interpretation of similar deep models of protein coding potential could help identify new sequence features regulating translation.

In this paper, we describe bioseq2seq, a novel neural network model of biological translation based on the sequence-to-sequence (seq2seq) paradigm commonly used for machine translation of human languages. Although the genetic code follows a well-understood mapping between nucleic acid codons and amino acids, we demonstrate that learning to predict the protein sequence from the sequence of its message improves neural network performance in distinguishing mRNAs from lncRNAs. Adapting recent advances in token mixing neural architectures, we introduce Local Filter Network (LFNet), a computationally efficient network layer based on the short-time Fourier transform. We leverage perturbation-based feature importance values to extract sequence patterns which impact the model prediction and generate hypotheses about the regulatory elements that could differentiate coding RNA *in vivo*. Lastly, we offer evidence that while our LFNet-based bioseq2seq model robustly uncovers biological rules to learn protein-coding potential, it presents challenges for approximate interpretation techniques in deep learning. We address these challenges by introducing mutation-directed integrated gradients (MDIG), which we show has a strong correlation with synonymous sequence perturbations, and can be used to identify regions in transcripts that are important for defining protein-coding potential.

## 2 Results

### 2.1 Translation training objective improves classification performance

We downloaded lncRNA primary sequences and mRNAs matched with their encoded proteins from the NCBI RefSeq annotations of eight mammalian species. Following the encoder-decoder framework widely used in sequence-to-sequence learning, we trained two major types of deep learning models on this dataset. The primary is bioseq2seq, which outputs a class prediction of *(NC)* for lncRNAs or *(PC)* followed by a predicted protein sequence for coding mRNAs. To test the benefits of a translation-based learning objective, we trained a secondary encoder-decoder model type to predict only the RNA class and not its translation, which we called Encoder Decoder Classifier (EDC). The common architectural framework enables a fair comparison between these two training settings. We designed a novel neural network layer, LFNet, to efficiently apply a short-time (local) Fourier transform to the high-dimensional vectors representing each input nucleotide and perform sequential updates via frequency-domain filtering. Several LFNet layers were composed into an encoder stack to process the RNA. A stack of transformer decoders operates on the encoder hidden representations to produce an output, autoregressively consuming its own predictions to produce the next character, as necessary (Vaswani et al. 2017). Within this general framework, summarized in Fig 1, we optimized several hyperparameters, including hidden dimension and number of encoder and decoder layers, for bioseq2seq and EDC separately (Supplementary Table 1). Bioseq2seq performance was optimized with 12 LFNet encoder and 2 transformer decoder layers, while EDC selected 16 of each for a substantially larger model. After optimizing the hyperparameters for bioseq2seq and EDC, we trained four model replicates of each from different random initializations. We also trained replicates for the EDC task using the optimal hyperparameters for bioseq2seq, referring to this as EDC-small, in contrast to the optimized EDC, which we refer to as EDC-large. We report the classification accuracy on a withheld test set for our two model types in Table 1. In the case of bioseq2seq, which produces a variable-length peptide decoding at inference time, decoding was halted after the leading classification token was predicted. The bioseq2seq replicate with the best performance on F1 score achieved a score of 0.958, while the worst-performing on this metric scored 0.947. We compared our models with five replicates of RNAsamba trained on our dataset, as well as CPC2 and CPAT, two machine-learning methods based on engineered features. The best model for bioseq2seq exceeds the performance of CPC2 and CPAT and is competitive with RNAsamba (0.956-0.961 F1) without explicit inclusion of any auxiliary features such as ORF k-mers, although RNAsamba appears slightly better according to all evaluation metrics except recall. EDC-large ranged in performance between 0.924-0.932 in F1. EDC-small was clearly the worst of all models and so from this point we will only consider EDC-large and refer to it simply as EDC. The markedly better performance of bioseq2seq in comparison to its classification-only analogues makes it clear that the translation task improves the performance of an LFNet model on the binary classification task.

**Figure 1.**
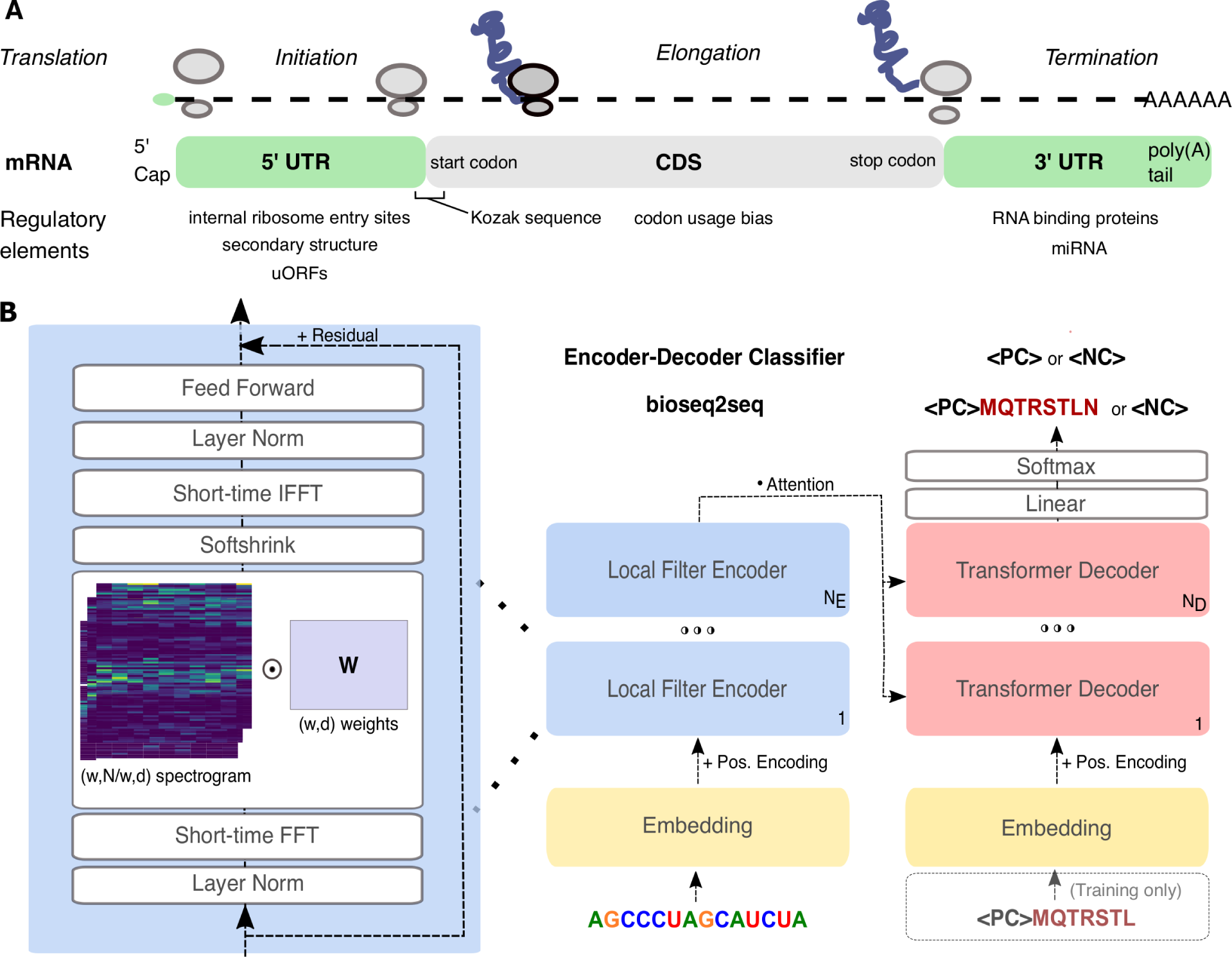
Overview of problem setting and computational method. (A) Summary of messenger RNA functional regions and known elements regulating translation. See (Gebauer and Hentze 2004) for a review of known regulatory elements. (B) Neural network sequence-to-sequence architecture. We designed LFNet (left) to apply a learned filter matrix *W* to a 1D short-time Fourier transform (spectrogram) of the hidden representations, enabling frequency-domain filtering of the 3-base periodicity present in coding sequences. We trained this architecture for two problem settings: in Encoder-Decoder Classifier (EDC), the expected output is a classification token, for bioseq2seq, the protein translation is also predicted.

**Table 1.** Classification Performance. Bioseq2seq was compared with an EDC model whose hyperparameters were tuned independently (large) and an EDC model with identical hyperparameters to bioseq2seq (small). Several top-performing machine learning models were evaluated on our dataset for comparison. For our models, predictions were made using the leading ‘classification’ token *(PC)* or *(NC)* of the first beam, terminating inference before the peptide prediction. For our models and RNAsamba, multiple replicates were trained with different random seeds. Evaluation metrics were calculated with *(PC)* as the positive class and listed as mean *±* std. dev. where multiple replicates are available.

| Model | Accuracy | F1 | Recall | Precision | MCC |
| --- | --- | --- | --- | --- | --- |
| EDC (small) | 0.885 ± 0.017 | 0.896 ± 0.014 | 0.913 ± 0.011 | 0.880 ± 0.024 | 0.769 ± 0.034 |
| EDC (large) | 0.922 ± 0.004 | 0.927 ± 0.003 | 0.910 ± 0.011 | 0.945 ± 0.014 | 0.845 ± 0.009 |
| bioseq2seq | 0.950 ± 0.006 | 0.954 ± 0.005 | 0.953 ± 0.012 | 0.955 ± 0.017 | 0.900 ± 0.011 |
| RNAsamba | 0.957 ± 0.002 | 0.960 ± 0.002 | 0.949 ± 0.004 | 0.970 ± 0.002 | 0.913 ± 0.004 |
| CPAT | 0.939 | 0.944 | 0.947 | 0.940 | 0.876 |
| CPC2 | 0.911 | 0.912 | 0.856 | 0.976 | 0.830 |

**Table 2.**
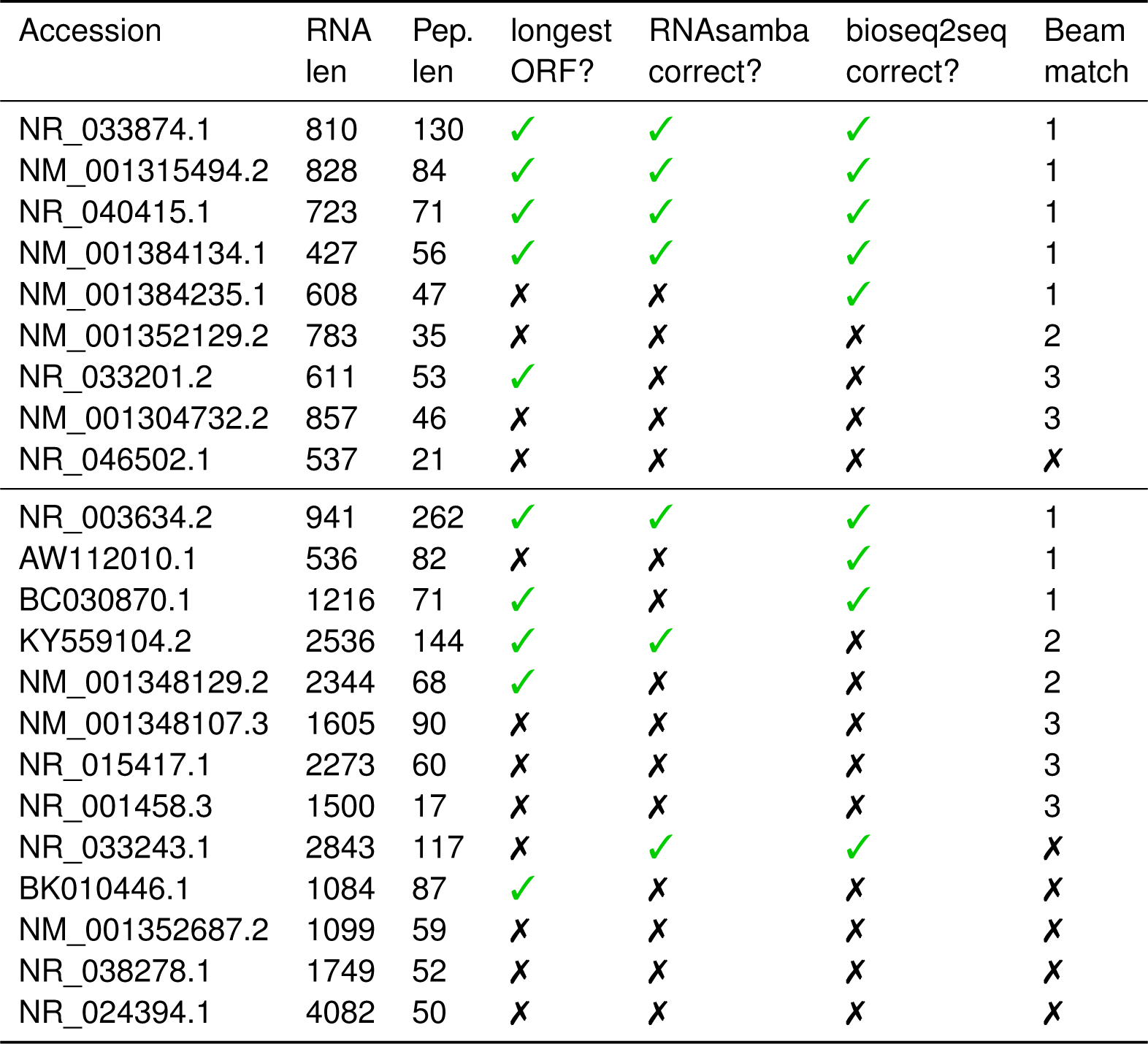
Results on twenty-two validated micropeptides. Samples above the horizontal bar were in our training set and those below were not. A bioseq2seq prediction was counted as correct if it began with *(PC)*, regardless of the official class label. The matching beam indicates the first beam peptide decoding from bioseq2seq achieving *≥* 90% alignment with the annotated micropeptide, if one exists.

As bioseq2seq is capable of performing translation on top of classification, we also report the percentage identity between the ground truth protein and the translation produced by bioseq2seq using the Needleman-Wunsch global alignment. A large majority, 82.4 %, are exact matches with the ground truth. Notably, when bioseq2seq was allowed to predict a full-length protein rather than halted after the classification token as in the results from the previous section, the classification performance of the best model deteriorated slightly to 0.940 F1. This suggests a slight trade-off at inference time between an accurate peptide decoding and the classification task, though the bioseq2seq training strategy as a whole clearly improves classification performance relative to EDC.

### 2.2 Alternate decodings of lncRNAs harbor plausible micropeptides

The bioseq2seq formulation can produce and rank multiple candidate decodings for a given RNA using beam search. For sequences annotated as lncRNAs and correctly classified by bioseq2seq, the lower beams (second highest scoring and on) will with high probability begin with *(PC)*. We investigated the predicted peptides for insights into the potential translation of lncRNAs. First we confirmed that the peptides matched a true ORF within the lncRNA by using the EMBOSS package to find the top Needleman-Wunsch alignment score between the three-frame translation and all generated peptides from a beam size of four (Rice et al. 2000). In 59.7 % of cases, the best match was a perfect alignment, meaning that most peptide decodings were translations of ORFs actually present in the lncRNA.

We applied bioseq2seq to a set of transcripts previously or currently annotated as lncRNAs but considered by the database lncPEP to have been validated by supporting literature to express a micropeptide (Liu et al. 2022). Starting from the lncPEP "validated" set, we implemented a number of quality control measures, removing redundant transcripts, linking the transcript names listed on the lncPEP website with RefSeq accession numbers via the underlying primary literature and the NCBI search function. This yielded twenty-two putative micropeptide-encoding transcripts (provided as Supplementary Table 3), of which nine were found in our training set. The best model for RNAsamba predicted 3 of the remaining 13 to be protein coding. Using bioseq2seq, 3 were also predicted as coding when terminating inference after the classification token, and 4 when running peptide decoding to completion.

We aligned all beams from bioseq2seq with the lncPEP micropeptides and found that in most cases the model also successfully identified the correct ORF to translate, with 3 of 4 predicted coding transcripts having alignment identity *≥* 90%. If lower beams are considered, 8 have identity *≥* 90%, including a very short 17-aa peptide. The examples found in our training set are of potential interest as well because in several cases the class label that we trained on contradicts the prediction that bioseq2seq makes. For example, *LINC00266-1* with NCBI accession NR_040415.1 is currently annotated as a lncRNA but was found in (Zhu et al. 2020) to express a 71-amino acid oncopeptide. Bioseq2seq perfectly predicts the peptide in its highest beam – a false positive according to the class label in the training data. Examples like this and NR_033874.1 highlight the generalizability of the rules learned by bioseq2seq and RNAsamba, even when presented with false annotations. One example, NM_001384235.1 in the training set underscores a crucial distinction between bioseq2seq and prior methods like RNAsamba. In these transcripts, the micropeptide is not coded for by the longest ORF. RNAsamba only explicitly considers the longest ORF in each transcript and may fail to identify alternate sources of coding potential, as it does here. The translation product for AW112010.1 in the test set comes from an instance of non-AUG initiation (Jackson et al. 2018), and while our method cannot perfectly predict the protein product in such cases we successfully identify it as a coding transcript and predict a partial match from the canonical portion of the CDS.

### 2.3 Local Filter Networks emphasize 3-nt periodicity

The core feature of each LFNet layer is its learned frequency-domain filters. We visualized the filter weight matrices to investigate the frequency response of the model to signals in the intermediate vector representations, including separate plots for their magnitude *|z|* and phase *θ* for the complex weights *z* = *|z|e^iθ^*. The resulting images for all layers in both bioseq2seq and EDC are given in Figure 3. Visually, the most prominent signal in both model types is a band at a frequency bin equivalent to a period of 3 nt. This illustrates that most layers and hidden dimension across the LFNet stack learned to emphasize the 3-base periodicity of coding regions. Notably, every layer of EDC (panel B) shows a more clear dependence on the 3-nt property than bioseq2seq (panel A), with every layer having a clean visual band of low magnitudes along this frequency range. In contrast, lower layers of bioseq2seq do not appear to emphasize this feature. However, bioseq2seq has phase values close to zero along the 3nt band (panel C), while the phase activity of EDC is somewhat more random (panel D). We observed in Supplementary Fig S1 that for bioseq2seq, the periods other than 3-nt are associated with phases peaked around *−π* and *π*, which correspond to phase components of the weights being *e^iθ^* = *−*1, such that the output of the LFNet layer would negate the residual when they are added. While the bioseq2seq LFNet weights shift toward the positive real-axis in the higher layers for three-nucleotide signals, they shift toward the negative real axis for other periods. This trend is found clearly in bioseq2seq, and less so EDC, where the weights are smaller and more centered at zero (Supplementary Fig S2). Furthermore, while weights corresponding to three-nucleotide signals are mostly zero for EDC, creating a band in Fig 3, the weaker band for bioseq2seq is explained by many positive weights in bioseq2seq at this band, which would amplify three nucleotide signals. We hypothesize that the inductive bias of LFNet facilitates a reliance on the 3-base property, and the translation task leads to the amplification of specific 3-base signals.

**Figure 2.**
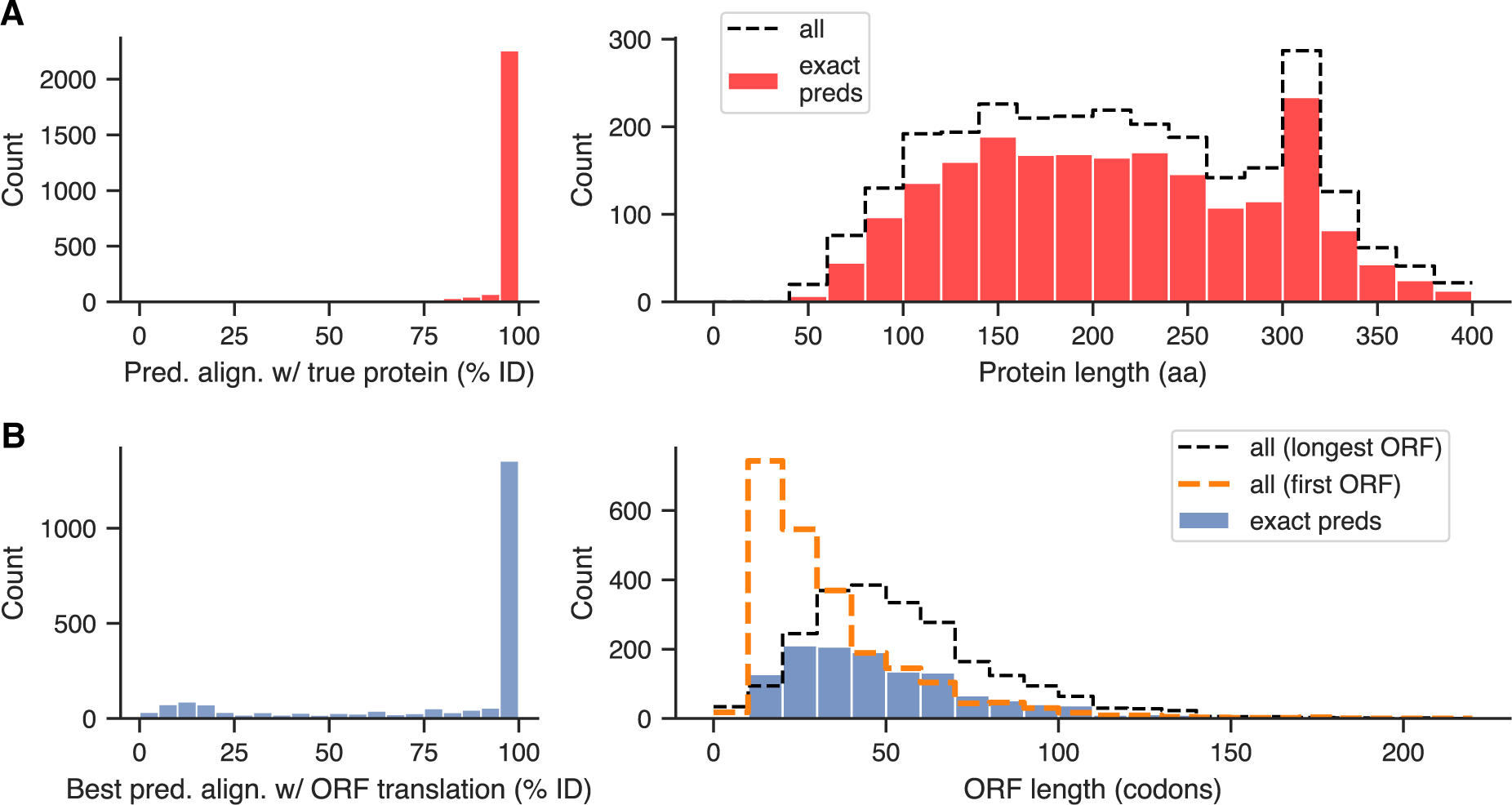
Analysis of translation products predicted by best bioseq2seq replicate. (A) Global alignment identity between the top-beam protein decoding predicted by bioseq2seq for true positive mRNAs and the ground truth protein (left), and length distribution of perfect translations (right). Black dashed line indicates the complete distribution of protein lengths. (B) Highest global identity found from all-by-all alignment of the three-frame translation of a lncRNA with its lower-beam *(PC)*+ peptide predictions from bioseq2seq (left) and length distribution of perfectly translated sORFs (right). Black dashed line indicates the length distribution of hypothetical translations of the longest ORF found in each lncRNA and orange dashed line denotes the same for the most 5’ ORF.

**Figure 3.**
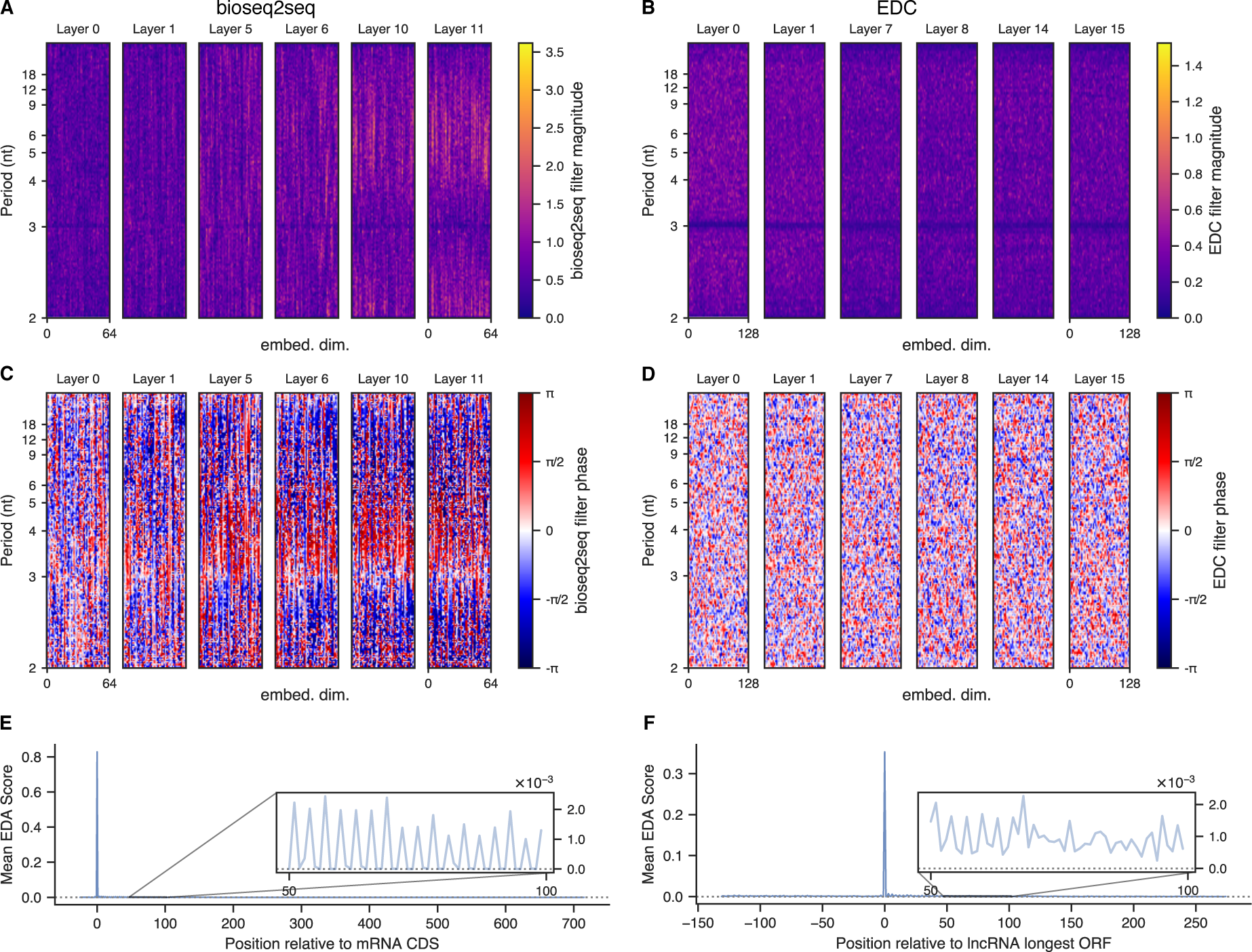
Frequency-domain content in model representations. LFNet filters from selected layers, with complex filter weights visualized in terms of magnitude (bioseq2seq in panel A, EDC in B) and phase (bioseq2seq in C, EDC in D). For each layer heatmap, the x-axis represents the hidden embedding dimension, and the y-axis refers to a discrete frequency bin, with annotations for the equivalent nucleotide periodicity. Both model types learned weights with a pronounced structure around 3-nt periodicity, visible mostly clearly in the phase for bioseq2seq and in the magnitude for EDC. (E) A nucleotide-resolution metagene consisting of average encoder-decoder attention scores from mRNAs aligned relative to their start codons. Attention distributions for this plot were taken from head 6 of the lower bioseq2seq decoder layer, which primarily attends to the start codon and places attention downstream of the start in a periodic fashion. (F) The equivalent plot for the same attention head applied to lncRNAs aligned relative to the start of the longest ORF, illustrating the loss of attention rhythmicity downstream of the leading spike.

Three-base periodicity is also apparent in our models’ encoder-decoder attention (EDA) distributions, which are probability weightings for encoder hidden embeddings in the context of each decoder layer. We aligned each encoder-decoder attention distribution for every transcript relative to its start codon and averaged to create nucleotide-resolution consensus attention metagenes. For lncRNAs, we investigated the longest ORF to define metagenes and to compare and contrast mRNAs and lncRNAs in the rest of this manuscript. We considered the two classes separately and discarded relative positions not present in at least 70% of the data, leaving relative position indices of (-25,+715) for mRNAs and (-131,+274) for lncRNAs. Depicted in Fig 3 are metagenes for a particular EDA head in the lower decoder layer of bioseq2seq that responds very differently to mRNAs (panel E) and lncRNAs (panel F), attending highly to the AUG/longest ORF in both classes but losing periodicity in lncRNAs. Sharp differences in attention such as this likely implement aspects of the model’s classification logic. We present more detailed analysis of EDA metagenes in Supplementary Fig S3.

### 2.4 Translation task improves reproducibility and biological plausibility of variant effect predictions

We evaluated all of our model replicates on every possible single-nucleotide variant of transcripts from a subset of our test data, consisting of 220 verified mRNAs and 220 verified lncRNAS. This technique, known as saturated *in silico mutagenesis* (ISM), is commonly used to computationally predict variant effects and can provide insight into input features that machine learning models recognize as important to their predictive task (Zhou and Troyanskaya 2015; Koo et al. 2021; Tareen et al. 2022). We calculated ISM using the function 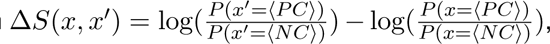, where *x* and *x^i^* are RNAs, with *x^i^* being a single-nucleotide variant of *x*. We calculated the Pearson correlation between the ISM scores predicted by two different replicates for a given transcript, making pairwise comparisons between all replicates. We also computed the cosine similarity between the character-level (A,G,C,U) vectors of mutation scores at each transcript position, using the median of this quantity as an transcript-level summary metric that does not consider the scaling of mutation scores at different positions. Both metrics were averaged across comparisons to produce a single value for each transcript, with the resulting distributions depicted in Fig 4-B. The inter-replicate agreement of bioseq2seq is much higher than that of EDC in terms of Pearson correlation (median of r= 0.813 vs. median of r= 0.560). The relaxed metric of median position-specific cosine similarity shows a minimal difference between bioseq2seq and EDC, which suggests that the gap in reproducibility between the model types is largely due to bioseq2seq’s more stable ranking of positional importance.

**Figure 4.**
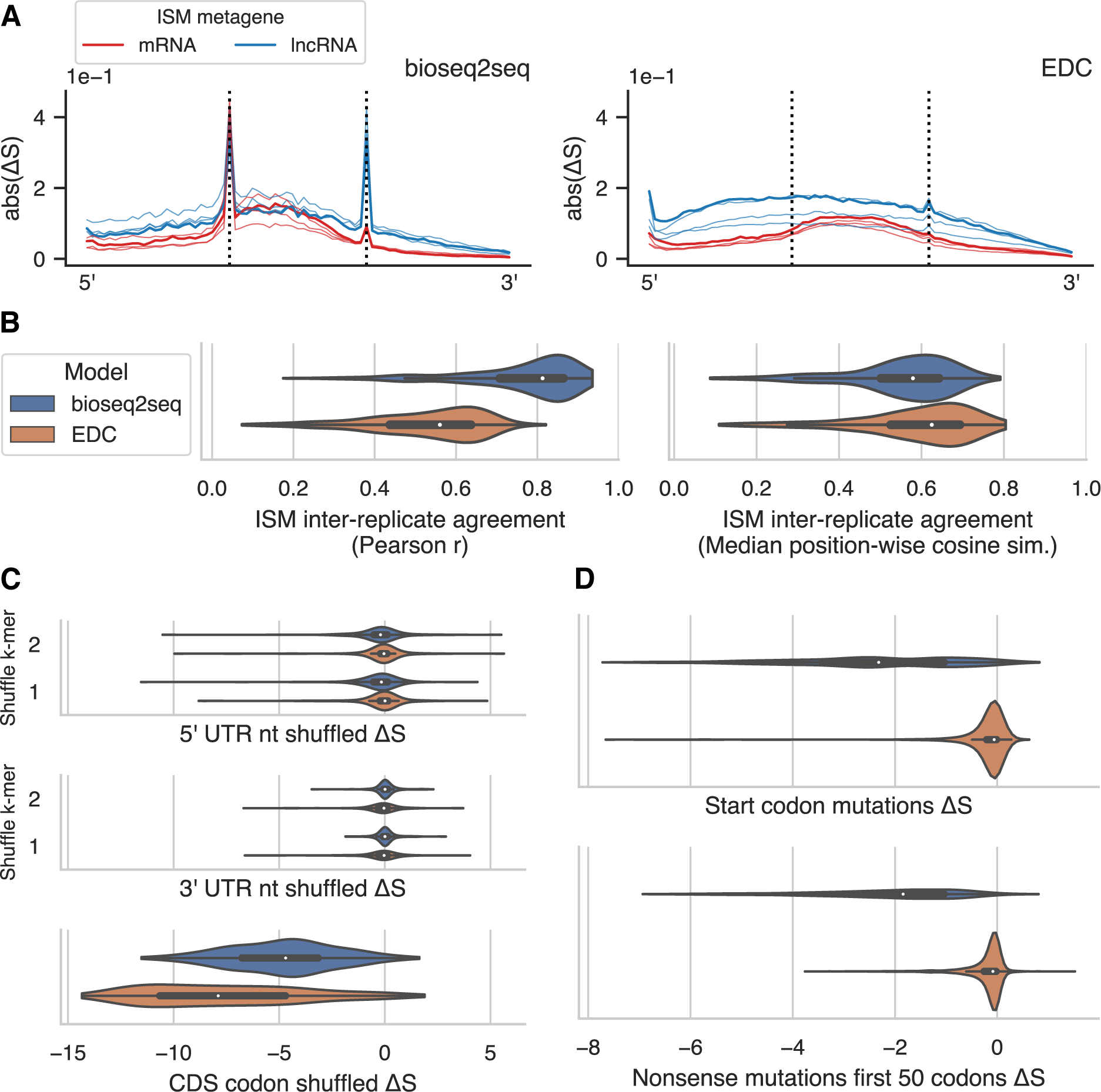
Predicted mutation effects by model type on a subset of testing data. (A) Metagene plots of saturated in silico mutagenesis (ISM) Δ*S* scores, i.e. the difference in log(*P* (*(PC)*)/*P* (*(NC)*)) between single-nucleotide variants and their wild-type sequence. The absolute value of Δ*S* was averaged within each of 25 positional bins and across all three possible mutations in each position, with mRNAs and lncRNAs depicted separately for both bioseq2seq (left) and EDC (right). Vertical dashed lines denote the first and last bin of the CDS for mRNAs and the longest ORF for lncRNAs. Metagenes from all four replicates are shown, with the best-performing model colored using the darkest hue. (B) Per-transcript average of Pearson correlation (left) and median position-specific cosine similarity (right) of ISM scores from pairwise comparison of model replicates. (C) Changes in score relative to wildtype for mRNAs shuffled within each functional region. UTRs were shuffled to preserve mononucleotide or dinucleotide frequencies. Codon shuffling excluded the start and stop codons to preserve CDS length. (D) Changes in score for mRNAs from nucleotide substitutions that knock out a start codon or introduce a stop codon within the first 50 codons of the CDS. Note: panels C and D follow the legend from panel B.

We next probed the ISM scores for changes disrupting essential mRNA features. We investigated the changes in score due to substantial sequence perturbations of each test mRNA by shuffling various functional regions. Specifically, we shuffled every 5’ UTR longer than 25 nt in the verified test set, using both an unrestricted shuffle and one preserving dinucleotide frequencies, and likewise for 3’ UTRs separately. We produced another set of variants by shuffling all codons besides the start and stop codon within CDS regions. This has the effect of preserving the original CDS length while likely disrupting 3-nt periodicity and leading to atypical orderings of nucleotides and amino acids. We calculated Δ*S* for each shuffled variant relative to its wild-type and found that UTR shuffling had minimal impact on on the predictions of either bioseq2seq or EDC (Fig 4-C). However, EDC is somewhat more reliant on the endogenous trinucleotide patterns of wildtype CDS regions than bioseq2seq, as indicated by the stronger negative Δ*S* after shuffling internal codons of CDS sequences. In contrast, mutations to the annotated start codon tended to produce large negative Δ*S* scores in bioseq2seq but not in EDC (Fig 4-D). Similarly, bioseq2seq responded negatively to mutations that introduced a stop within the first 50 codons. These observations suggest that while both models detect periodic sequence features, bioseq2seq has learned contextual sequence features, including start and stop codons, that more comprehensively align with our understanding of translation.

### 2.5 In silico mutagenesis reveals features predictive of coding potential

In light of the gap in biological robustness between our two model types, we investigated the response of bioseq2seq to sequence perturbations, using its best replicate to obtain ISM predictions for the remainder of the test set. We aggregated ISM scores for all synonymous point mutations inside of mRNA CDS regions into fine-grained metagenes for each amino acid, computing the mean Δ*S* along each of 25 positional bins. Selected amino acids are highlighted in Fig 5-A and all twenty are depicted in Supplementary Fig S4. As expected for a highly contextual model, there are large deviations away from the mean. On average however, the amino acids with only two codons all learn a preference for a single codon across the length of the whole transcript, with correspondingly negative scores for the opposite mutation. The amino acids with more than two-fold degeneracy are more complex to interpret but the sign for the mean mutation effect tends not to change with position. When considering all synonymous mutations, the model appears to have learned a preference for particular nucleotides in the codon positions. For example, most codons ending in C having a positive effect on Δ*S* on average, and most ending in T having a negative effect (Fig 5-B). Bioseq2seq’s estimates of synonymous mutation effects also captured some of the variation from an external measure of translation efficiency called tRNA Adaptation Index (tAI) (Reis et al. 2003). The mean Δ*S* for point mutations leading to synonymous changes show a moderate correlation (*r* = 0.394, *ρ* = 0.418) with the differences in tAI between the two codons, using codon values calculated from (Tuller et al. 2010). We used ISM scores as a feature explanation method by assigning each nucleotide within a transcript an importance score based on the magnitude of Δ*S* from the mutation in that position that most disrupts bioseq2seq classification towards the opposite class. For example, an endogenous *x_i_* within an mRNA was defined as contributing towards a true positive classification of the *(PC)* class to the extent that substituting any of the three alternate bases in position *i* produced a highly negative Δ*S*. One representative example mRNA and lncRNA are visualized in Fig 5-C and D, respectively, with raw ISM scores from positions of interest shown in a heatmap. The transcript sequences are overlaid above with their heights drawn proportionally to the importance setting for their true class – *↑ PC* for the mRNA and *↑ NC* for the lncRNA. The samples were chosen from among the five lncRNAs and mRNAs closest to the median value for inter-replicate agreement (see Fig 4-B). In the example mRNA, the start codon is a highly salient region, while the stop codon receives little importance. The ISM scores for the nucleotides surrounding the start codon imply a preference for G in position +1 relative to the start and A or C in position *−*2, consistent with the Kozak consensus sequence. The most important feature occurs in a region where many possible point mutations would introduce a stop codon, and we observed widespread avoidance of nonsense mutations early in the coding sequence. For the lncRNA, the TGA ending the longest ORF receives high importance according to *↑ NC*, but a different TGA upstream of the longest ORF is the highest overall.

**Figure 5.**
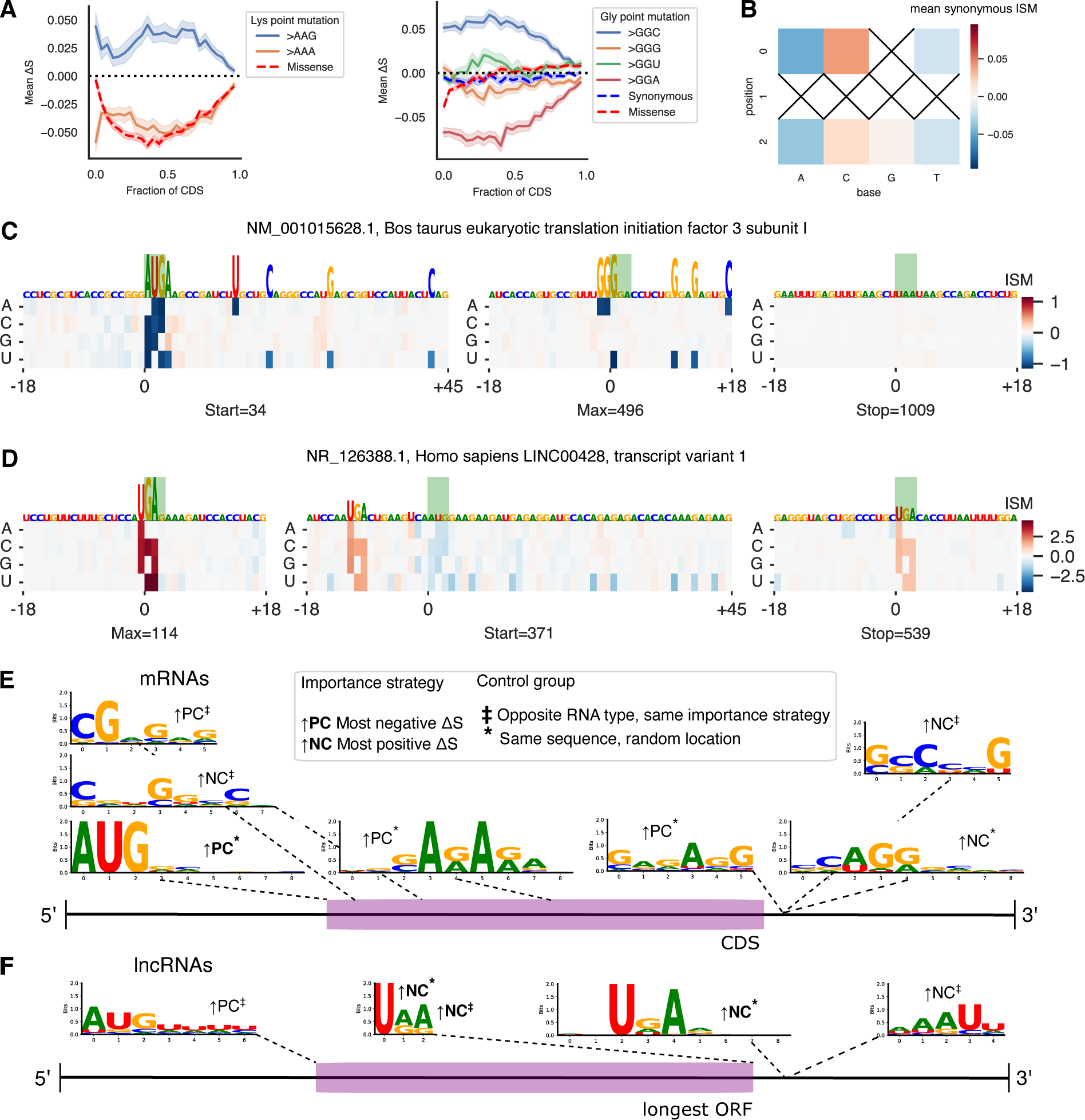
Detailed analysis of *in silico mutagenesis* (ISM) on the full test set. (A) Plots of ISM metagenes for selected amino acids lysine (left) and glycine (right). Mean Δ*S* is shown for 25 positional bins across mRNA CDS regions with mutations listed based on the resulting codon. The red line represents the average across all missense/nonsynonymous mutations. For amino acids with more than two codons, the blue dashed line depicts the average synonymous mutation for comparison. (B) Mean ISM for synonymous point mutations by codon position and nucleotide. X’s denote substitutions which do not exist as synonymous changes. (C) An example protein-coding transcript with NCBI accession NM_001015628.1. Signed ISM scores for the transcript are depicted as a heatmap and the RNA sequence is portrayed with characters scaled according to the *↑ PC* importance strategy, i.e. regions with highly negative ISM weights depicted in dark blue. The subregions shown are windows around the start codon, the position of maximum importance, and the stop codon, respectively. (D) Same as panel B with an example long noncoding RNA with NCBI accession NR_126388.1. The endogenous sequence is scaled according to *↑ NC*, or highly positive ISM values drawn in dark red. (E) mRNA motifs discovered in our test set with STREME using ISM importance values from bioseq2seq to determine sequence regions in which to search for enriched signals. Annotations denote the importance and control strategy for each trial, with boldfaced annotations signifying that importance values were not masked and ordinary typeface indicating that feature importance at start and stop codons and nonsense mutations were excluded. Motifs are positioned near the regions in which they were enriched. (F) Same as panel D showing discovered lncRNA motifs.

To systematically extract general patterns that bioseq2seq recognizes as predictive of coding potential, we performed de novo discovery of motifs frequently found in transcript subsequences with high ISM importance. First, we identified the most important nucleotide with respect to both *↑ PC* and *↑ NC* from each functional region (5’ UTR, CDS, 3’ UTR) of each test-set mRNA and likewise for the regions demarcated by the longest ORF of a lncRNA. We extracted 21-nt windows centered around each such important site to form a primary sequence database for the differential motif discovery tool STREME (Bailey 2021). A control set for STREME was constructed either using (1) random positions from the same transcript and region as the primary sequences but not overlapping them (2) the most important positions using the same importance setting as in the primary sequence but from the opposite RNA class. These controls necessitate different interpretations of the discovered motifs, with strategy 1 intended to establish whether bioseq2seq places importance on consistent features of a transcript, and strategy 2 intended to uncover differences in how bioseq2seq treats roughly comparable regions of coding and noncoding transcripts. We also ran motif discovery using a purely random strategy – e.g. with randomly chosen subsequences of a 5’ UTR as primary and random upstream regions of a lncRNA as control. We present only strategy 2 motifs that do no match a motif from the purely random trials according to TOMTOM (Gupta et al. 2007), as these experiments were specifically guided by ISM importance.

We ran every combination of primary sequence region, control method, and importance setting as its own STREME experiment and discovered four significant motifs between mRNAs and lncRNAs. Finally, we ran a second set of experiments in the same manner except with importance for endogenous start and stop codons and counterfactual missense mutations masked out in order to reveal important signals beyond the most prominent set found in the first run. This yielded an additional seven motifs, and both sets are shown in Fig 5-E for mRNAs and F for lncRNAs, with boldface annotations for the unmasked motifs. The experiments with random controls largely confirm the observations we made in our example transcripts, with a start codon/partial Kozak motif found in the beginning of mRNA CDS regions and several stop codon motifs prominent throughout lncRNAs. Beyond this, repeated GA patterns appear enriched in regions that push mRNAs towards a true positive classification and both control strategies uncover motifs that push mRNAs towards a false negative. Similarly, As and Us downstream of AUGs influence bioseq2seq towards a false positive prediction on lncRNAs, but such a motif receives comparatively little weight in the model’s assessment of bona fide coding transcripts. Additional details including positional and frame biases and enrichment, can be found in Supplementary Tables S4 and S5. We note a potential match with the binding site motif for an RNA-binding protein *ACO1* from (Ray et al. 2013), listed as motif #1 in Supplementary Table S5.

### 2.6 Approximation quality of gradient-based mutagenesis depends on model complexity

Saturated ISM is costly to apply to a large amount of sequences because it requires 3*L* model evaluations, where *L* is the transcript length. We explored the feasibility of approximating ISM using neural network input gradients, which are efficiently computable in parallel via automatic differentiation. Building from the Integrated Gradients (IG) method, we developed a novel proxy for ISM called Mutation-Directed Integrated Gradients (MDIG). MDIG involves numerically integrating input-output gradients along the linear interpolation path between a sequence of interest and a sequence of the same length consisting of all the same type of nucleotide, e.g. all guanines. A parameter *β ∈* (0, 1] limits how far to travel towards the poly(b) baseline embedding during integration. (See Methods). As a favorable value for *β* is not obvious from first principles, we tuned this parameter on a subset of our validation set consisting of 206 verified mRNAs and 206 lncRNAs, applying the same criteria from the previous section. To benchmark attribution stability across stochastic training, we measured the inter-replicate agreement of each mutation approximation method using Pearson correlation as in the previous section. We then computed the per-transcript Pearson correlation of scores from different settings of MDIG-*β* with the ISM scores from the same replicate. This metric indicates MDIG’s capacity to approximate the input-output behavior of a given deep learning model, which ISM accomplishes directly but at substantially greater computational cost. For reference with other gradient-based perturbations, we perform the same analyses using a first-order Taylor approximation of ISM scores and IG with a uniform [0.25, 0.25, 0.25, 0.25] baseline. Results on the validation set according to these evaluation metrics are summarized with their median value in (Fig 6-A), and full violin plots in Supplementary Fig S5. On the basis of these results, MDIG-0.5 was selected as the best approximation method for bioseq2seq and MDIG-0.1 for EDC. This illustrates that the MDIG method can predict the effect of input perturbations better than the basic Taylor approximation.

**Figure 6.**
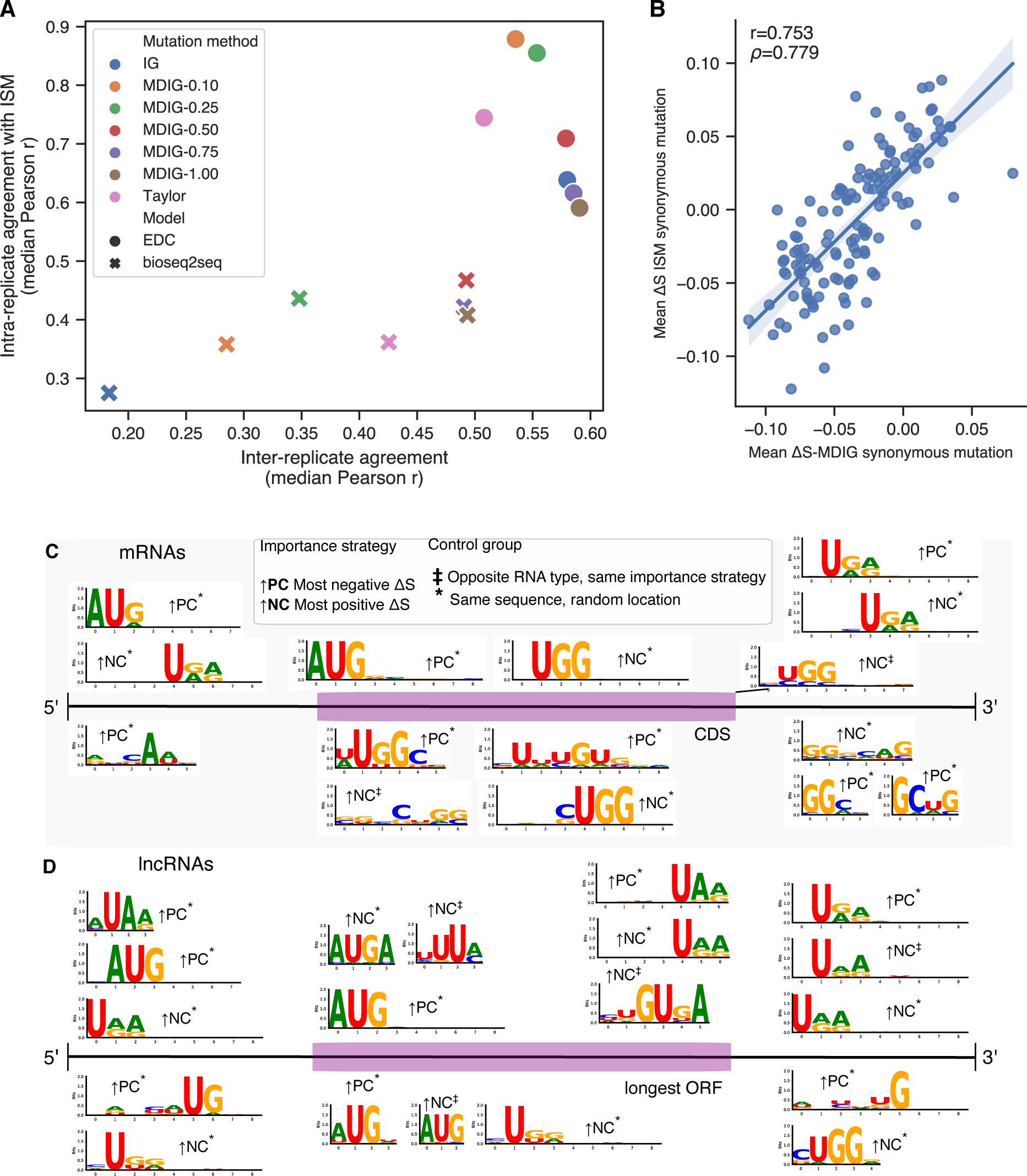
Gradient-based approximation performance. (A) Summary results from tuning of *β* hyperparameter for MDIG alongside baseline methods. Inter-replicate agreement is shown on the x-axis and correlation with ISM on the y-axis, using the median across transcripts as a point estimate for both metrics. (B) Scatter plot of Δ*S* for all possible synonymous point mutations, i.e. every wildtype>variant pair differing at one position, from MDIG on the training set (x-axis) versus the same for ISM on the test set. (C) mRNA motifs discovered in our training set with STREME using MDIG importance values from bioseq2seq to determine sequence regions in which to search for enriched signals. Results from unmasked importance are shown above the transcript diagram and those from the masked trials are shown below. (D) lncRNA motifs discovered in the training set using MDIG importance values from bioseq2seq, depicted in the same manner as panel C.

On the whole, we observed a large gap in approximation quality between the model types, with the best method for bioseq2seq lagging substantially behind the worst for EDC. To investigate the implications of MDIG’s reduced performance on bioseq2seq, we used the test data and method from the previous section to compute bin-based metagenes from the best MDIG versions and observed that this averaged representation closely captures the same general trends as expected from ISM (Supplementary Fig S6). Across the bioseq2seq replicates, MDIG metagenes have an average correlation of *r* = 0.897 for mRNAs and *r* = 0.944 for lncRNAs with their ISM equivalent, in comparison to *r* = 0.999 and *r* = 0.997, respectively for EDC. For a more detailed evaluation on bioseq2seq we approximated Δ*S* for every synonymous point mutation using MDIG on the test set and compared it with the true Δ*S* scores from ISM in the form of a scatterplot in Fig 6-B. The high correlation between metagenes and codon scores for ISM and MDIG indicates that despite its reduced transcript-level accuracy in predicting bioseq2seq mutation effects, MDIG largely captures the same class-level features as ISM when averaged across examples.

To take advantage of MDIG’s improved efficiency relative to ISM for improving the statistical power of motif discovery on our most biologically robust model, we applied MDIG-0.5 on bioseq2seq for the full training set. This consists of *∼*52k examples, balanced between the two RNA classes, and took about two days of GPU-time, much faster than our extrapolated estimate of more than a month for ISM (See Supplementary Table 2). We used the resulting MDIG mutation effect estimates as drop-in replacement for ISM importance values in our motif discovery pipeline, with the results presented in Fig 6-C for mRNAs and D for lncRNAs. Motifs from masked trials are placed below the transcript diagrams and those from unmasked trials are above. In comparison to the ISM motifs discussed previously, the MDIG motifs better underscore that bioseq2seq places importance on start and stop codons in regions besides the CDS. Start codons are predicted by MDIG to increase bioseq2seq coding probability in both mRNA 5’ UTRs and lncRNA upstream regions, while stop codons push the classification towards noncoding in the 5’ regions. Notably, the UTR motifs typically lack a bias towards a particular frame of the transcript, while most ORF features have a consistent frame bias. This is supportive of the idea that such elements outside the ORF are flagged in part to determine the frame. A number of interesting mRNA motifs emerge from masking, including multiple strong UGG motifs in a variety of sequence contexts and positions. The masked lncRNA motifs closely resemble those from the unmasked strategy, implying that the masked maximums are nucleotides adjacent to the start and stop codons. This comparative lack of diversity could mean that bioseq2seq largely defines lncRNAs as a class in terms of a lower quality or incorrect context of protein-coding features rather than distinctly ‘noncoding’ features. It also likely implies that MDIG is most adept at estimating the strongest mutation effects for bioseq2seq, with diminished reliability for less influential signals. One the whole, aggregating over instance-level MDIG scores to drive motif discovery appears to emphasize broad global features on which both MDIG and ISM both place high importance, while revealing additional signals beyond those identifiable with smaller-scale ISM experiments alone. As for ISM, the MDIG motifs are shown in greater detail in Supplementary Tables S6 and S7. We note a potential motif match to a binding site for an RNA-binding protein *SAMD4A* from (Ray et al. 2013) discovered in mRNA 3’ UTRs as ‘motif 1’ in Supplementary Table S7 and alongside possible ISM matches in Supplementary Fig S7.

## 3 Discussion

The genetic code makes it straightforward to predict protein sequences given an mRNA sequence, but our results suggest that requiring a neural network to learn the translation task improves its ability to identify protein-coding RNAs. We hypothesize that translation acts a regularization strategy by requiring the model to preserve precise positional information in a way that improves its contextual representations. Our findings are consistent with a related observation from RNAsamba, which performed worse when a network branch processing the longest ORF sequence was ablated (Camargo et al. 2020). Bioseq2seq differs from RNAsamba in that the translated protein sequence is an output rather than an input to the network. To our knowledge, bioseq2seq is the first attempt to use machine learning to output the encoded protein for an input RNA by explicitly learning the sequence mapping underlying biological translation. It accomplishes this from sequence alone, without introducing prior knowledge about the genetic code. Our models remain competitive with the best prior approaches without engineered sequence features, with bioseq2seq achieving on-par accuracy (less than 1% difference) and a higher recall.

The translation task also appears to significantly improve the quality of the nucleotide-level features identified by our models as predictive of protein-coding potential. The correlation of ISM mutation effects across multiple replicates is considerably higher for bioseq2seq than for EDC. Inter-replicate agreement quantifies the low epistemic uncertainty of mutation effect predictions made by an ensemble of bioseq2seq models. In the absence of experimentally characterized mutation effects, this suggests a robustness in the learned biological rules that can inform the plausibility of insights derived from feature interpretation. Besides this improvement in feature consistency, we found that the translation task confers an additional context-awareness to the model in a way that matches biological intuition. Even though simple features like ORF length are obvious correlates of ribosomal translation activity in the cell, the training process does not automatically impart this mechanistic insight into a neural network. We observed that EDC did not respond strongly towards mutations to either start codons or premature stop codons, suggesting such elements play a minimal role in its classification logic despite its relatively high best-case performance of 0.932 F1. Similarly, although mRNN recognizes start codons, it responded primarily to certain codons found 100-200 nt downstream of the start codon, rather than waiting for the stop codon (Hill et al. 2018). Bioseq2seq, however, responds negatively to start codon mutations, stop codon mutations, and nonsense mutations, suggesting that its decision-making is strongly influenced by its learned ORF features. Bioseq2seq’s faithful modeling of ORF features and mRNA periodicity improves the chances that it also makes biologically relevant effect predictions with respect to synonymous mutations and motif discovery, which require greater detail. We believe that the translation task steers the network toward more robust and meaningful representations that align with biological knowledge and show relative stability across replicates. In our view, these properties are vital prerequisites to enable a broader reliance on machine learning feature interpretation as a tool for scientific discovery.

Our treatment of gradient-based attributions is a contribution to the ongoing debate in the machine learning literature about the trustworthiness of such methods as neural network explanations. We benchmark gradient-based mutation effect predictions in the biological sequence domain against *in silico mutagenesis*, which is the concrete model response to meaningful sequence perturbations. Strikingly, the translation task appears to adversely affect the quality of gradient approximations, with all methods achieving relatively poor correlation with ISM for bioseq2seq but acceptable approximation quality in EDC. At a minimum, our results suggest that users of gradient-based feature explanations for genomics should follow a protocol similar to ours to validate gradient-based mutation effect predictions against more expensive but direct input perturbations. It might suggest that for some problems it is better to restrict architecture choices to convolutional neural networks, for which speedups of ISM exist (Schreiber et al. 2022). More fundamentally, there could be a practical trade-off between model complexity and accurate gradient approximation such that reduced fidelity of fast model perturbations is a price to pay for the superior classification performance and biologically plausible feature importance values that we observed in bioseq2seq.

We also introduce MDIG as a novel heuristic approximation for ISM, which we demonstrate can improve over Taylor approximation at a constant increase in computational complexity. MDIG is largely based on IG, but uses a more realistic mutation-specific baseline, and only integrates part of the way to the baseline value, staying closer to the original sequence. Despite the limited capacity of MDIG to estimate bioseq2seq mutation effects at the local, i.e. transcript level, we show its utility for identifying the most impactful sequence features at the global, i.e. class-wide level. This is supportive of recent work finding that the usefulness of approximate feature attributions can be improved by ensembling across alternative models (Gyawali et al. 2022). The similarity of important motifs and metagene representations derived from MDIG to their ISM analogues indicates that in aggregate MDIG retains interpretive value even where it does not faithfully model every individual mutation effect. Subject to appropriate validation, MDIG could be used where large-scale ISM experiments are infeasible or as a first-pass method to flag interesting sequences for more detailed review.

Interpreting bioseq2seq using ISM and MDIG revealed putative signals of regulatory information, which emerged purely from the learning process without prior specification. From a certain point of view, learning the sequence features that distinguish translated mRNAs from lncRNAs with untranslated ORFs would be informative for promoting ribosomal engagement and would promote translation.

We therefore expect that sequence features predicted to increase coding potential will correlate with codon bias. Common methods for assessing codon usage bias, such as Codon Adaptation Index, predict coding sequences according the relative skew of synonymous codons for a particular acid towards the codons most common in highly expressed genes. Bioseq2seq learned strong preferences within synonymous groups, as evidenced by consistently high mean value of Δ*S* across the entire transcript for specific codons. Codon preferences were noticeably grouped by the nucleotide in the third codon position, with substitutions towards nearly all codons ending in C having a positive mean effect, while nearly all ending in T/U have a negative effect. The existence of codon preference trends along the length of the transcript could reflect the fact that synonymous codon usage is known to be biased positionally, including towards rare codon clusters (Chaney et al. 2017). Replacing codons with those preferred by bioseq2seq in the average case could perform a similar function to optimizing based on CAI, but bioseq2seq learns mutation effects in the context of a codon’s transcript position and sequence neighborhood. Our mutation effect predictions are therefore a much richer source of information, and future work could test via experiment whether these preferred mutations impact translational efficiency and have potential to guide mRNA sequence optimization. The discovered motifs also reflect sensible biological intuitions, with the MDIG motifs in particular emphasizing upstream AUGs as increasing coding potential and stop codon trinucleotides as decreasing coding potential. This is consistent with evidence that upstream ORFs act to suppress the translation of the main ORF (Johnstone et al. 2016). Our motifs have a number of possible matches to RNA-binding proteins (RBP), which play essential roles in regulating transcript stability and translational activity. A potential match to the binding motifs for *SAMD4A*, a human RBP from the CIS-BP-RNA database (Ray et al. 2013) involved in the regulation of mRNA translation, was highlighted within regions of mRNA 3’UTRs which increase coding probability according to MDIG, consistent with the model treating this binding site as a valuable marker of coding potential. Several mRNA motifs reflect the Kozak sequence, and we find a contrasting pattern downstream of lncRNA AUGs with downstream Us and As which locally improves coding potential but is ultimately depleted in true protein coding sequences. The UGG trinucleotide recurs across several MDIG motifs in a variety of sequence contexts and positions. This could be explained in a number of ways: UGG is the unique codon for tryptophan, the rarest amino acid (Barik 2020), and is also one mutation away from the stop codons UGA and UAG.

Our demonstration that bioseq2seq can recover potentially translated micropeptides is a proof-of-concept for using machine predictions to explore this cryptic space of the proteome. Though the recovery rate of putative micropeptides from lncPEP is low overall, any such capability is incidental to our training setup and bioseq2seq mildly outperforms RNAsamba on the available data. Crucially, bioseq2seq is not inherently limited to only translating the longest ORF, which could prove to be a modeling advantage for this application given that many micropeptides are known to be harbored in ORFs other than the longest in a transcript (Makarewich and Olson 2017). Increased availability of validated micropeptide annotations and improved procedures for autoregressive decoding – see (Yang et al. 2018) for an example – could help a future method based on bioseq2seq to achieve higher reliability.

We anticipate that the LFNet architecture will be of broad utility in biological sequence modeling tasks, with frequency-domain multiplication enabling larger context convolutions than in common convolutional architectures and lower computational complexity of *O*(*N* log *N*) in comparison to transformers. Our extension of GFNet from (Rao et al. 2021) bridges older signal processing approaches for gene discovery with the flexibility of deep models. We also note the complementarity of our method with (Tseng et al. 2020), which, instead of empolying the Fourier-transform as a token-mixing method, used it to enforce a smoothness prior for importances on biological sequence models. Other applications of LFNet could include biological sequence data with variable periodic signals, such as nucleosome positioning (Epps et al. 2011) and gene organization (Wright et al. 2007), as well as other periodic non-biological data such as music. We designed the LFNet architecture based on an intuition that it could effectively leverage 3-nt periodicity, but such periodic structure is not necessarily an inherent requirement – the GFNet model was originally intended for computer vision.

There are numerous possible follow-up directions based on this work. Future versions could scale to a larger and more phylogenetically diverse dataset beyond the eight mammalian transcriptomes used here, as well as to longer sequence lengths. In this work we have treated coding potential as a binary classification problem, but the methods presented are readily applicable to the more general problem of predicting translational efficiency as a regression problem. The periodicity inductive bias in particular is likely to transfer to this task – Ribo-Seq data is also characterized by a 3-nt periodicity of footprint density, and this has informed the development of many ribosome profiling data analysis tools (Calviello et al. 2016; Xu et al. 2018). The regression setting could also increase the prospects for discovering novel regulatory features, such as in the UTRs, which our model treated as less important than the CDS. A network trained to stratify transcripts according to a quantitative measure of protein expression would likely learn more fine-grained distinctions than one modeling a binary separation between mRNAs and lncRNAs. Finally, our results raise the possibility that general-purpose nucleic acid language models could benefit from joint training with protein foundation models in a similar translation-like setup.

## 4 Methods

### 4.1 Seq2seq architecture for translation

Our model follows the encoder-decoder sequence-to-sequence (seq2seq) framework common in machine translation of natural languages (Vaswani et al. 2017). We call the model bioseq2seq because it applies the seq2seq paradigm to biological translation — with nucleotides and amino acids rather than human languages as the vocabularies. The output of bioseq2seq is a classification token 〈*PC*〉 for protein coding and 〈*NC*〉 for noncoding – followed by the translated protein in the case of 〈*PC*〉 and nothing in the case of 〈*NC*〉. Note that the network is not provided the location of the CDS, so it must learn to identify valid ORFs and select between potential protein translations.

Training bioseq2seq in this way allows us to test the hypothesis that the translation task will require the model to learn precise representations of each nucleotide, which will in turn help to attribute model decisions to specific sequence patterns. As a comparison with bioseq2seq, we also trained a model for binary classification. This secondary model, which we denote as Encoder-Decoder Classifier (EDC), has an identical network design to bioseq2seq, but a different training data format, as it was trained to output only the classification token without the additional protein product for mRNAs^I^. We developed our models in PyTorch based on a fork of the OpenNMT-Py repository for machine translation (Klein et al. 2018).

### 4.2 Local Filter Network

We initially experimented with transformer neural networks (Vaswani et al. 2017) for both the encoders and decoders but failed to produce competitive models, as biological sequences incur excessive memory costs as model sizes and sequence lengths grow. In these experiments, we found that the transformer encoders for bioseq2seq learned self-attention heads which principally attended to a small number of relative positional offsets while calculating the input embeddings. Additionally, feature attributions showed evidence of a strong 3-nucleotide periodicity (See Supplementary Fig S8).

A variety of recent papers have introduced efficient architectures which aim to preserve the ability of transformers to globally mix information at lower computational cost. A number of these approaches have used the Fourier transform as a substitute for self-attention, because it is an efficient global operation computable in *O*(*N* log *N*) time via the fast Fourier transform (FFT) algorithm (Lee-Thorp et al. 2021; Guibas et al. 2021). One such example for computer vision is the Global Filter Network, which takes the FFT of image patches and applies a learnable frequency-domain filter via elementwise multiplication, before inverting the FFT to return the representation to the time domain.

As the 3-base periodicity property is localized to coding regions within transcripts, we propose a simple modification to Global Filter Networks by substituting the global FFT with the short-time Fourier transform (STFT). While GFNet operates on non-overlapping patches of the input, we follow common practices for STFT using a stride equal to half the window size and weighting with the Hann function. To emphasize that our modification applies time-frequency analysis to sequence representations, we refer to this layer as a Local Filter Network (LFNet). A learned weight matrix *W* is applied equally to each window of the STFT and then the modified frequency content is returned to the time domain via the inverse FFT. A residual term is added to the result to carry along the previous representation. Following (Guibas et al. 2021), we apply the soft-shrink function after the weight multiplication to promote sparsity in the LFNet weights. LFNet layers are only used in the encoder stack of our networks, while the decoder stack consists of transformer decoder layers. This is because PyTorch currently lacks an implementation of causal masking for FFT, as would be necessary to efficiently train an autoregressive model with only LFNet layers.

### 4.3 Dataset

We built training and evaluation data sets using available RefSeq transcript and protein sequences for eight mammalian species: human, gorilla, rhesus macaque, chimpanzee, orangutan, cattle, mouse, and rat from RefSeq release 200 (O’Leary et al. 2016). We collected all RNA sequences annotated as mRNA or lncRNA and excluded transcripts over 1200 nucleotides (nt) in, which reduces the available data to 63,272 transcripts. Next, we linked each mRNA with the protein translation identified by RefSeq and partitioned the data into 80/10/10 training/validation/testing splits. To maximize the diversity of the dataset, we included transcripts with predicted coding status (XR_ and XM_ prefixes in Ref-Seq), as well as the curated transcripts (NM_ and NR_). For the training set, we used a balanced split between mRNAs and lncRNAs, selecting the split to equalize the length distribution of the two classes as much as possible. Finally, we ran CD-HIT-EST-2D to exclude from the test set all transcripts that exceed 80 % similarity with any transcript in the training set (W Li and Godzik 2006). The resulting test set contains 2288 lncRNAs and 2703 mRNAs.

### 4.4 Hyperparameter tuning and training

We used dynamic batch sizes, so that RNA-protein training pairs were binned based on approximate length to reduce the amount of padding. The maximum number of input tokens per batch was set to 9000 for both model types, and eight steps of gradient accumulation was used to increase the effective batch size. All models were trained to minimize a log cross-entropy objective function computed from each amino acid character in the output.

The hyperparameters including number of encoder and decoder layers, model embedding dimension, learning rate schedule, and L1 sparsity parameter were tuned via the Bayesian Optimization Hyperband (BOHB) algorithm provided in the Ray Tune library (Liaw et al. 2018). Candidate models were trained in parallel on four Tesla M10 GPUs with 8 GB GPU RAM and 640 CUDA cores, with one GPU per model. To enable a fair comparison between the bioseq2seq and EDC training objectives, hyperparameter tuning was run for each separately over an identical hyperparameter space from an initial starting point used during LFNet development. We also trained replicates for EDC using the best hyperparameters for bioseq2seq and refer to the best EDC model as EDC-large and the EDC with equivalent hyperparameters to bioseq2seq as EDC-small. We then produced four replicates for each of bioseq2seq, EDC-large, and EDC-small. For further details on hyperparameter tuning and model training see the Supplementary Details.

### 4.5 Mutation effect prediction

Estimating the effects of sequence mutations can provide insight into the importance that the model assigns each input nucleotide. The gold standard for computationally scoring mutation effects, known as *in silico mutagenesis* (ISM), requires comparing the model predictions for all single-nucleotide variants with that of the original sequence (Zhou and Troyanskaya 2015). The computational expense of this procedure – 3*L* model evaluations for a transcript of length *L* – motivates us to explore the effectiveness of gradient-based approximations.

Below we refer to the network output function by *S*, and the output gradient with respect to its input as *∇_x_S*(*x*). In general, *S* can be any scalar output, and here we use *S* = *l_(PC)_ − l_(NC)_*, the difference in logits, i.e unnormalized log probabilities, for the RNA classification tokens in the first decoding position. We denote the two sequences being compared as *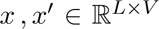* for one-hot encodings of categorical variables and *V* as the input vocabulary size.

### Taylor series approximation

The simplest ISM surrogate begins with a Taylor expansion of a differentiable function *F* around a point of interest *x^i^*.

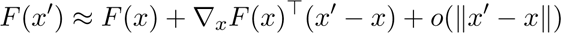

In this fashion, we can expand around *S* and discard all higher order terms for a first-order Taylor approximation of the difference in *S*.

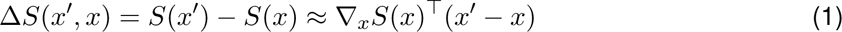

Since we confine our analysis to single-mutations, this simplifies to

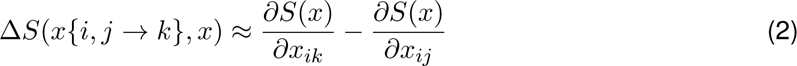

where *x{i, j −→ k}* is the result of mutating RNA *x* at position *i* from nucleotide *j* to *k*. Thus, all 3*L* values are computable from *∇_x_S*(*x*) in just one forward/backward pass of the network.

#### Mutation-Directed Integrated Gradients

The input gradient represents only an infinitesimal change in the input-output behavior of the network, rather than the effect of a full character substitution as in ISM. When a local approximation does not accurately describe the global function behavior, this is a well known limitation called gradient saturation (Shrikumar et al. 2019). As a more sophisticated proxy for ISM, we adapt a procedure called Integrated Gradients (IG), which was designed to reduce the the effect of gradient saturation and satisfies several desirable axioms for importance metrics (Sundararajan et al. 2017). IG uses a baseline input *x^i^* and computes an integral using input gradients for a differentiable function *F* along the linear path between *x^i^* and *x*.

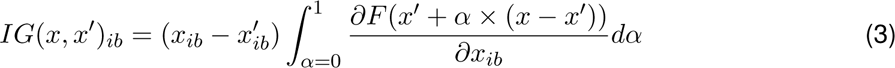

This equation relates to Taylor-approximation in that, given one hot encodings, 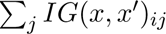 is equal to integrating the right hand side of Eq. 2 with *x* ranging over the interpolation path from Eq. 3. Based on this view, we propose a rough heuristic for estimating ISM using four evaluations of IG, which we dub Mutation Directed Integrated Gradients (MDIG).

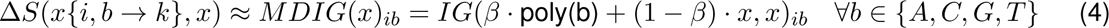

Here, poly(b) is a sequence of all nucleotide *b* of the same length as *x*, e.g. all guanines, and *β ∈* (0, 1] is a hyperparameter that balances distance of the baseline from *x*, which is needed to reduce gradient saturation, and distance from *x^i^*, a sequence largely unrelated to *x*. Note the order of arguments, which re-frames the baseline as the destination rather than the source.

To compare MDIG against a traditional usage of Integrated Gradients, we constructed an alternate baseline by placing the vector [0.25, 0.25, 0.25, 0.25] – a uniform probability mass function over the four bases – in all input positions. The mutation effect scores are then defined as Δ*S*(*x{i, b −→ k}, x*) *≈ IG*(*U* (*x*)*, x*)*_ik_ − IG*(*U* (*x*)*, x*)*_ib_* where *U* (*x*) represents the uniform baseline. In this way, the uniform IG approach requires just one evaluation of Eq 3 overall, while MDIG requires one evaluation per base *b*.

### 4.6 Evaluation metrics for gradient attributions

We compared *L ×* 3 vectors (sequence length *×* 3 possible mutations) of mutation effect predictions using the metrics

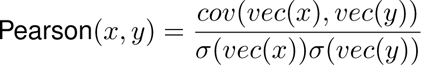

Median position-wise cosine similarity 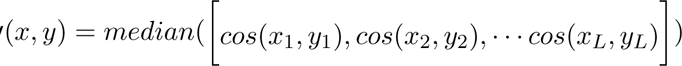

Where *cov*() is the covariance, *σ*() is the standard deviation,*vec*() is the vectorization operator, which flattens a matrix into a vector, 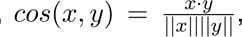, and *x_i_* refers to a row vector of matrix *x*. The inter-replicate agreement is

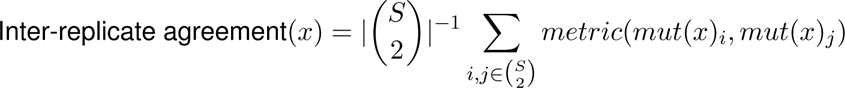

where 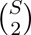 is the set of all possible subsets of cardinality 2 from the set of model replicates *S*, *mut*(*x*)*_i_* is the mutation effect prediction coming from replicate *i* for a given RNA *x*, and *metric* is one of Pearson r or median position-wise cosine similarity, as described above. The agreement with ISM is defined with intra-replicate comparisons.

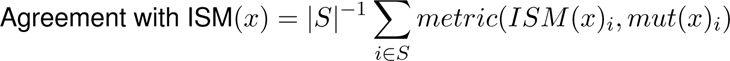

### 4.7 Motif discovery from mutation effect predictions

To uncover sequence elements salient to bioseq2seq predictions, we converted ISM scores into importance scores for the endogenous characters. In particular, we set the importance score of an endogenous base with respect to a given class as equal to the absolute value of Δ*S* for the strongest mutation in the direction of the *counterfactual* class, following (Kelley et al. 2016) which used the equivalent from regression models for visualizing importance. For example, an endogenous *x_i_* within an mRNA was defined as contributing towards a true positive classification of 〈*PC*〉 to the extent that substituting any of the three alternate bases in position *i* produces a highly negative Δ*S*, which pushes the prediction towards a false negative of 〈*NC*〉. We calculated importance using both classes on all transcripts. For instance, we looked for strong local contributions towards a prediction of 〈*PC*〉 within annotated lncRNAs.

For a given importance setting, we then extracted a window of 10 nt upstream and 10 nt down-stream around the position with the highest importance score for a total length of 21 nt. This process was run separately for mRNA 5’ and 3’ UTRs and CDS sequences, and similarly for lncRNAs using the longest ORF and its upstream and downstream regions. We used the STREME motif discovery tool to efficiently identify sequence motifs occurring frequently in these regions of interest (Bailey 2021). STREME estimates p-values for motifs, and after collecting all discovered sequence logos, we reported all that were significant at the 0.001 level after applying the Bonferroni correction for multiple testing.

## 5 Data Access

Code for running our trained models and replicating the experiments and figures in this paper is provided at https://github.com/josephvalencia/bioseq2seq and pretrained models and data at https://osf.io/xaeqg/.

## 7 Author Contributions

J.D.V and D.A.H conceived of the project. J.D.V conceived of the LFNet architecture. J.D.V performed all coding and analysis while supervised by D.A.H. J.D.V. wrote the manuscript, and D.A.H provided edits.

## Footnotes

I Although including a decoder is somewhat atypical when producing a single output classification, we do this to enable a direct comparison between the training tasks under a common architecture. The role of the decoder in the EDC setting is to calculate multi-headed attention distributions over the encoder hidden states, with the pre-pended ‘start-of-sentence’ token playing a similar role to the ‘[CLS]’ in encoder-only classification setups.

