## Supplementary Methods for "Improving deep models of protein-coding potential with a Fourier-transform architecture and machine translation task"

April 4, 2023

**1** School of Electrical Engineering and Computer Science, Oregon State University, Corvallis, OR , USA

**2** Department of Biochemistry and Biophysics, Oregon State University, Corvallis, OR, USA

#### A Architecture and training details

The short-time fast Fourier transform in LocalFilterNetwork was implemented with the torchaudio library (Yang et al. 2022) using the Spectrogram and InverseSpectrogram classes. We used pad_mode='constant' to pad the input with zeros where the LFNet window overhangs the sequence, which has the effect of upsampling the sequence within such windows to match the filter frequency resolution. We also padded batches to accommodate input examples with different numbers of tokens.

We implemented hyperparameter tuning with the Ray Tune PyTorch library (Liaw et al. 2018) and the Bayesian Optimization HyperBand (BOHB) method (Falkner et al. 2018). We chose BOHB because it allows models based on different hyperparameter settings to be evaluated simultaneously while terminating less promising trials early for a desirable degree of parallelism and efficiency. At the same time, BOHB uses Bayesian optimization to adaptively sample from the hyperparameter search space for improved model performance. One of the four initial settings was set to a configuration found to work well during development. The range of hyperparameters that we allowed to be randomly sampled is shown in Table 1 along with the optimal set of hyperparameters found for each model type. We used the learning-rate warmup schedule from (Vaswani et al. 2017), with the number of warmup steps as a hyperparameter.

| Range | Seed | bioseq2seq | EDC |
|---|---|---|---|
| Model dim = [32,64,128] | 64 | 64 | 128 |
| # Encoder layers = [1,2,4,8,12,16] | 12 | 12 | 16 |
| # Decoder layers = [1,2,4,8,12,16] | 12 | 2 | 16 |
| Dropout probability = [0.1,0.2,0.3,0.4,0.5] | 0.2 | 0.2 | 0.1 |
| Learning rate warmup steps = [2k,4k,6k,8k,10k] | 4000 | 2000 | 4000 |
| LFNet window size = [100,150,200,250,300,350,400] | 200 | 250 | 200 |
| L1 sparsity multiplier $\sim LogUniform$(1e-1,1.0) | 0.5 | 1.1e-2 | 4e-3 |

**Table 1.** Hyperparameter search space and optimal hyperparameters for bioseq2seq and EDC.

The LFNet filters learned different frequency-domain strategies in the two model types, with bioseq2seq weights having more extreme phase values, as seen in Fig 1. Phase values for the 3-nt band are shown as a kernel density plot with the remaining periods as a histogram, with bioseq2seq in panel A and EDC in panel B. Values close to $\pm\pi$ imply a partial cancellation effect when filtered representations are added to the residual term at the end of each LFNet layer. Bioseq2seq has learned to suppress periodic signals besides 3-nt periodicity while modulating the critical 3-nt signal to a lesser extent. EDC filters show less discrimination between 3-nt and the remainder in terms of phase activity.

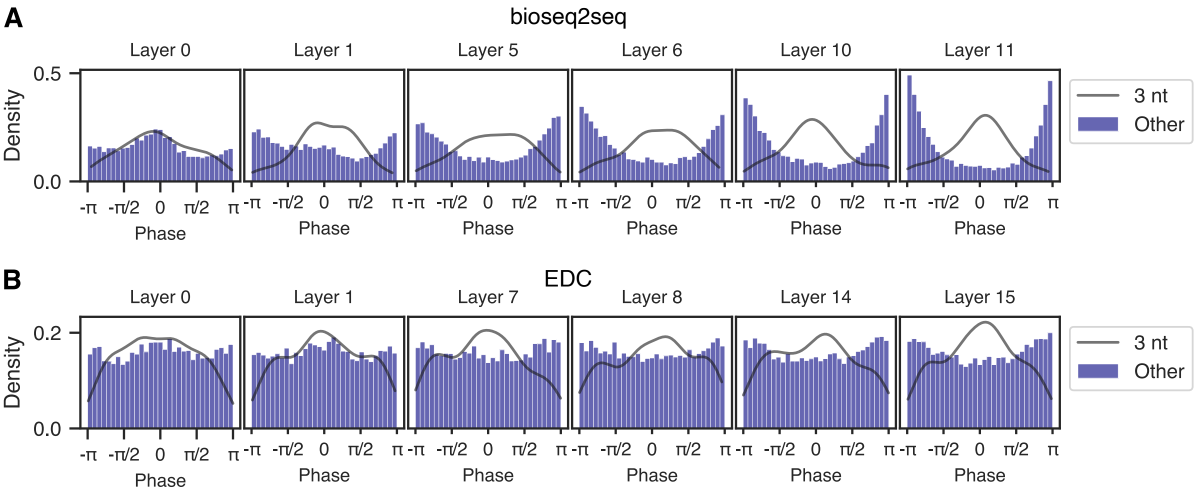


**Figure 1.** LFNet filter phase histograms. (A) Density of phase values for selected layers of bioseq2seq, with 3-nt as a kernel density plot denoted by the black line and the remaining frequencies as a purple histogram. (B) The same as panel A for EDC.

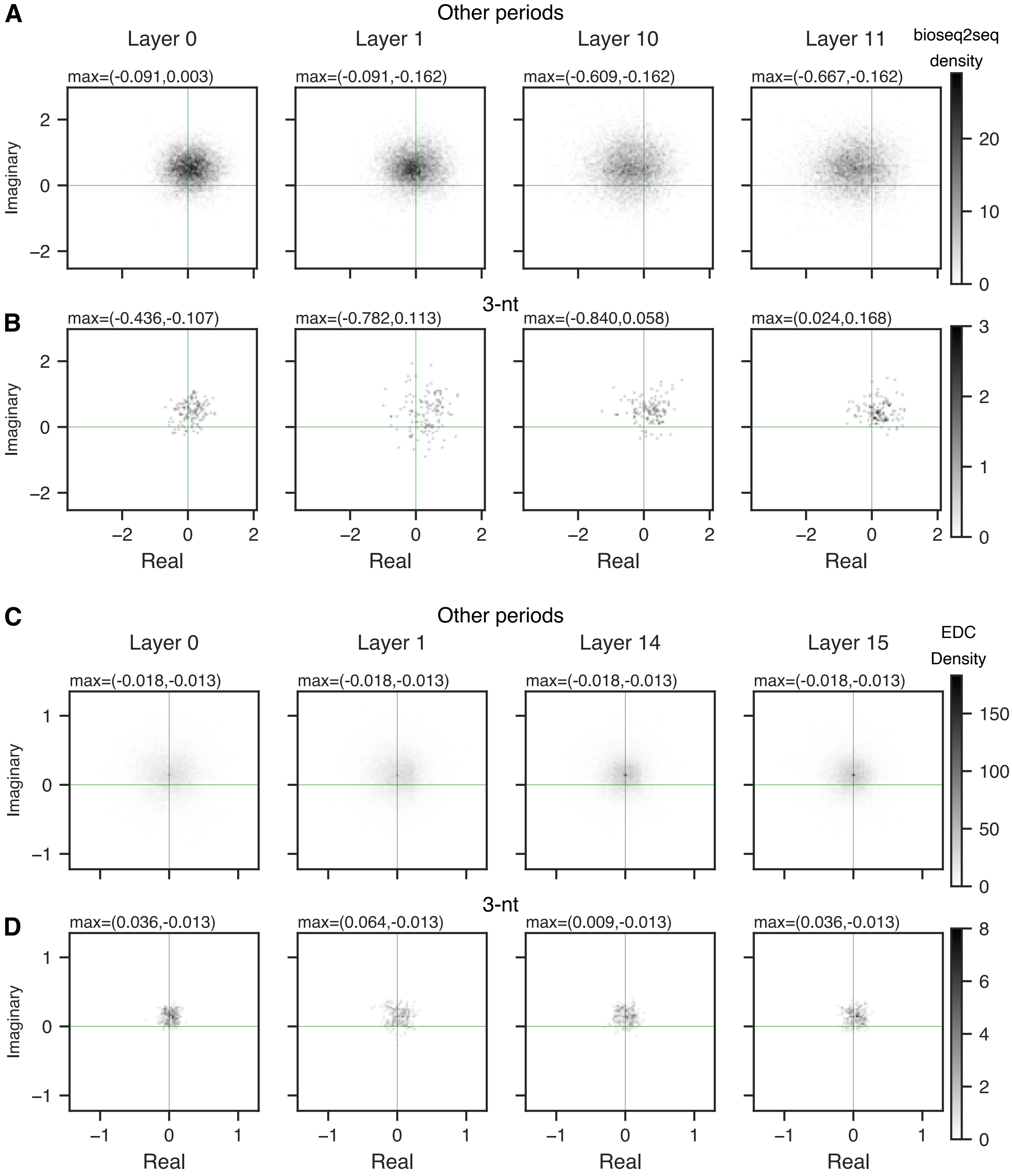


**Figure 2.** Analysis of real and imaginary components from LFNet Filters (A) Frequencies besides that for 3-nt periodicity in bioseq2seq. (B) Frequencies for 3-nt periodicity in bioseq2seq. (C) Frequencies besides 3-nt in EDC. (D) Frequencies for 3-nt periodicity in EDC.

#### B Interpreting encoder-decoder attention

We built nucleotide-resolution metagenes for encoder-decoder attention (EDA) by aligning the attention distributions of each transcript relative to its start codon or AUG of the longest ORF. We used Welch's method for spectral density estimation to decompose the attention metagenes into a density over a range of frequencies, plotting the resuling power spectrum decompositions (PSD) in Fig 3. Respectively, panels A-D contain PSDs for: bioseq2seq on mRNAs, EDC on mRNAs, bioseq2seq on lncRNAs, and EDC on lncRNAs. Bioseq2seq distinguishes the two classes more clearly, with one attention head in its upper decoder layer losing periodicity altogether for lncRNAs. EDC attention values lead to a somewhat noisier power spectrum in lncRNAs than in in mRNAs.

We also binned and averaged encoder-decoder attention (EDA) values within functional regions in the same fashion as previously described for ISM and MDIG. The resulting metagenes are conceptually distinct from those calculated at nucleotide-resolution because the averaging process clarifies positional trends by drowning out the 3-nt periodicity that dominates at nucleotide-resolution. Plots of bin-based metagenes for bioseq2seq are shown for every attention head, with heads from the lower decoder layer in panel E and those from the higher decoder in panel F. With the caveat that LFNet encoder embeddings integrate contextual information from many different input positions, we observed that two heads from Layer 0 (1 and 6) allocated almost all attention to the position of the start codon for mRNAs or the AUG of the longest ORF for lncRNAs. The metagenes in Layer 1 indicate a stronger differentiation between mRNAs and lncRNAs than in Layer 0. Several heads (1,2,4) distributed attention for mRNAs in a bellcurve-like shape centered close to the middle of the CDS, with no such pattern for lncRNAs. The metagenes for mRNAs and lncRNAs follow somewhat opposite trajectories in heads 0 and 5, with average attention decreasing along the length of the transcript for mRNAs and increasing for lncRNAs. This could mean that the first decoder layer marks basic RNA features while the second emphasizes higher-order features that separate coding and noncoding transcripts.

#### C Data subsampling

To explore feature consistency across model types and replicates while keeping computational costs manageable, we restricted several experiments to a subset of our testing data consisting of the tran-

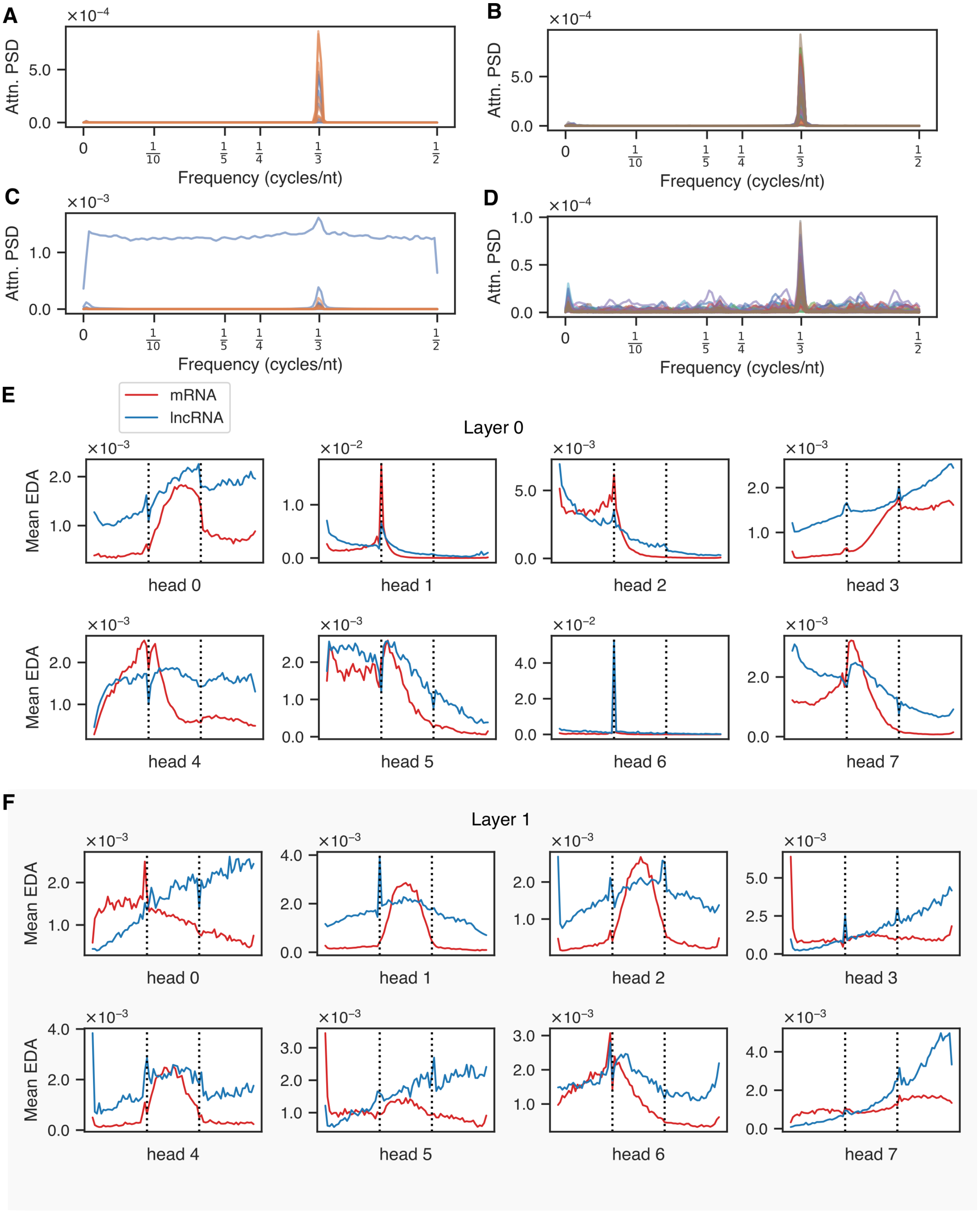


**Figure 3.** Analysis of encoder-decoder attention (EDA). (A) Power spectrum decomposition of EDA metagenes for mRNAs on bioseq2seq. Equivalent nucleotide positions relative to the start codon in mRNAs were aligned and corresponding attention scores from each attention head were averaged to form the nucleotide-resolution metagene. (B) The same as panel A using encoder-decoder attention from EDC (C). Power spectrum decomposition for lncRNAs aligned relative to the AUG of the longest ORF on bioseq2seq. (D) The same as panel C using encoder-decoder attention from EDC. (E) EDA from Layer 0 of bioseq2seq, the lower in the decoder stack, with attention values averaged within 25 positional bins for each functional region. First and last bins of the CDS or longest ORF marked with dotted vertical lines. (F) EDA metagenes from Layer 1, the higher decoder in bioseq2seq.

scripts with verified status in RefSeq. For the experiment described in section 2.6 of the main text, we took all 220 transcripts with ‘NM’ ids in the test set and sampled an equal amount of transcripts with ‘NR’ ids. This is referred to here and in the main text as the "verified test set". The verified validation set used to tune the $\beta$ value of MDIG in section 2.6 was created using the same criteria.

In several experiments we analyzed transcripts by functional region – CDS and UTRs for mRNAs or the longest ORF and regions upstream and downstream for lncRNAs. For the bin-based metagenes, we used 25 positional bins for each functional region, which led us to require that every functional region be at least 25 nt in length. For the full test set, 1675 mRNAs and 1977 lncRNAs met these criteria. For the verified test set these numbers were 152 mRNAs and 197 lncRNAs.

### D Gradient-based mutation effect prediction

For all mutagenesis experiments, we used the function $F = l_{\langle PC \rangle} - l_{\langle NC \rangle}$, where $l_c$ is the neural network activation before the final softmax layer for class $c$. These activations are also called the logits and express finer differences than the post-softmax probabilities. Our choice of $F$ ensures that evidence from both RNA classes contributes to gradient-based attributions.

All our gradient-based attributions are calculated at the one-hot level, but PyTorch sequence models, including ours, are commonly trained with an embedding layer based on a dictionary-like object. To introduce the one-hot representation into the computational graph at inference time, the weights matrix object from the torch.nn.Embedding layer is extracted and the input tensors are converted to a one-hot encoding. An explicit matrix multiplication between the one-hot encoding and the embedding weights preserves the behavior of the Embedding layer while enabling differentiation with respect to the one-hot vectors.

#### D.1 Mutation Directed Integrated Gradients

MDIG is intended to balance the competing goals of (1) accumulating global information correlated with point mutations and (2) maintaining distance from the uninformative baseline poly(b), while the linear IG interpolation permits parallelism in the sequence dimension. Integrated Gradients satisfies the property $\sum_i \sum_j IG_{ij}(x, x') = F(x) - F(x')$ (Sundararajan et al. 2017). Since we are seeking to relate IG scores to $3L$ single-nucleotide variants rather than the baseline $x'$ itself, this identity is of reduced importance

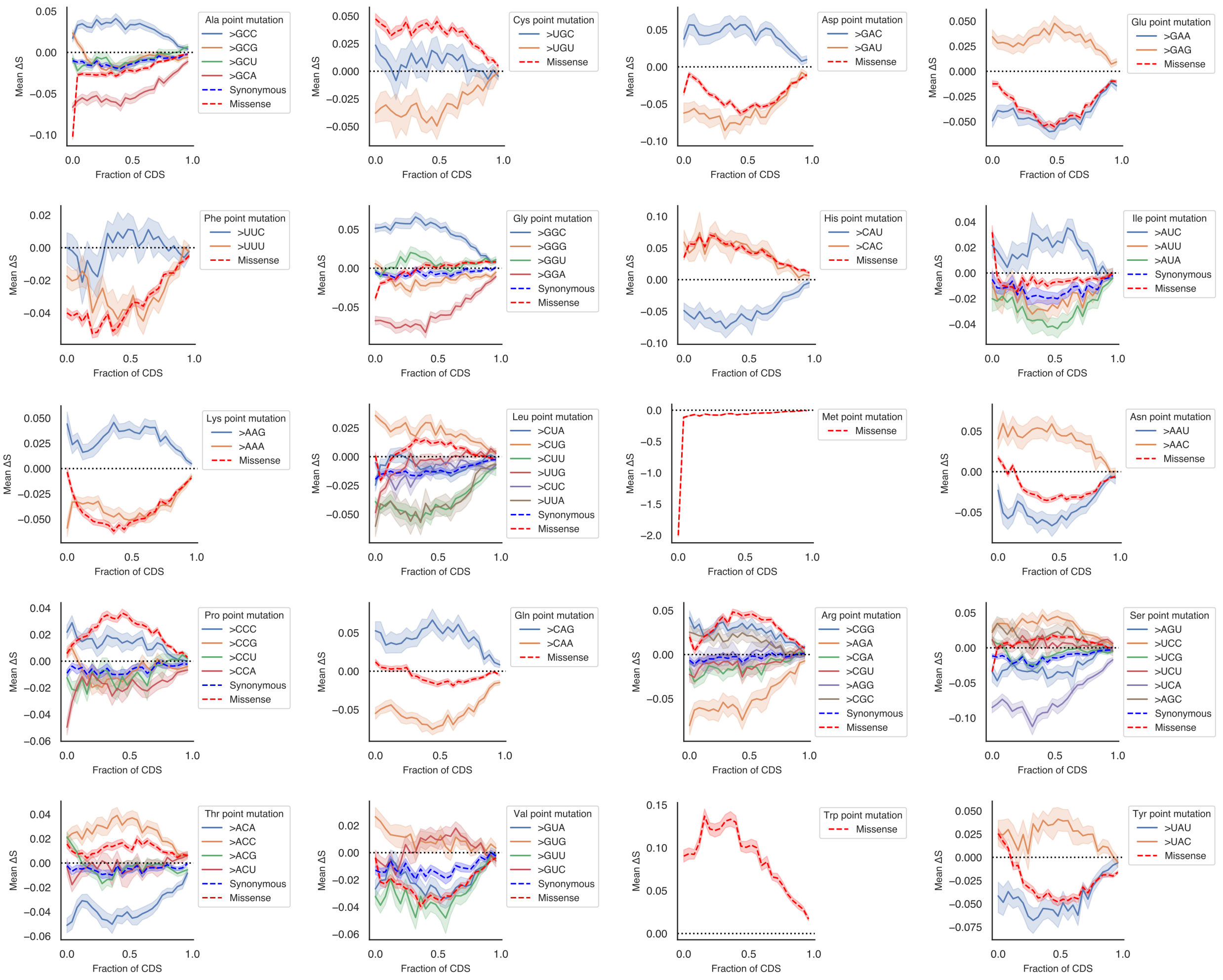


**Figure 4.** Plots of ISM metagenes for all twenty amino acids. Mean $\Delta S$ is shown for 25 positional bins across mRNA CDS regions. Mutations are listed based on the resulting codon. The red line represents the average across all missense/non-synonymous mutations. For amino acids with more than two codons, the blue dashed line depicts the average synonymous mutation for comparison.

compared to typical IG use cases. Still, we used this property to compute approximation error and inform our selection of 32 numerical integration steps for the experiments in the main text, which also provided for adequate convergence of IG values.

Fig 5 depicts the full distributions from evaluating each mutation effect algorithm according to its intra-replicate agreement with ISM (panel A) and inter-replicate agreement (panel B) on the verified validation set. A possible explanation for the values of $\beta$ that proved best during tuning MDIG is that at $\beta < \frac{1}{2}$, the interpolated embedding $Embed(\beta \cdot \text{poly(b)} + (1 - \beta) \cdot x)$ is row-wise closer to $Embed(x)$ rather than $Embed(\text{poly(b)})$ by Euclidean distance. In other words, the closest embedding of any discrete/non-interpolated sequence prior to this point is the original sequence, so the accumulated gradients maintain a connection to the original sequence, whereas at $\beta > \frac{1}{2}$, the embedding is closer to the poly(b) baseline. Proximity of embeddings by Euclidean distance does not ensure any particular relationship between their network outputs, but we found empirically that MDIG can correlate quite well with ISM for EDC and pick up the most important features for bioseq2seq.

Using MDIG-0.5 as the best setting for bioseq2seq and MDIG 0.1 for EDC, we built metagenes for all replicates on the verified test set. The general positional trends for mRNAs and lncRNAs closely mirror those from the equivalent ISM metagenes, as do the spikes in $|\Delta S|$ at the borders of the CDS/longest ORF (see Fig 6). We explored the implications of using MDIG-0.5 as a proxy for large-scale ISM experiments by applying it on bioseq2seq for the entire training set of 52,078 samples. For increased efficiency, we used just 8 integration steps per poly(b) baseline. Time was measured with GNU time, with the estimate for ISM extrapolated from its observed runtime on the test set. The projection of more than five GPU-weeks for ISM versus just over two GPU-days for MDIG demonstrates the possible efficiency gains from making this approximation.

| Inference mode | Per-sample model passes (fwd./back.) | GPU time (days:hrs:mins) |
|---|---|---|
| Prediction only | 1/0 | 00:00:38 |
| Taylor approx. | 1/1 | 00:01:56 |
| MDIG | 32/32 | 02:03:32 |
| ISM | $3L$/0 | 39:03:06** |

**Table 2.** Computational cost of evaluating bioseq2seq on the training set (52,078 RNAs, class balanced) with various inference/attribution methods. Time was measured using GNU Time and MDIG was run using eight numerical integration steps per baseline for a total of 32 forward-backward passes.** Extrapolated from full test set (≈90 GPU-hrs / 4491 RNAs )

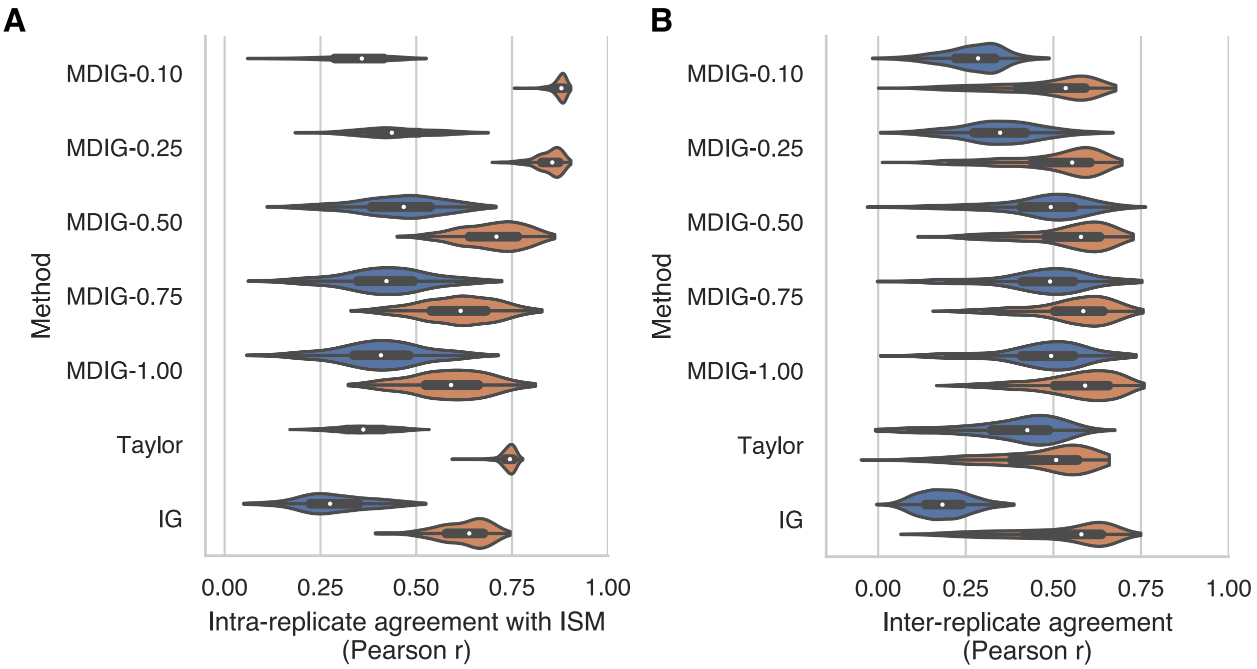


**Figure 5.** Evaluation metrics from tuning MDIG on the validation set, alongside Taylor approximation and Integrated Gradients as baseline attributions. (A) Agreement in terms of Pearson correlation between each mutation effect algorithm and *in silico mutagenesis* on the same replicate (B) Agreement in terms of Pearson correlation of each mutation effect algorithm with itself across replicates.

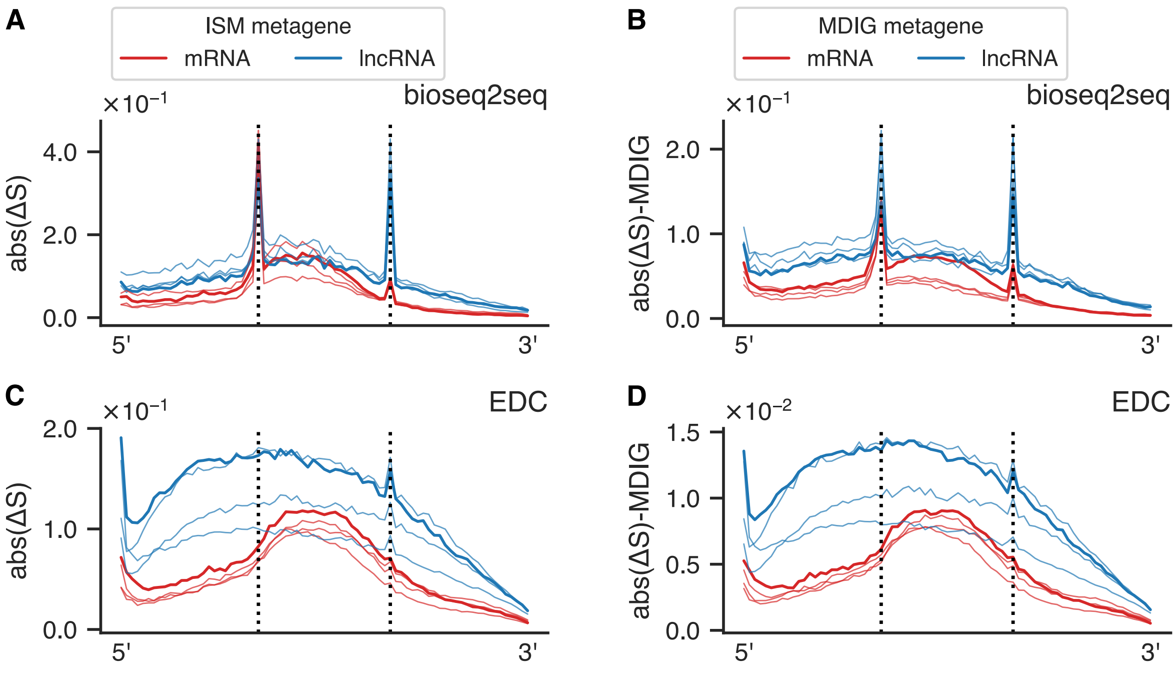


**Figure 6.** Extended results from best settings for MDIG. (A) Plot of bin-based metagene for bioseq2seq using ISM, reproduced from the main text. (B) Plot of bin-based metagene for bioseq2seq using MDIG-0.5. (C) Metagene from ISM on EDC. This is the same as the plot from the main text without y-axis scaling to match bioseq2seq. (D) Plot of bin-based metagene for EDC using MDIG-0.1.

#### E Motif comparison

We used TOMTOM (Gupta et al. 2007) to compare motifs in several portions of our analysis. The program was run with default parameters except with '-norc' added to compare motifs found in control strategy 2 matching ones from the purely random control strategy. Matches at this significance threshold were omitted from our analysis to emphasize those uniquely identifiable with our bioseq2seq importance scores. Masked motifs for both ISM and MDIG were run against human and mouse RBPs from (Ray et al. 2013) using more stringent parameters of '-norc -thresh 0.1 -min-overlap 5'. The resulting possible matches are shown in Fig 7.

#### F Motivation from Transformer encoder

Earlier in model development, we experimented with transformer neural networks (Vaswani et al. 2017) for both the encoder and decoder stacks based on the widespread success of the transformer in natural language processing. While this architecture successfully learned the genetic code, its classification accuracy lagged noticeably behind the state of the art methods such as RNAsamba. Whereas LocalFilterNet uses a short-time Fourier transform and frequency domain filters, transformers use self-attention to calculate an all-by-all similarity between input embeddings. This $O(N^2)$ complexity of time and especially memory contributed to our inability to train a competitive model with transformers. However, we observed interesting properties of the learned transformer representations that inspired our eventual design of LFNet, particularly a striking dependence on relative positional offsets in self-attention.

For a transcript of length $L$, a self-attention head produces an $L \times L$ dimensional matrix, with each row containing a probability distribution weighting the local influence of each nucleotide position on each other. For each nucleotide $i$, we collected its self-attention argmax, recording both its absolute position $j$ and relative positional offset $j - i$. We then determined the mode of these values for the transcript, $mode_{abs}$ or $mode_{rel}$. Finally, we recorded the support for the mode – the percentage of indexes which equal the modal index $n_{abs}$ or $n_{rel}$. To summarize the behavior of each self-attention head, we aggregated these transcript-level statistics over every RNA in the test set. Fig 8 depicts these results, with the cells in the heatmap colored with the average value of $n_{abs}$ or $n_{rel}$. Whenever the variance of $mode_{rel}$ over all RNAs was less than 1, we included $mode_{rel}$ as an annotation. A dark

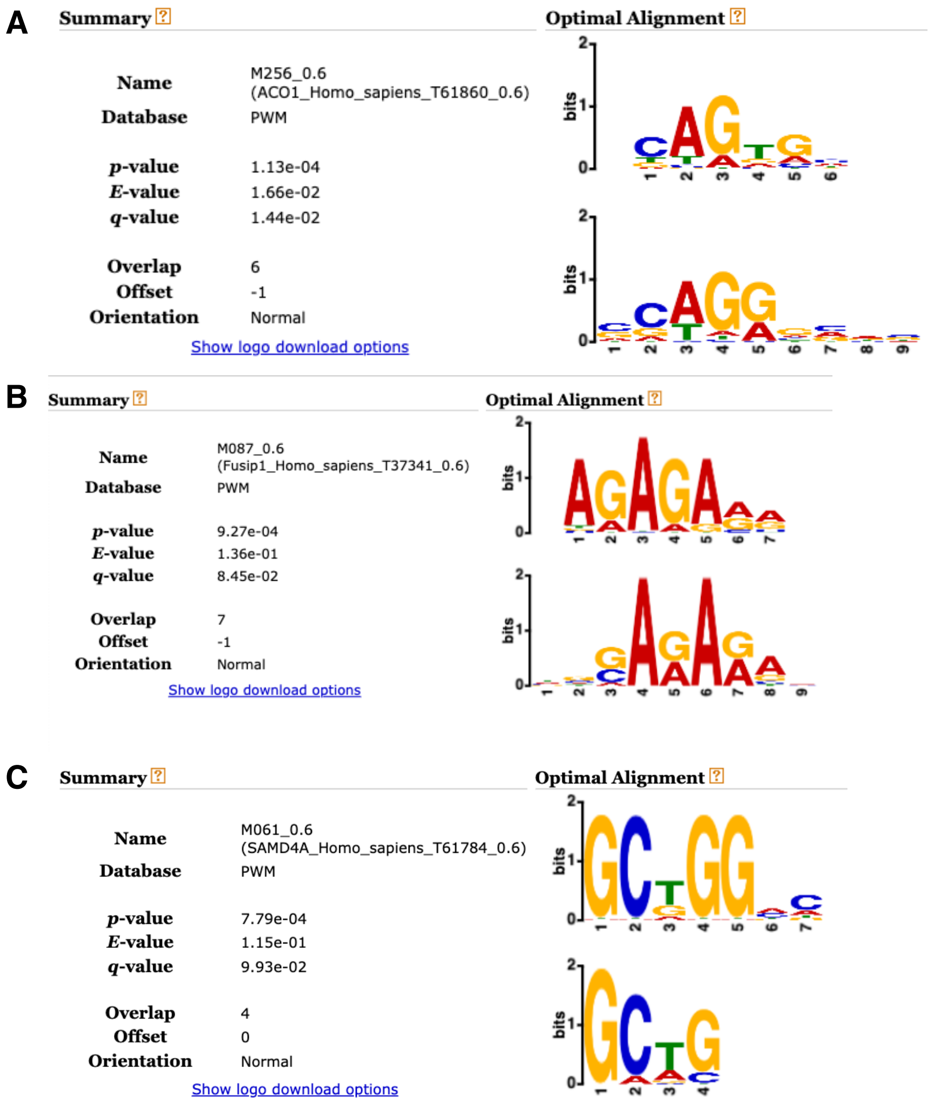


**Figure 7.** Matches with CIS-BP-RNA RNA-binding motifs from human and mouse. (A) Possible match with motif #1 from masked ISM motifs in Table S5 (B) Possible match with masked ISM motif #4 in Table S5 (C) A possible match with masked motif #1 from Table S6.

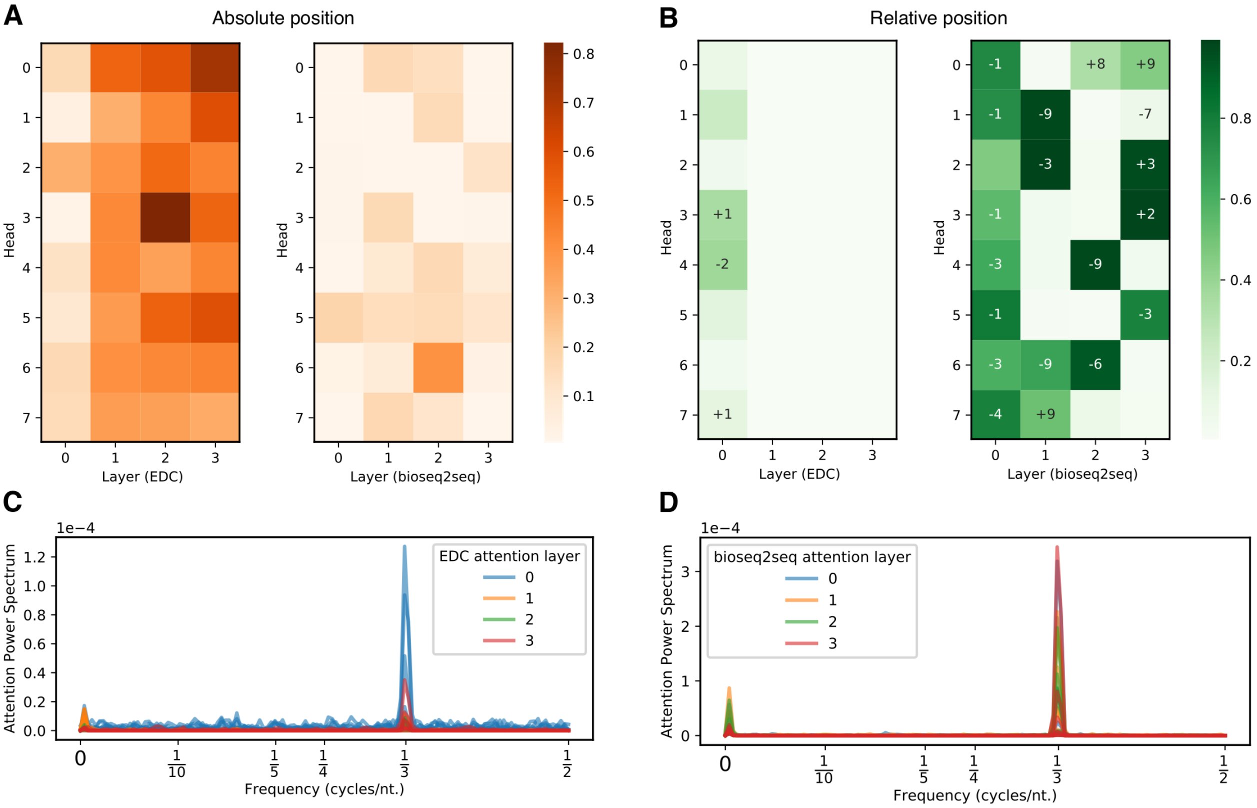


**Figure 8.** Evidence of periodicity from model version using a transformer encoder. (A) Multi-headed self-attention by transformer encoder layer, as a heatmap colored according to the average percentage of input positions in a transcript that maximally attend to a single absolute position. (B) The same as panel A but according to a relative position, with the dominant positional offset listed for each head. (C) Power Spectrum Decomposition for bioseq2seq-Transformer on mRNAs. (D) PSD for EDC-Transformer on mRNAs.

green cell with label $+1$ therefore denotes a head whose primary behavior is nearly always to pass information from the nucleotide immediately downstream of the current nucleotide.

The moderately high average values of $n_{abs}$ for the attention heads of EDC indicate that a single position within each transcript was often the largest contributor towards updating its encoder representations. By comparison, bioseq2seq devoted little attention to specific loci. However, relative positional heads were much more prevalent in bioseq2seq compared to EDC. Notably, 7/8 heads in Layer 0 of bioseq2seq had a mean value of $n_{abs} > 0.6$ and a zero-variance modal relative offset. Layer3 showed a different pattern, with four heads having mean value of $n_{abs} > 0.9$ and four heads with positional offsets at short distances divisible by three, corresponding to a range of two or three codons away. The prevalence of periodic activity in the learned transformer led us to revisit the importance of the 3-nt periodicity used in many classical gene-finding techniques, and ultimately to modify GlobalFilterNet as the basis for the LocalFilterNet (Rao et al. 2021).
