## Supplementary Table 4 for "Improving deep models of protein-coding potential with a Fourier-transform architecture and machine translation task"

| Motif # | Region | Positive Set (sites) | Negative Set (sites) | Pos. Sites | Neg. Sites | Cluster | Logo | Start site in region | Start site in window | Offset from ORF | E-value | p-value | Information |
|---|---|---|---|---|---|---|---|---|---|---|---|---|---|
| 0 | 3-prime | lncRNAs (↑ NC) | lncRNAs (random) | 1405/1984 (70.8%) | 939/1984 (47.3%) | 0 | 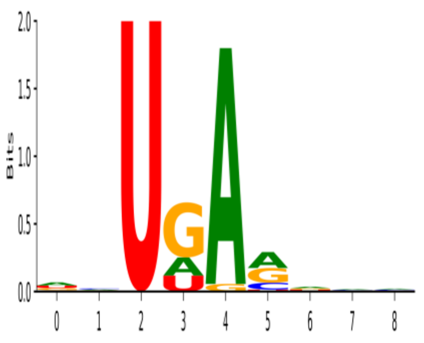 | 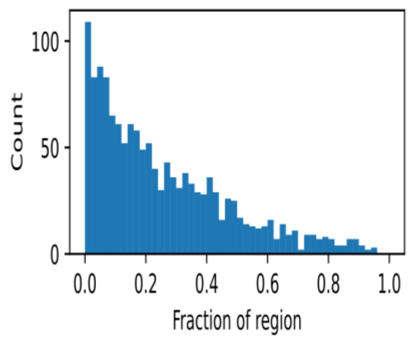 | 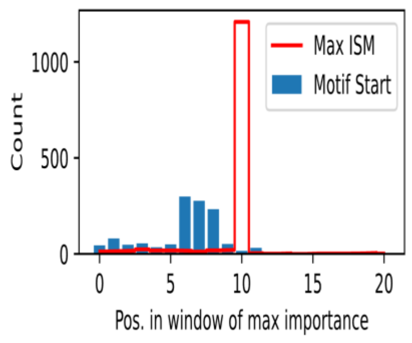 | 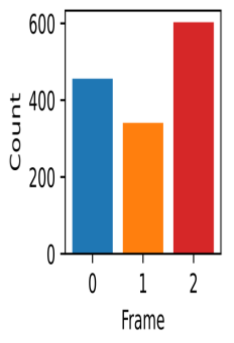 | 1.15E-04 | 8.90E-07 | 4.91 |
| 1 | ORF | mRNAs (↑ PC) | mRNAs (random) | 1298/2703 (48.0%) | 452/2703 (16.7%) | 1 | 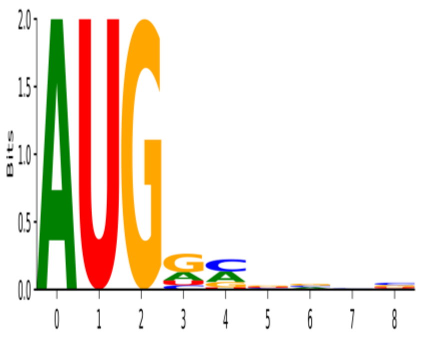 | 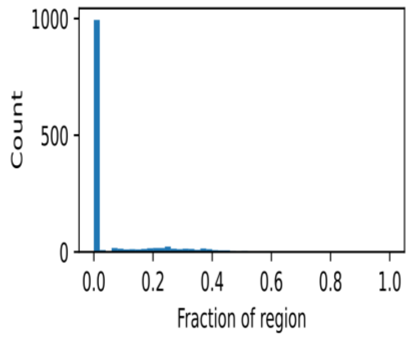 | 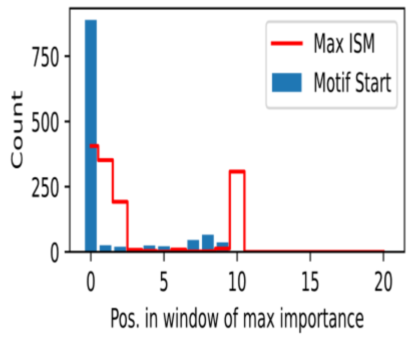 | 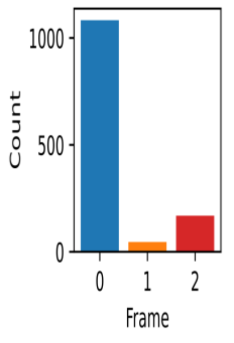 | 1.68E-15 | 1.30E-17 | 6.62 |
| 2 | ORF | lncRNAs (↑ NC) | lncRNAs (random) | 1788/2192 (81.6%) | 1051/2192 (47.9%) | 2 | 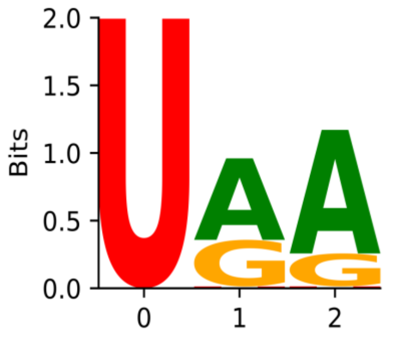 | 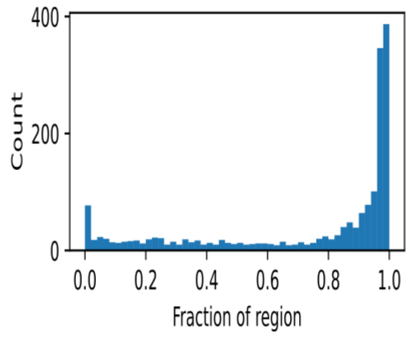 | 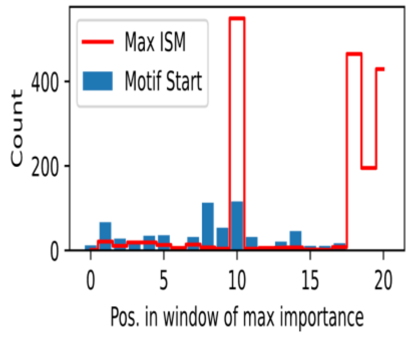 | 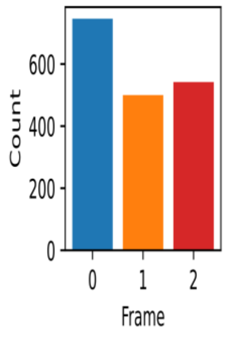 | 2.06E-14 | 1.60E-16 | 4.12 |
| 3 | ORF | mRNAs (↑ NC) | lncRNAs (↑ NC) | 1297/2703 (48.0%) | 510/2192 (23.3%) | 3 | 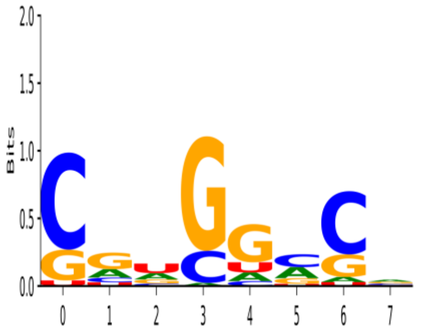 | 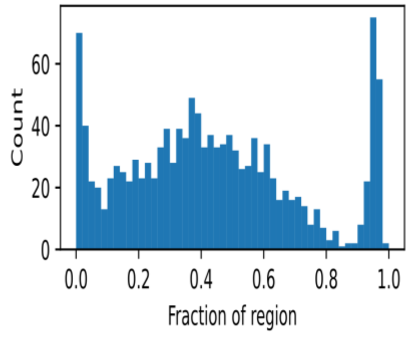 | 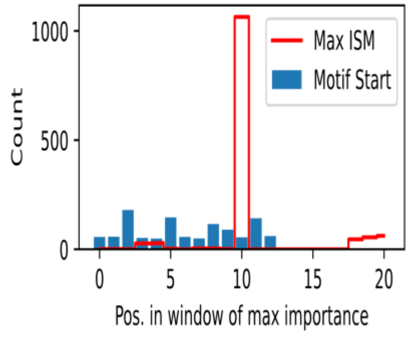 | 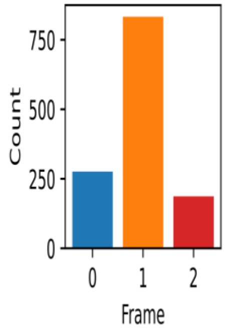 | 8.64E-05 | 6.70E-07 | 3.94 |

| Motif # | Region | Positive Set (sites) | Negative Set (sites) | Pos. Sites | Neg. Sites | Cluster | Logo | Start site in region | Start site in window | Offset from ORF | E-value | p-value | Information |
|---|---|---|---|---|---|---|---|---|---|---|---|---|---|
| 4 | ORF | lncRNAs (↑ NC) | mRNAs (↑ NC) | 1110/2192 (50.6%) | 629/2703 (23.3%) | 2 | 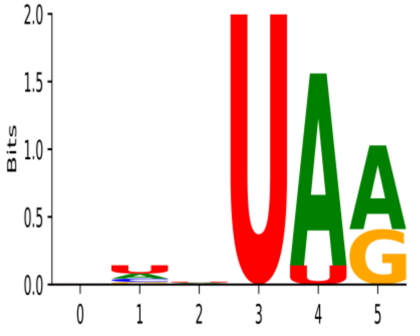 | 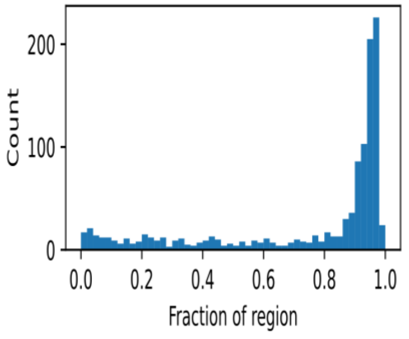 | 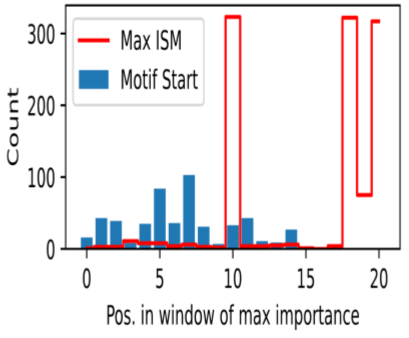 | 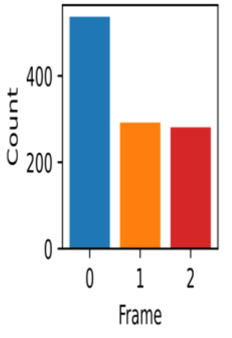 | 5.93E-08 | 4.60E-10 | 4.76 |
