## Supplementary Table 5 for "Improving deep models of protein-coding potential with a Fourier-transform architecture and machine translation task"

| Motif # | Region | Positive Set (sites) | Negative Set (sites) | Pos. Sites | Neg. Sites | Cluster | Logo | Start site in region | Start site in window | Offset from ORF | E-value | p-value | Information |
|---|---|---|---|---|---|---|---|---|---|---|---|---|---|
| 0 | 3-prime | mRNAs (↑ PC) | mRNAs (random) | 1389/1879 (73.9%) | 1000/1879 (53.2%) | 0 | 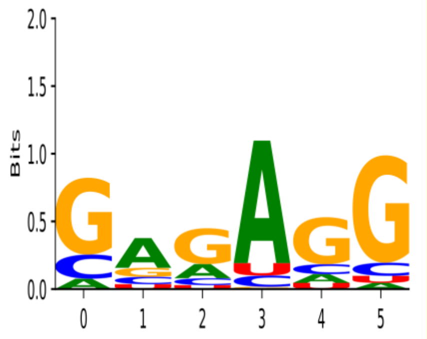 | 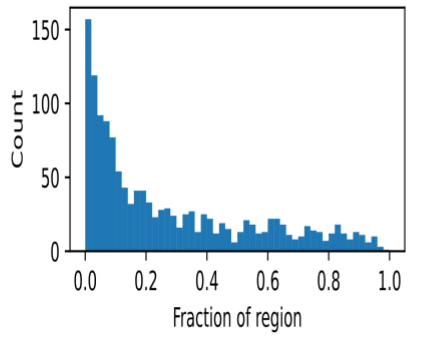 | 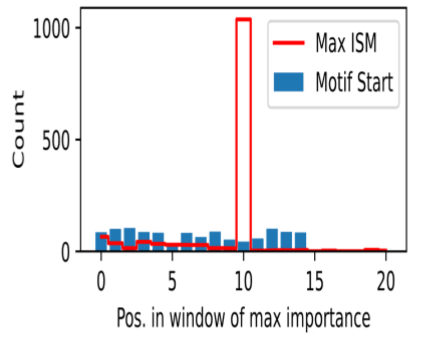 | 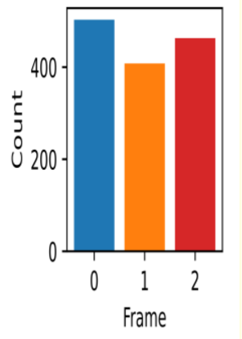 | 3.62E-04 | 2.70E-06 | 4.26 |
| 1 | 3-prime | mRNAs (↑ NC) | mRNAs (random) | 1130/1879 (60.1%) | 632/1879 (33.6%) | 1 | 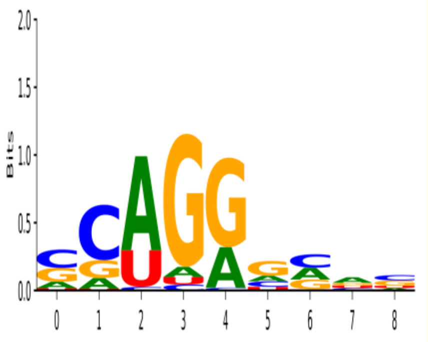 | 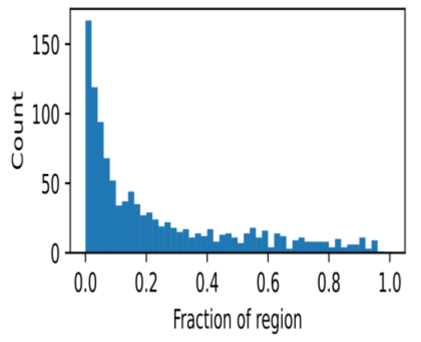 | 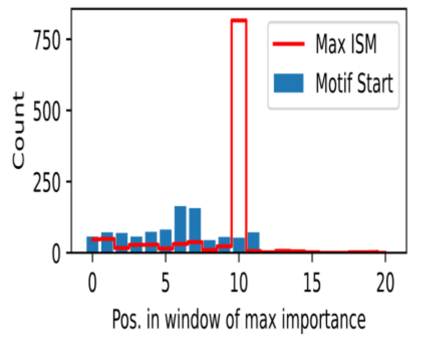 | 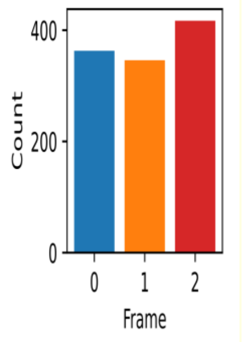 | 3.75E-06 | 2.80E-08 | 4.76 |
| 2 | 3-prime | mRNAs (↑ NC) | lncRNAs (↑ NC) | 547/1879 (29.1%) | 247/1984 (12.4%) | 2 | 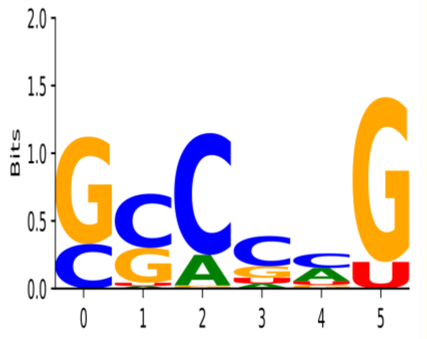 | 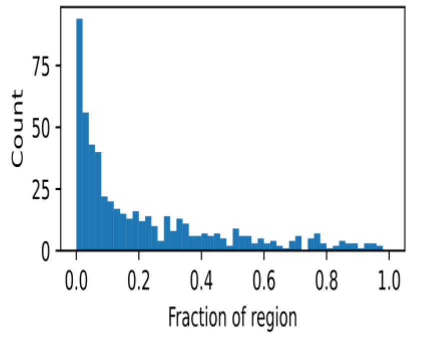 | 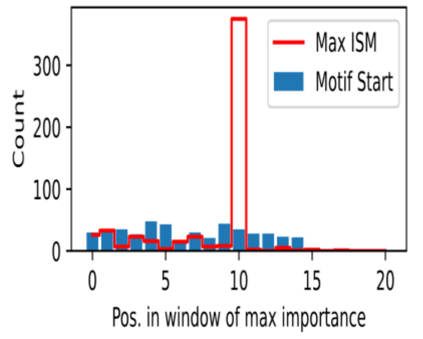 | 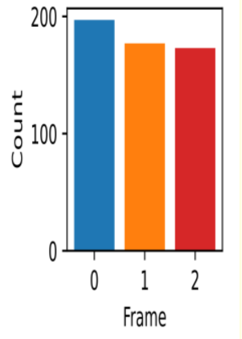 | 1.03E-04 | 7.70E-07 | 5.03 |
| 3 | 3-prime | lncRNAs (↑ NC) | mRNAs (↑ NC) | 1322/1984 (66.6%) | 860/1879 (45.8%) | 3 | 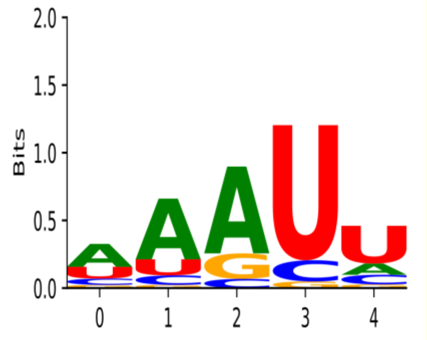 | 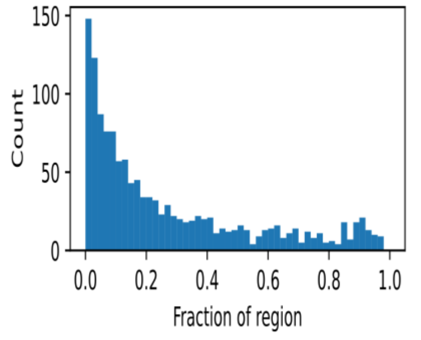 | 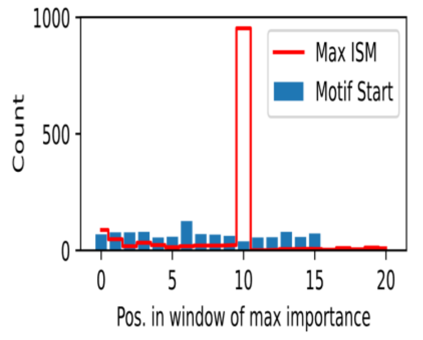 | 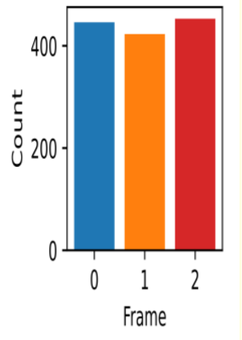 | 1.47E-04 | 1.10E-06 | 3.55 |

| Motif # | Region | Positive Set (sites) | Negative Set (sites) | Pos. Sites | Neg. Sites | Cluster | Logo | Start site in region | Start site in window | Offset from ORF | E-value | p-value | Information |
|---|---|---|---|---|---|---|---|---|---|---|---|---|---|
| 4 | ORF | mRNAs (↑ PC) | mRNAs (random) | 1277/2703 (47.2%) | 370/2703 (13.7%) | 4 | 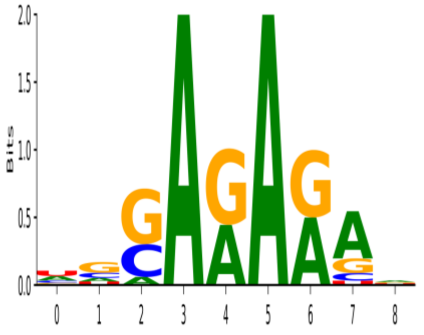 | 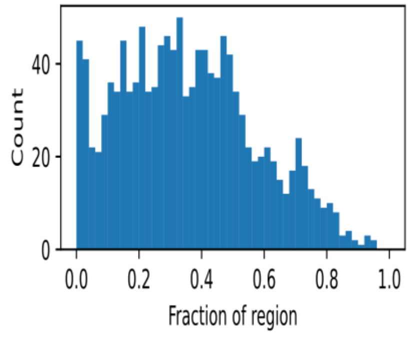 | 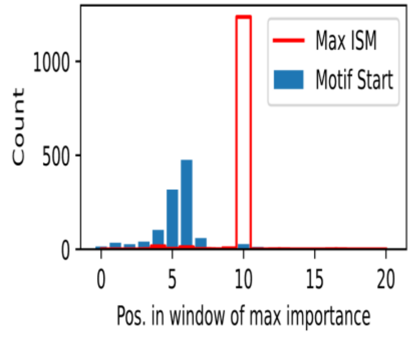 | 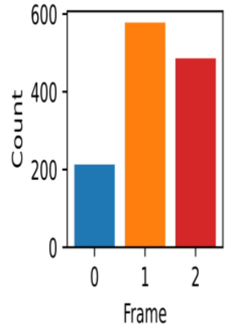 | 1.17E-17 | 8.70E-20 | 7.57 |
| 5 | ORF | mRNAs (↑ PC) | lncRNAs (↑ PC) | 897/2703 (33.2%) | 354/2192 (16.1%) | 0 | 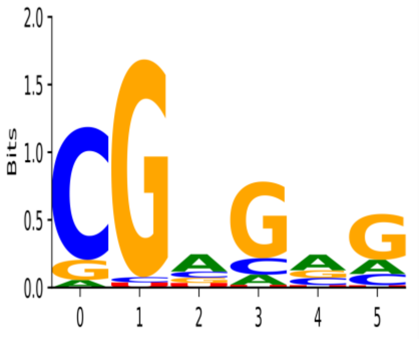 | 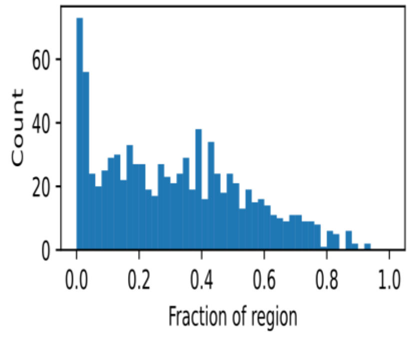 | 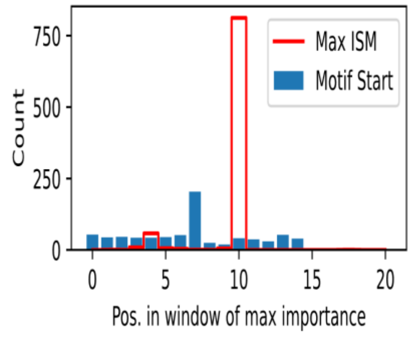 | 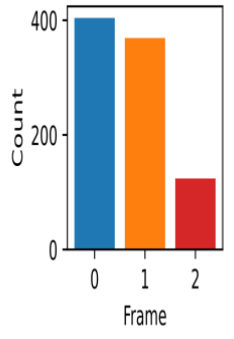 | 3.22E-06 | 2.40E-08 | 4.69 |
| 6 | ORF | lncRNAs (↑ PC) | mRNAs (↑ PC) | 842/2192 (38.4%) | 552/2703 (20.4%) | 5 | 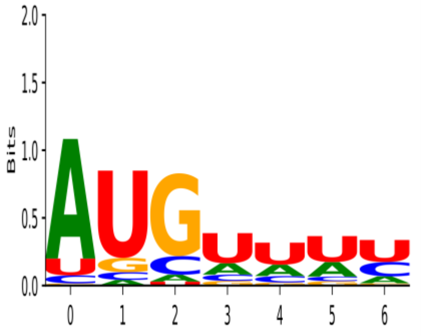 | 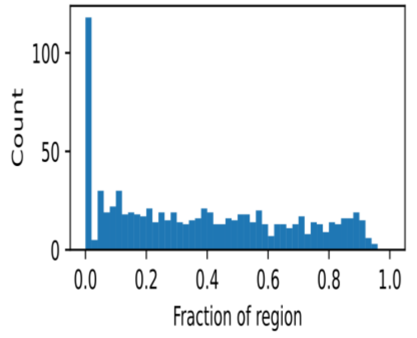 | 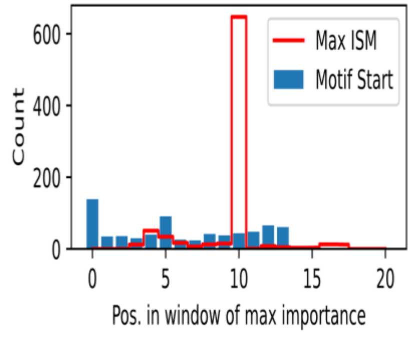 | 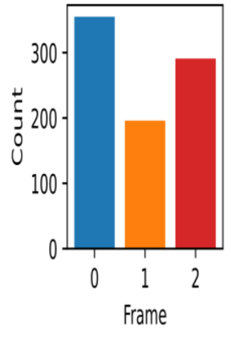 | 7.77E-04 | 5.80E-06 | 4.17 |
