## Supplementary Table 6 for "Improving deep models of protein-coding potential with a Fourier-transform architecture and machine translation task"

| Motif # | Region | Positive Set (sites) | Negative Set (sites) | Pos. Sites | Neg. Sites | Cluster | Logo | Start site in region | Start site in window | Offset from ORF | E-value | p-value | Information |
|---|---|---|---|---|---|---|---|---|---|---|---|---|---|
| 0 | 3-prime | mRNAs (↑ PC) | mRNAs (random) | 12660/18151 (69.7%) | 9109/18151 (50.2%) | 0 | 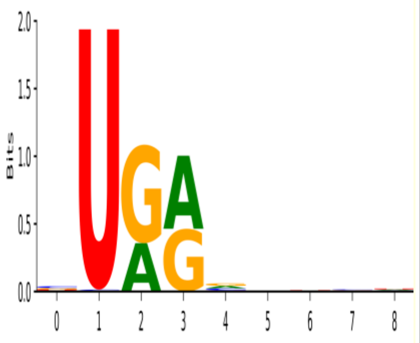 | 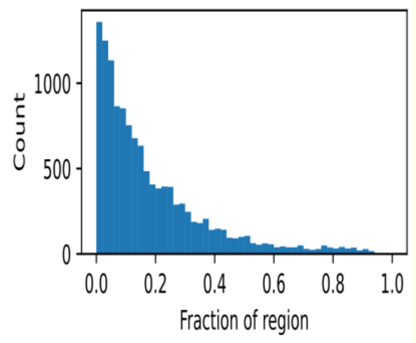 | 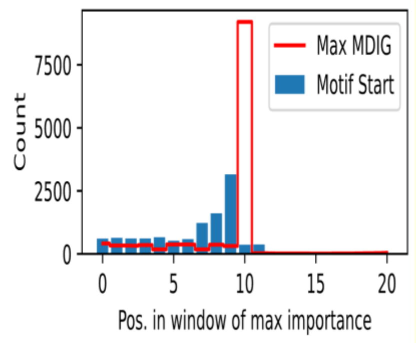 | 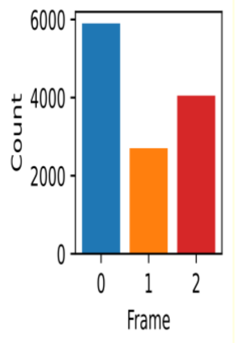 | 4.36E-35 | 2.00E-37 | 4.18 |
| 1 | 3-prime | mRNAs (↑ NC) | mRNAs (random) | 11988/18151 (66.0%) | 8241/18151 (45.4%) | 1 | 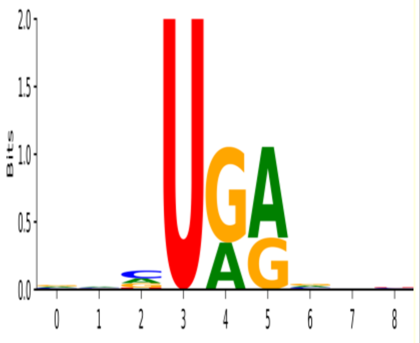 | 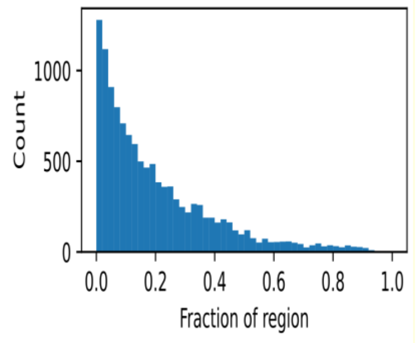 | 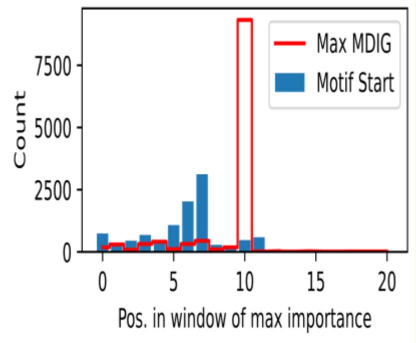 | 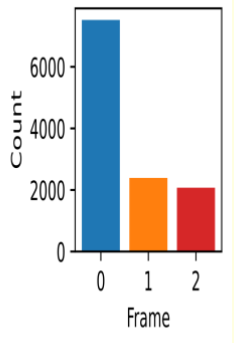 | 1.87E-29 | 8.60E-32 | 4.37 |
| 2 | 3-prime | lncRNAs (↑ PC) | lncRNAs (random) | 19817/22525 (88.0%) | 14971/22525 (66.5%) | 0 | 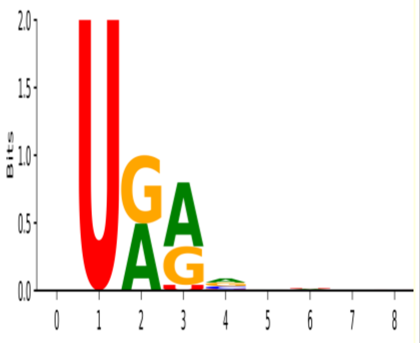 | 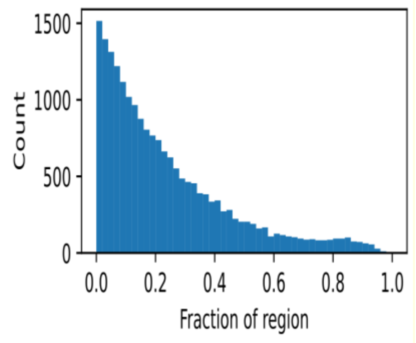 | 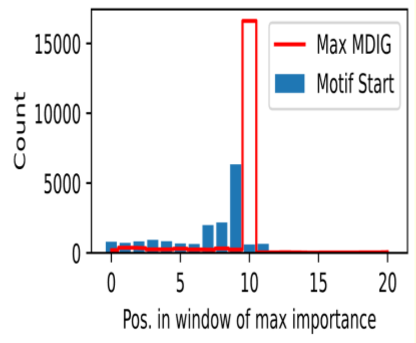 | 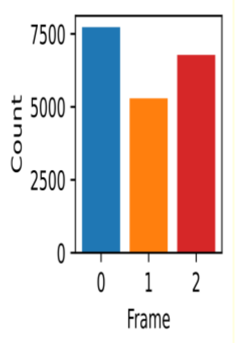 | 5.23E-48 | 2.40E-50 | 3.93 |
| 3 | 3-prime | lncRNAs (↑ NC) | lncRNAs (random) | 20568/22525 (91.3%) | 14035/22525 (62.3%) | 0 | 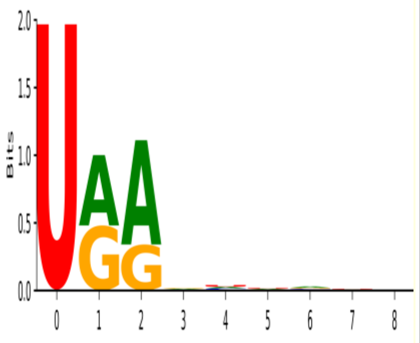 | 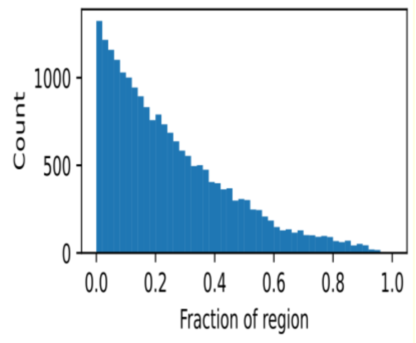 | 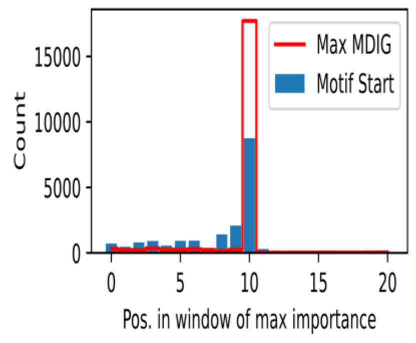 | 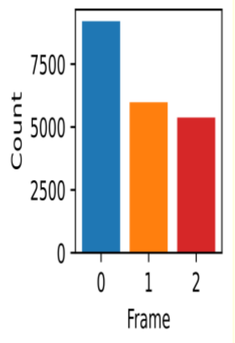 | 2.09E-120 | 9.60E-123 | 4.20 |

| Motif # | Region | Positive Set (sites) | Negative Set (sites) | Pos. Sites | Neg. Sites | Cluster | Logo | Start site in region | Start site in window | Offset from ORF | E-value | p-value | Information |
|---|---|---|---|---|---|---|---|---|---|---|---|---|---|
| 4 | 3-prime | lncRNAs (↑ NC) | mRNAs (↑ NC) | 20800/22525 (92.3%) | 14147/18151 (77.9%) | 0 | 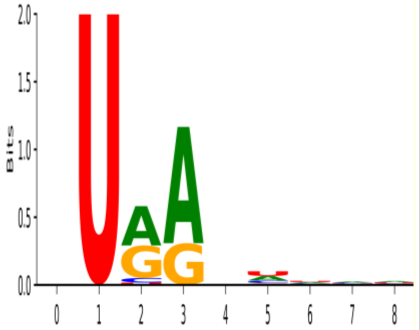 | 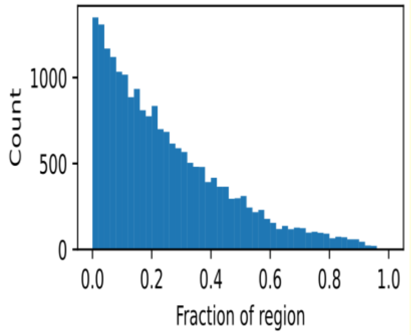 | 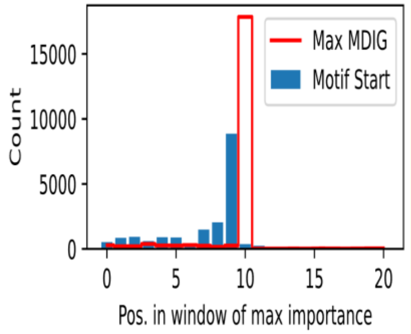 | 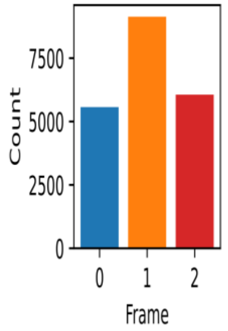 | 2.40E-44 | 1.10E-46 | 3.94 |
| 5 | 5-prime | mRNAs (↑ PC) | mRNAs (random) | 3847/15842 (24.3%) | 1498/15842 (9.5%) | 2 | 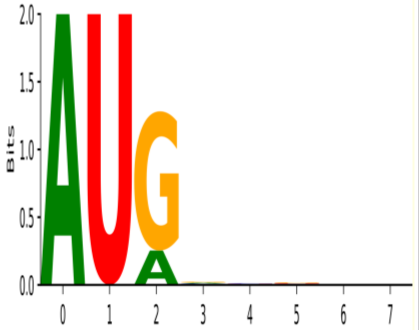 | 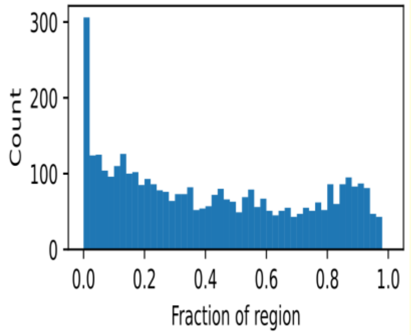 | 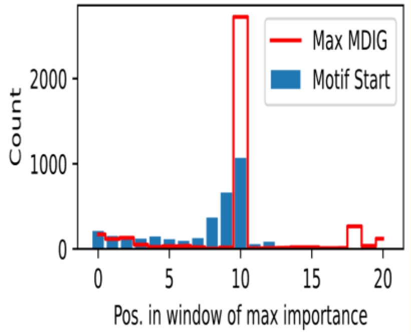 | 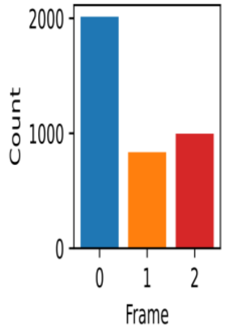 | 1.85E-19 | 8.50E-22 | 5.34 |
| 6 | 5-prime | mRNAs (↑ NC) | mRNAs (random) | 10868/15842 (68.6%) | 6358/15842 (40.1%) | 1 | 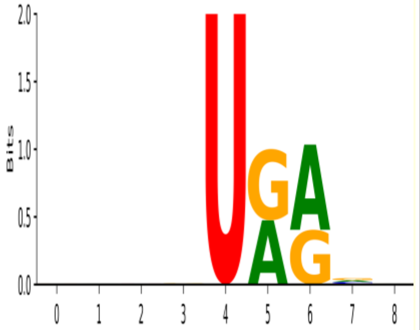 | 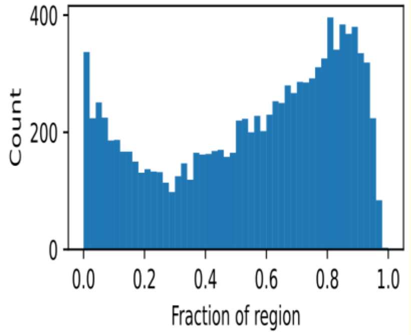 | 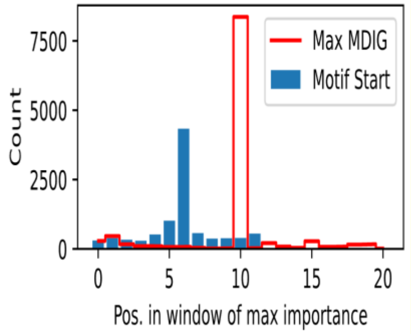 | 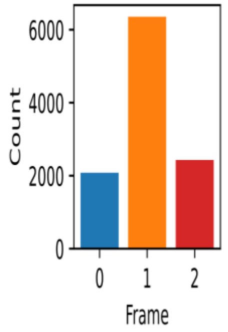 | 6.76E-55 | 3.10E-57 | 4.12 |
| 7 | 5-prime | lncRNAs (↑ PC) | lncRNAs (random) | 10720/23017 (46.6%) | 3743/23017 (16.3%) | 2 | 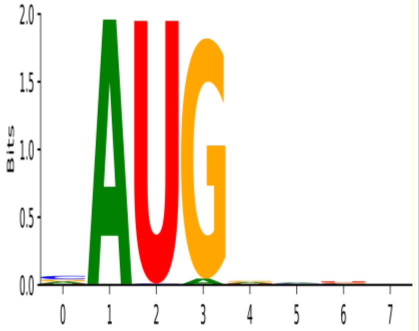 | 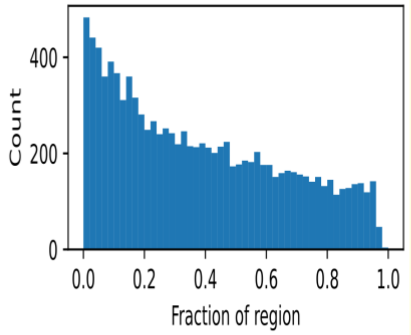 | 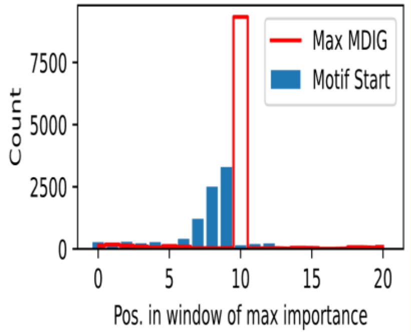 | 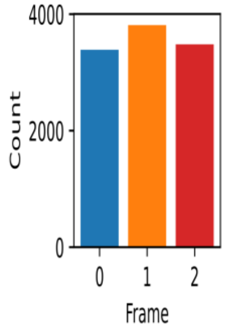 | 8.94E-98 | 4.10E-100 | 5.88 |

| Motif # | Region | Positive Set (sites) | Negative Set (sites) | Pos. Sites | Neg. Sites | Cluster | Logo | Start site in region | Start site in window | Offset from ORF | E-value | p-value | Information |
|---|---|---|---|---|---|---|---|---|---|---|---|---|---|
| 8 | 5-prime | lncRNAs (↑ PC) | lncRNAs (random) | 8477/23017 (36.8%) | 6918/23017 (30.1%) | 3 | 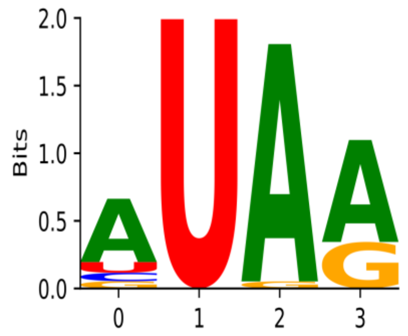 | 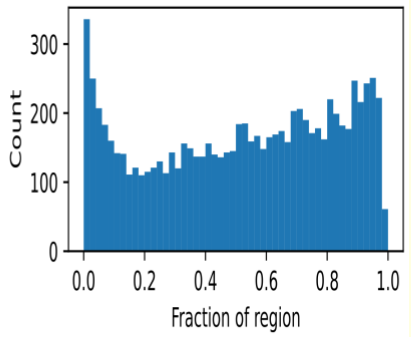 | 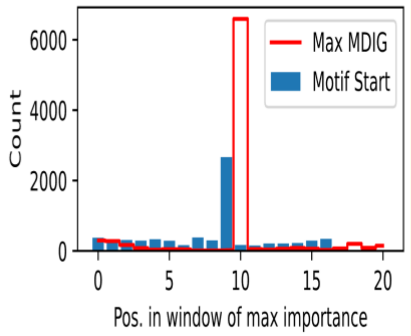 | 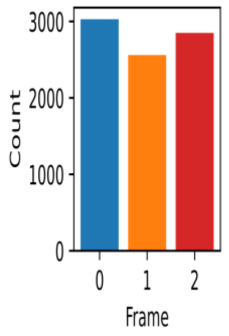 | 1.11E-05 | 5.10E-08 | 5.56 |
| 9 | 5-prime | lncRNAs (↑ NC) | lncRNAs (random) | 19527/23017 (84.8%) | 10954/23017 (47.6%) | 0 | 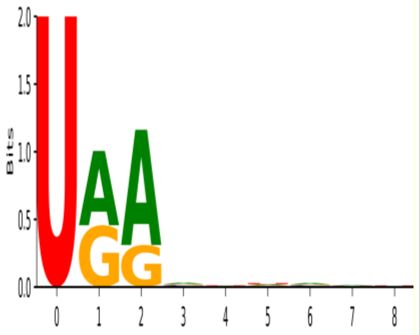 | 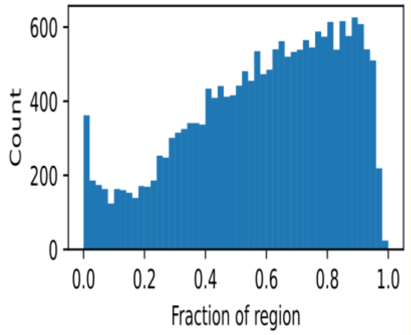 | 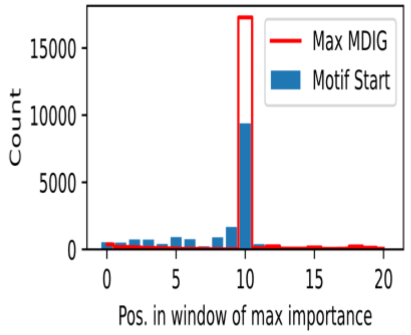 | 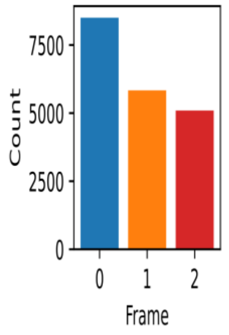 | 1.77E-164 | 8.10E-167 | 4.30 |
| 10 | ORF | mRNAs (↑ PC) | mRNAs (random) | 20837/26039 (80.0%) | 5327/26039 (20.5%) | 2 | 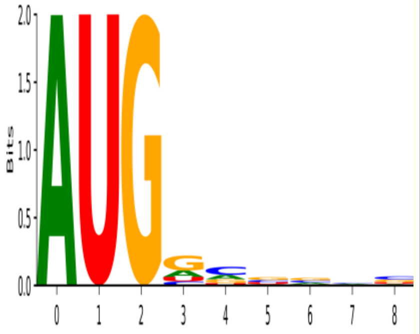 | 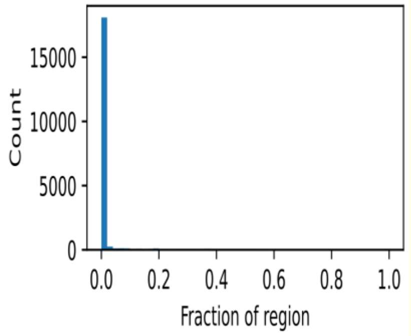 | 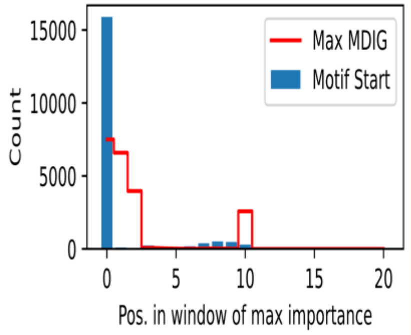 | 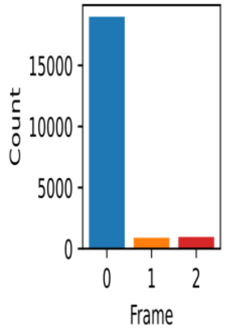 | 0.00E+00 | 0.00E+00 | 6.56 |
| 11 | ORF | mRNAs (↑ NC) | mRNAs (random) | 14400/26039 (55.3%) | 6927/26039 (26.6%) | 4 | 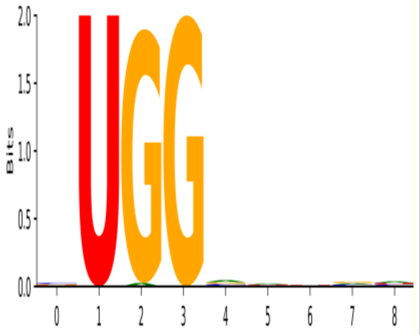 | 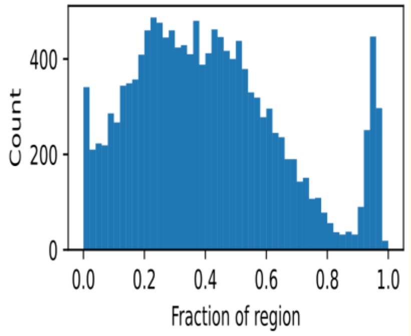 | 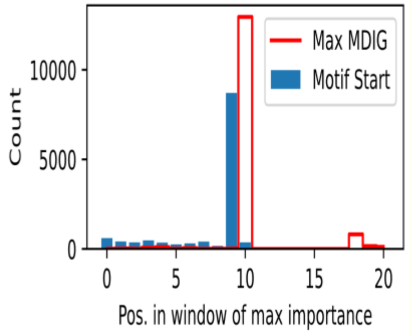 | 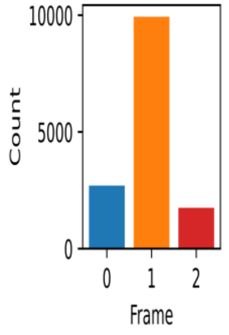 | 5.67E-96 | 2.60E-98 | 6.08 |

| Motif # | Region | Positive Set (sites) | Negative Set (sites) | Pos. Sites | Neg. Sites | Cluster | Logo | Start site in region | Start site in window | Offset from ORF | E-value | p-value | Information |
|---|---|---|---|---|---|---|---|---|---|---|---|---|---|
| 12 | ORF | lncRNAs (↑ PC) | lncRNAs (random) | 14490/24881 (58.2%) | 5202/24881 (20.9%) | 2 | 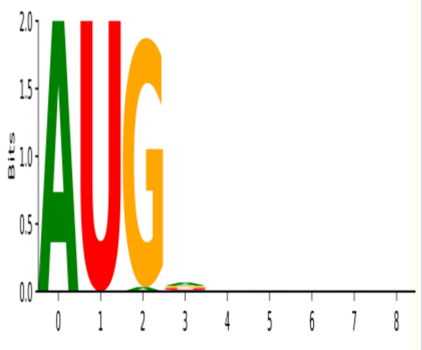 | 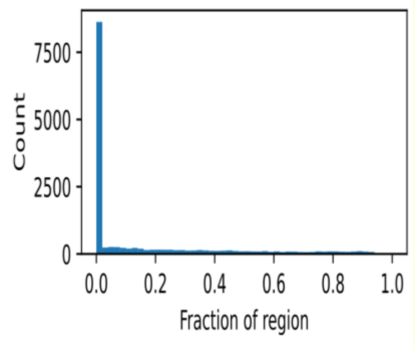 | 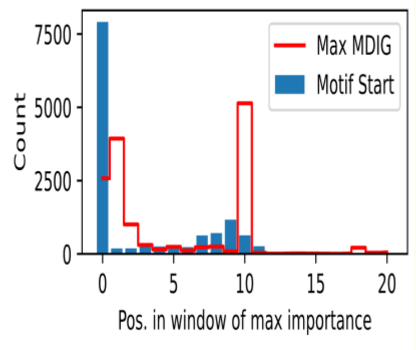 | 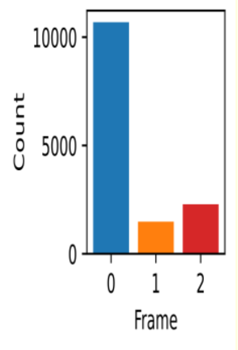 | 3.71E-170 | 1.70E-172 | 5.97 |
| 13 | ORF | lncRNAs (↑ PC) | lncRNAs (random) | 3852/24881 (15.5%) | 2629/24881 (10.6%) | 3 | 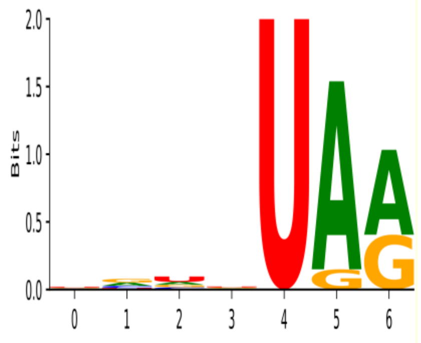 | 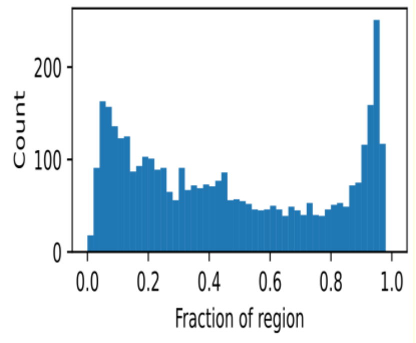 | 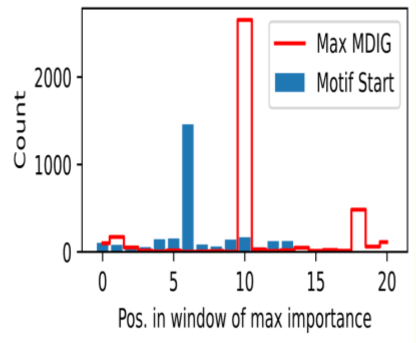 | 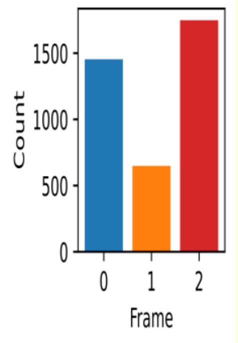 | 1.13E-08 | 5.20E-11 | 4.79 |
| 14 | ORF | lncRNAs (↑ NC) | lncRNAs (random) | 21958/24881 (88.3%) | 12882/24881 (51.8%) | 1 | 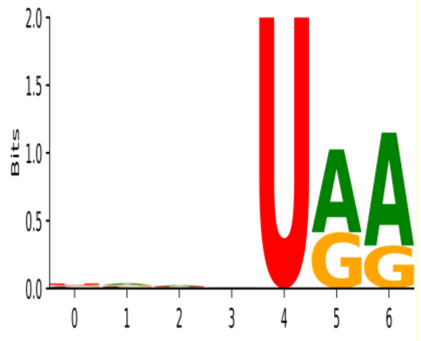 | 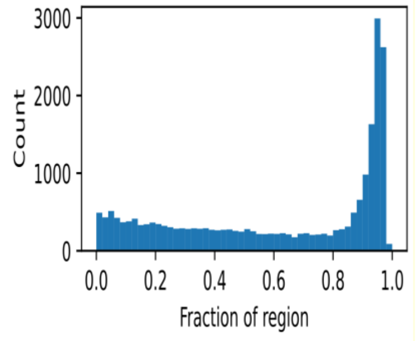 | 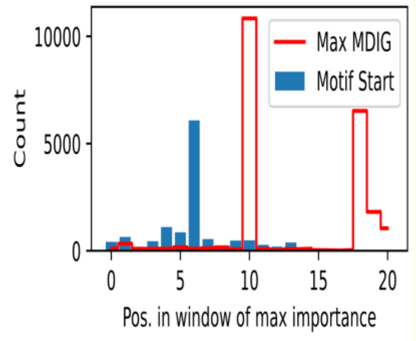 | 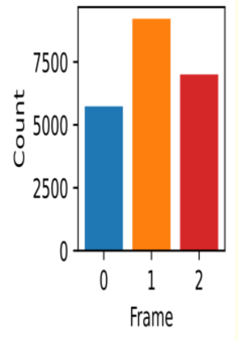 | 2.18E-173 | 1.00E-175 | 4.28 |
| 15 | ORF | lncRNAs (↑ NC) | lncRNAs (random) | 1245/24881 (5.0%) | 366/24881 (1.5%) | 5 | 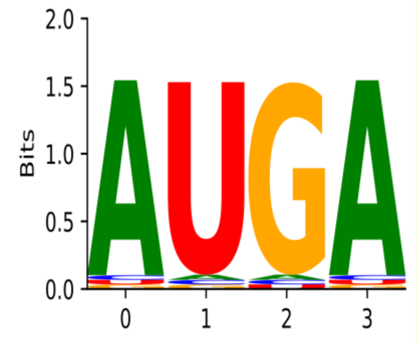 | 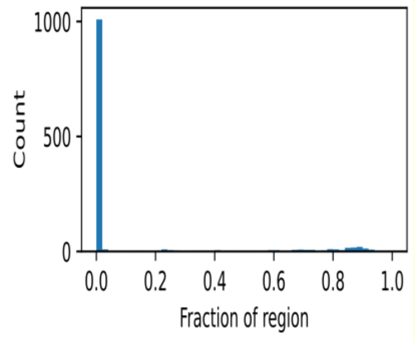 | 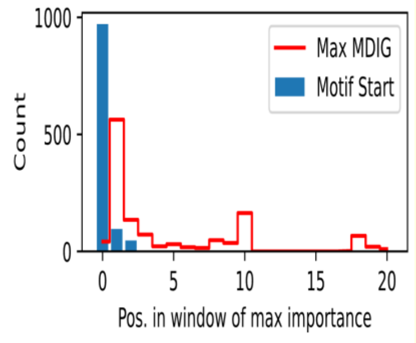 | 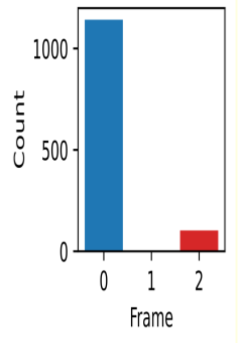 | 4.14E-15 | 1.90E-17 | 6.14 |

| Motif # | Region | Positive Set (sites) | Negative Set (sites) | Pos. Sites | Neg. Sites | Cluster | Logo | Start site in region | Start site in window | Offset from ORF | E-value | p-value | Information |
|---|---|---|---|---|---|---|---|---|---|---|---|---|---|
| 16 | ORF | mRNAs (↑ NC) | lncRNAs (↑ NC) | 18225/26039 (70.0%) | 10907/24881 (43.8%) | 4 | 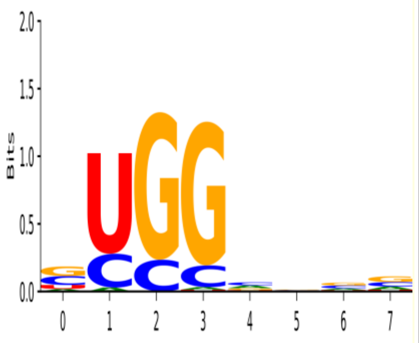 | 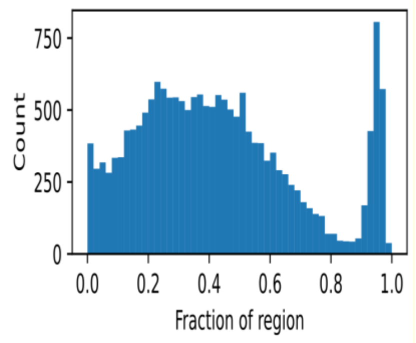 | 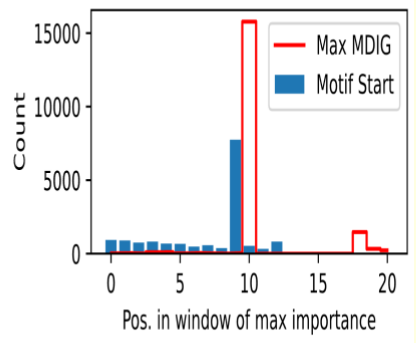 | 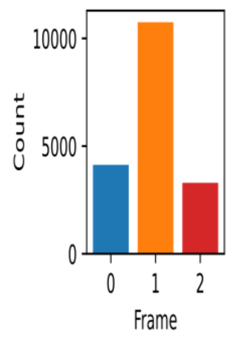 | 2.16E-77 | 9.90E-80 | 4.05 |
| 17 | ORF | lncRNAs (↑ NC) | mRNAs (↑ NC) | 1936/24881 (7.8%) | 1252/26039 (4.8%) | 6 | 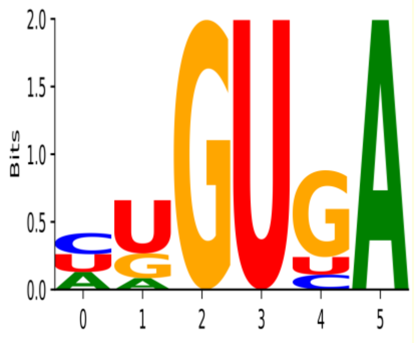 | 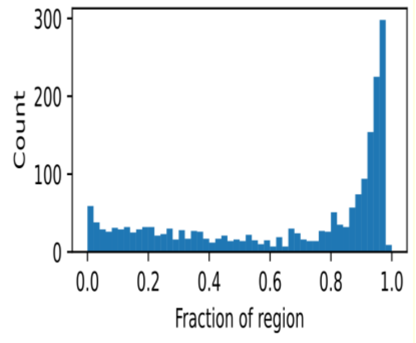 | 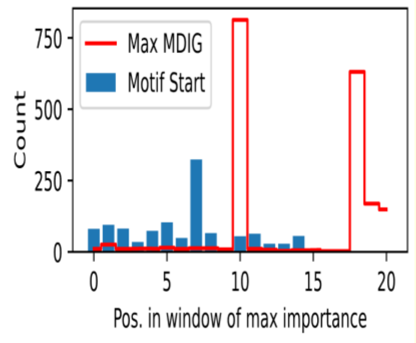 | 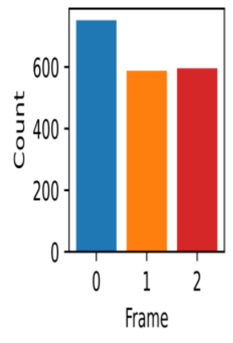 | 4.80E-05 | 2.20E-07 | 7.93 |
| 18 | ORF | lncRNAs (↑ NC) | mRNAs (↑ NC) | 1917/24881 (7.7%) | 1409/26039 (5.4%) | 7 | 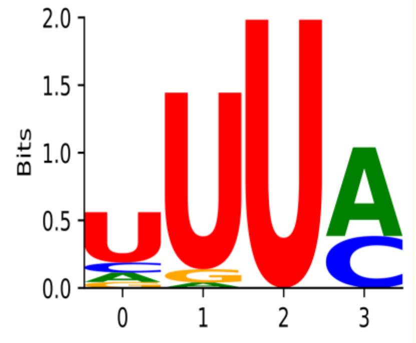 | 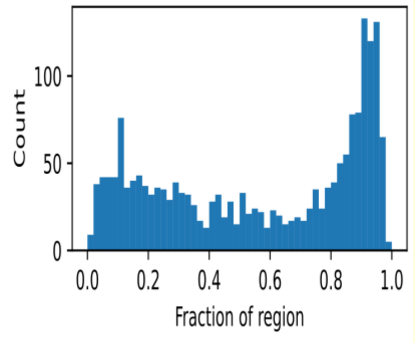 | 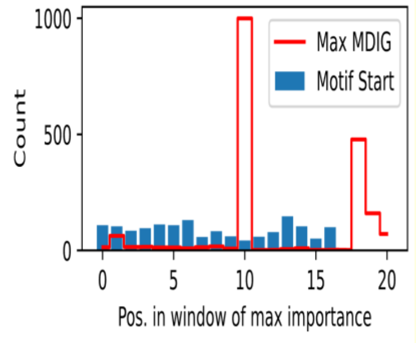 | 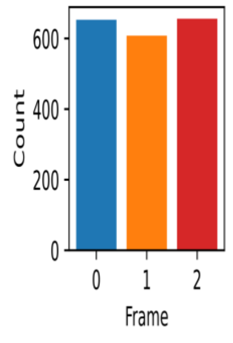 | 2.40E-04 | 1.10E-06 | 5.03 |
