## Supplementary Table 7 for "Improving deep models of protein-coding potential with a Fourier-transform architecture and machine translation task"

| Motif # | Region | Positive Set (sites) | Negative Set (sites) | Pos. Sites | Neg. Sites | Cluster | Logo | Start site in region | Start site in window | Offset from ORF | E-value | p-value | Information |
|---|---|---|---|---|---|---|---|---|---|---|---|---|---|
| 0 | 3-prime | mRNAs (↑ PC) | mRNAs (random) | 10895/18151 (60.0%) | 8366/18151 (46.1%) | 0 | 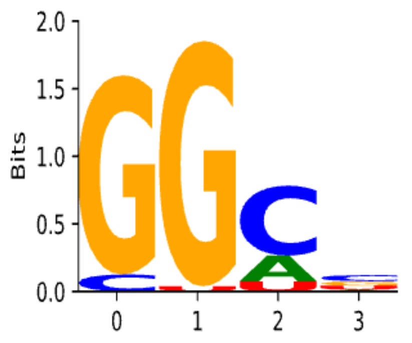 | 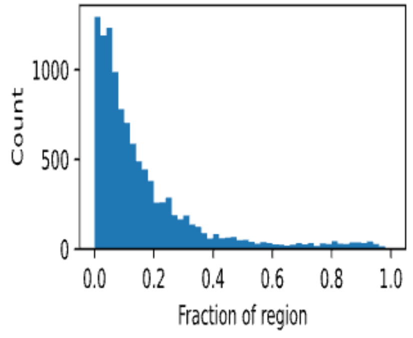 | 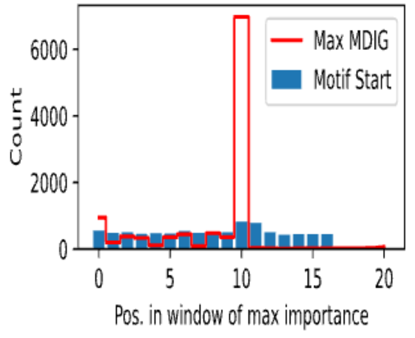 | 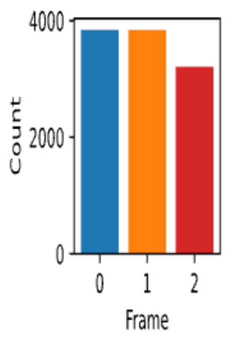 | 3.65E-07 | 1.60E-09 | 4.36 |
| 1 | 3-prime | mRNAs (↑ PC) | mRNAs (random) | 5850/18151 (32.2%) | 4728/18151 (26.0%) | 1 | 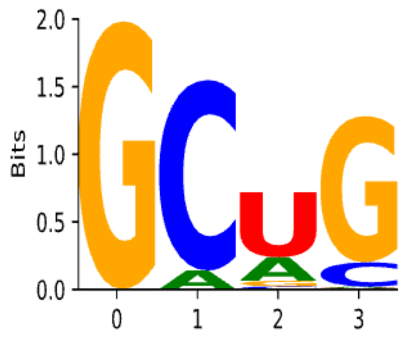 | 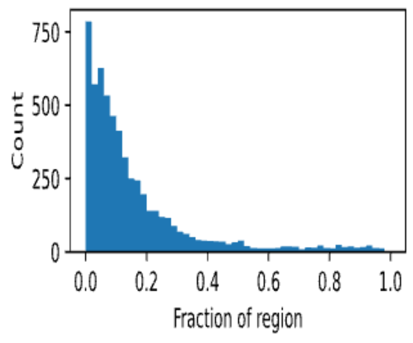 | 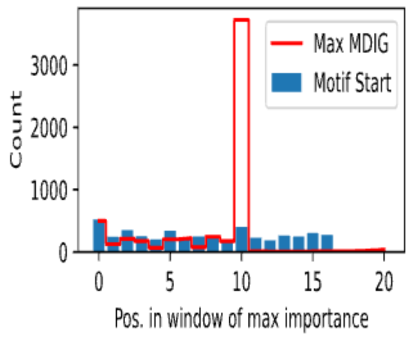 | 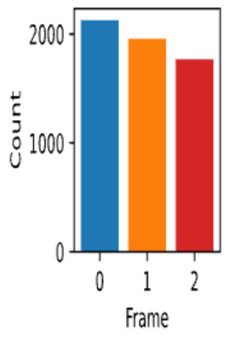 | 2.96E-04 | 1.30E-06 | 5.53 |
| 2 | 3-prime | mRNAs (↑ NC) | mRNAs (random) | 11803/18151 (65.0%) | 8335/18151 (45.9%) | 2 | 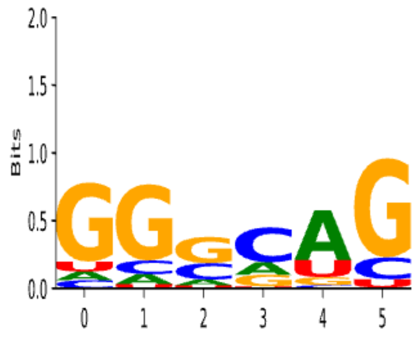 | 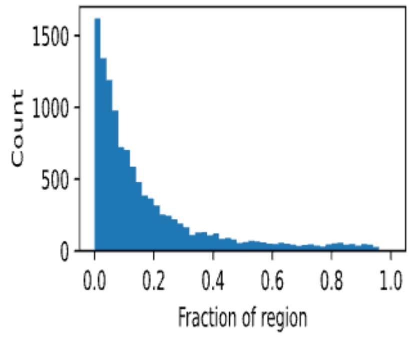 | 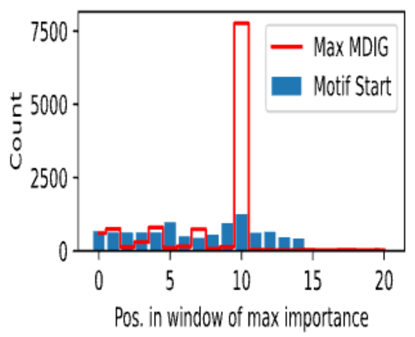 | 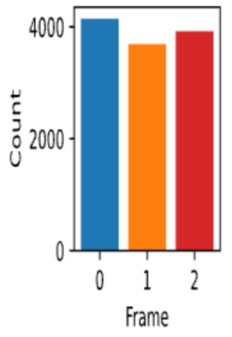 | 1.76E-26 | 7.70E-29 | 3.92 |
| 3 | 3-prime | lncRNAs (↑ PC) | lncRNAs (random) | 15804/22525 (70.2%) | 13061/22525 (58.0%) | 3 | 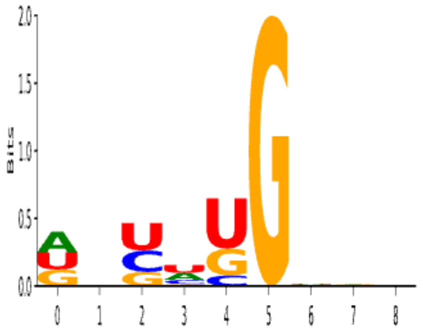 | 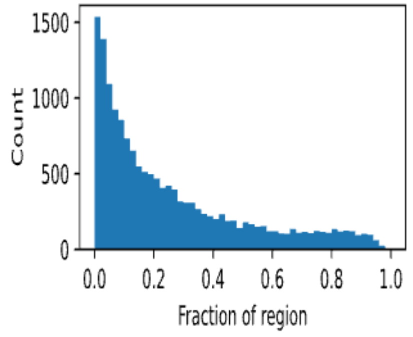 | 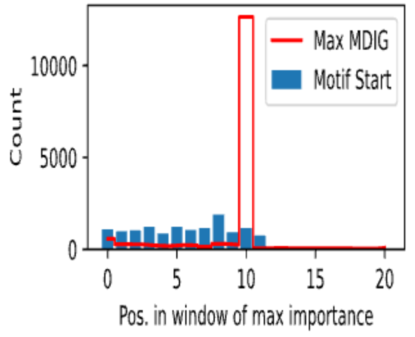 | 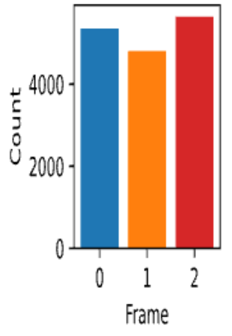 | 1.05E-15 | 4.60E-18 | 3.71 |

| Motif # | Region | Positive Set (sites) | Negative Set (sites) | Pos. Sites | Neg. Sites | Cluster | Logo | Start site in region | Start site in window | Offset from ORF | E-value | p-value | Information |
|---|---|---|---|---|---|---|---|---|---|---|---|---|---|
| 4 | 3-prime | lncRNAs (↑ NC) | lncRNAs (random) | 7821/22525 (34.7%) | 5366/22525 (23.8%) | 4 | 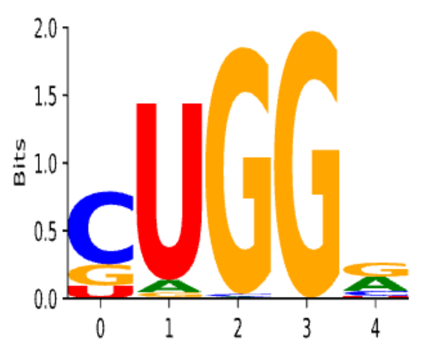 | 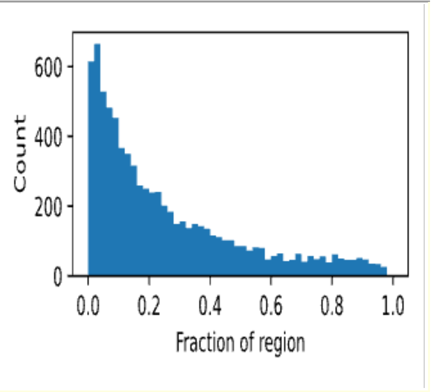 | 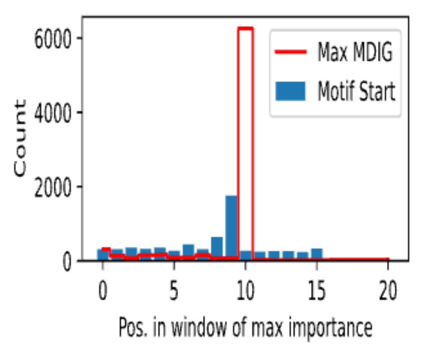 | 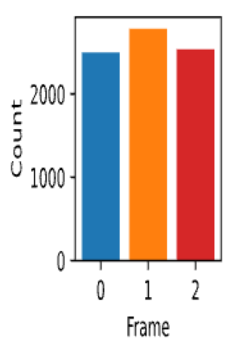 | 1.00E-09 | 4.40E-12 | 6.32 |
| 5 | 5-prime | mRNAs (↑ PC) | mRNAs (random) | 9175/15842 (57.9%) | 7655/15842 (48.3%) | 5 | 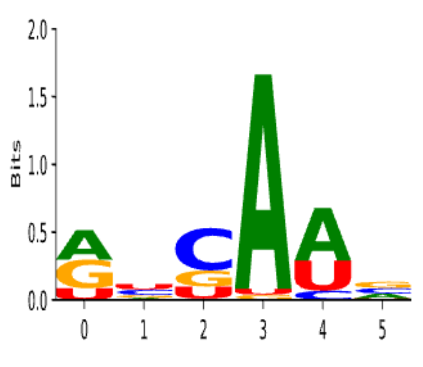 | 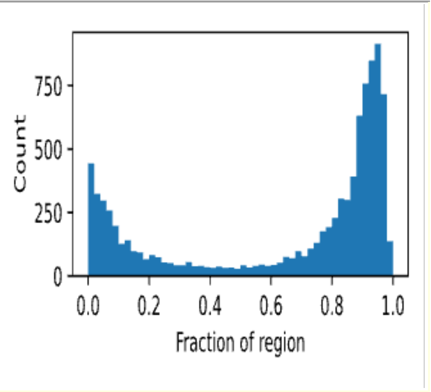 | 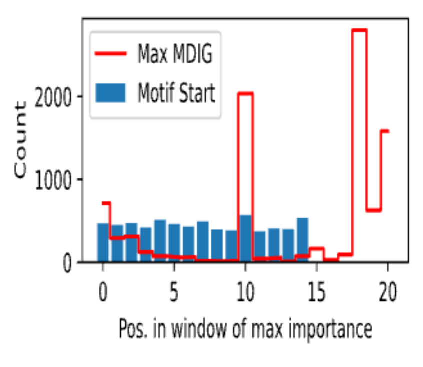 | 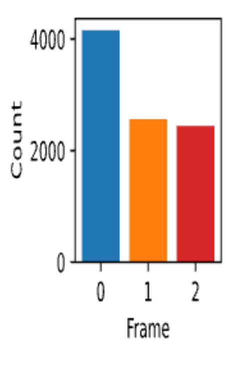 | 1.19E-04 | 5.20E-07 | 3.65 |
| 6 | 5-prime | lncRNAs (↑ PC) | lncRNAs (random) | 7004/23017 (30.4%) | 4949/23017 (21.5%) | 3 | 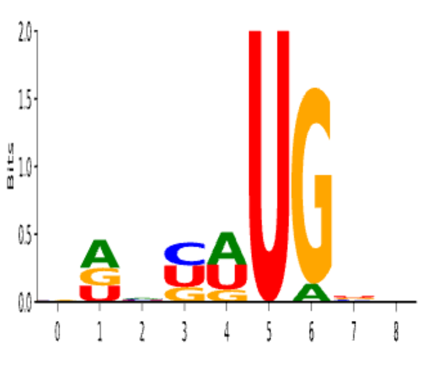 | 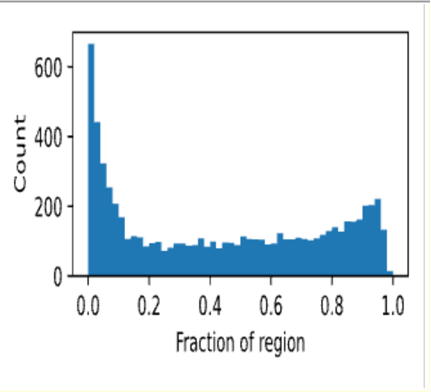 | 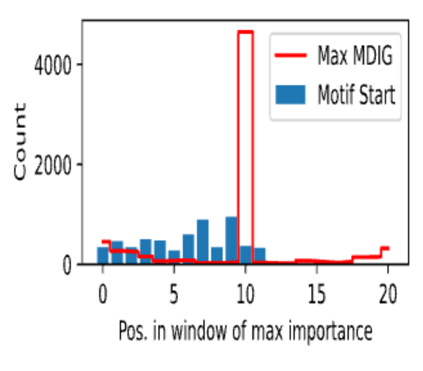 | 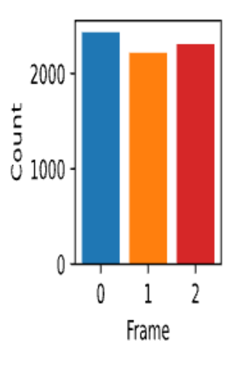 | 8.66E-10 | 3.80E-12 | 5.09 |
| 7 | 5-prime | lncRNAs (↑ NC) | lncRNAs (random) | 10017/23017 (43.5%) | 6633/23017 (28.8%) | 6 | 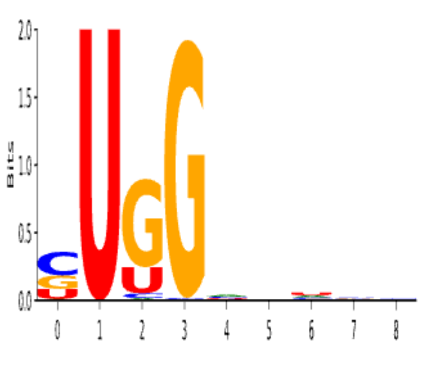 | 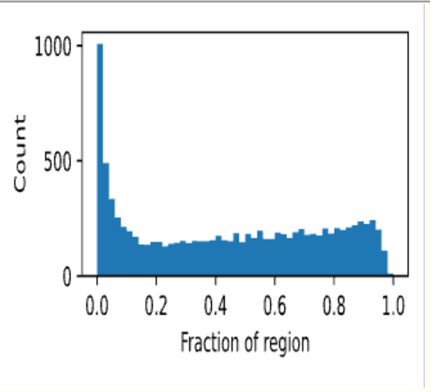 | 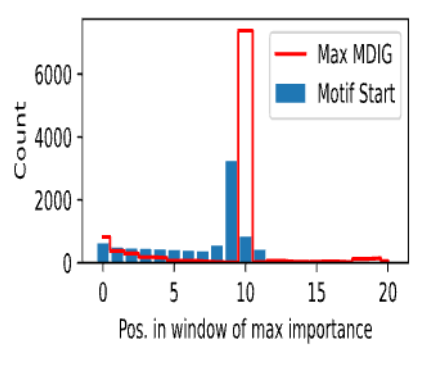 | 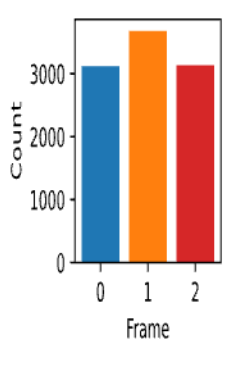 | 7.98E-25 | 3.50E-27 | 5.32 |

| Motif # | Region | Positive Set (sites) | Negative Set (sites) | Pos. Sites | Neg. Sites | Cluster | Logo | Start site in region | Start site in window | Offset from ORF | E-value | p-value | Information |
|---|---|---|---|---|---|---|---|---|---|---|---|---|---|
| 8 | ORF | mRNAs (↑ PC) | mRNAs (random) | 5940/26039 (22.8%) | 2650/26039 (10.2%) | 7 | 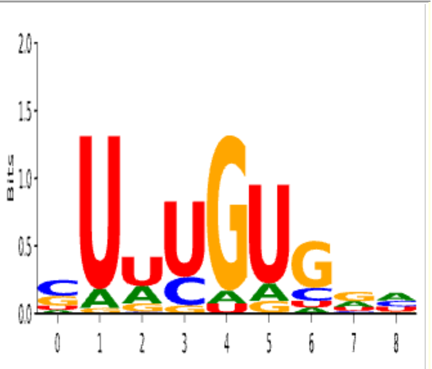  | 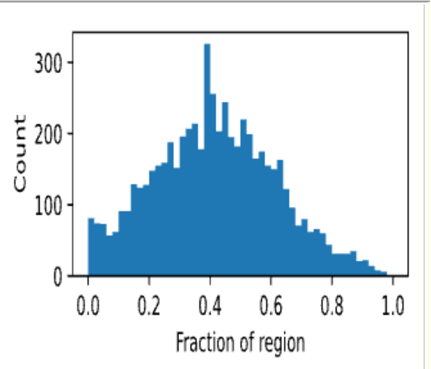  | 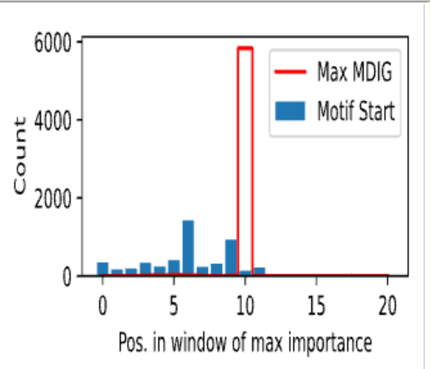  | 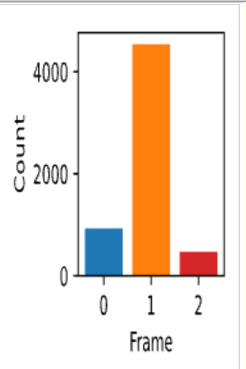  | 2.51E-32 | 1.10E-34 | 5.93 |
| 9 | ORF | mRNAs (↑ PC) | mRNAs (random) | 2746/26039 (10.5%) | 1587/26039 (6.1%) | 0 | 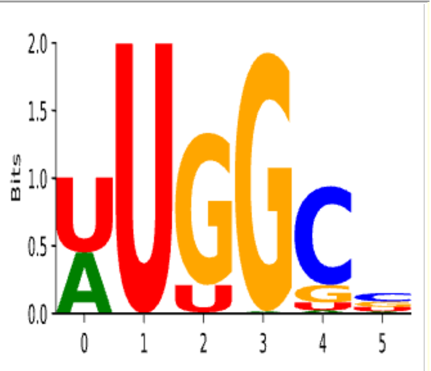  | 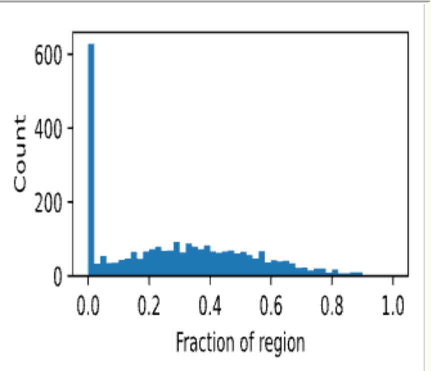  | 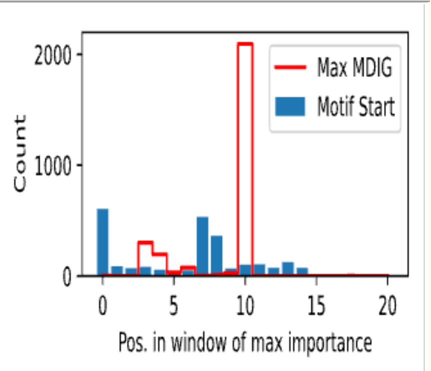  | 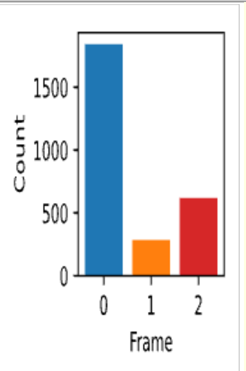  | 2.51E-05 | 1.10E-07 | 7.36 |
| 10 | ORF | mRNAs (↑ NC) | mRNAs (random) | 10732/26039 (41.2%) | 3947/26039 (15.2%) | 4 | 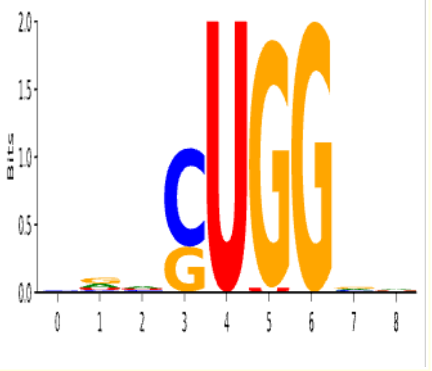  | 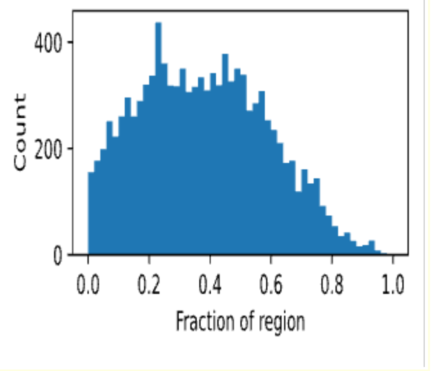  | 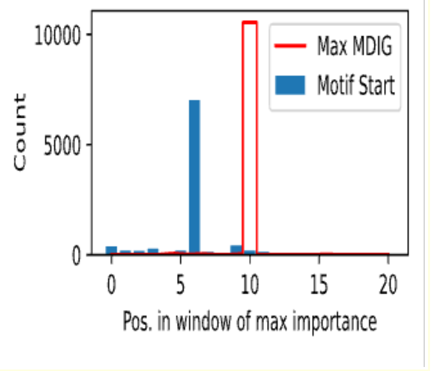  | 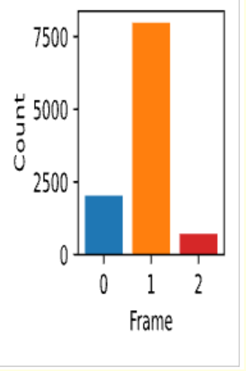  | 8.89E-89 | 3.90E-91 | 7.16 |
| 11 | ORF | lncRNAs (↑ PC) | lncRNAs (random) | 17070/24881 (68.6%) | 13797/24881 (55.5%) | 8 | 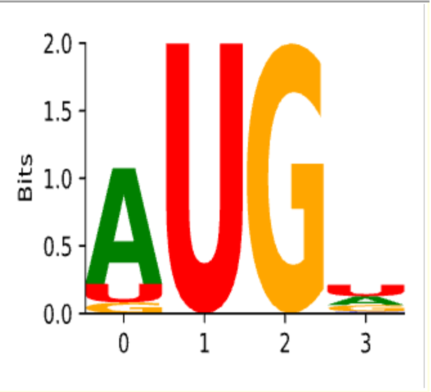  | 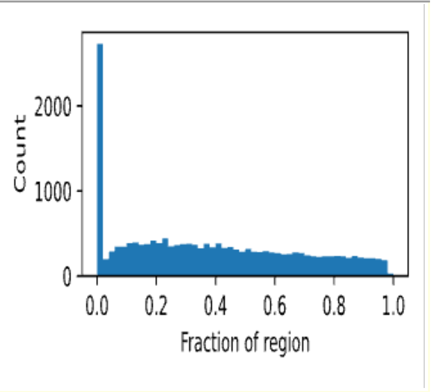  | 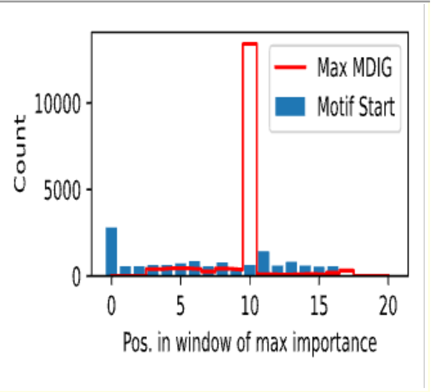  | 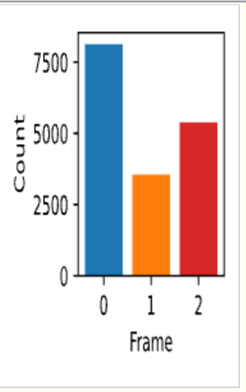  | 2.96E-25 | 1.30E-27 | 5.28 |

| Motif # | Region | Positive Set (sites) | Negative Set (sites) | Pos. Sites | Neg. Sites | Cluster | Logo | Start site in region | Start site in window | Offset from ORF | E-value | p-value | Information |
|---|---|---|---|---|---|---|---|---|---|---|---|---|---|
| 12 | ORF | lncRNAs (↑ NC) | lncRNAs (random) | 19282/24881 (77.5%) | 15880/24881 (63.8%) | 6 | 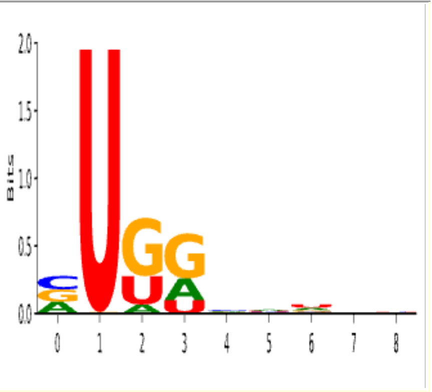 | 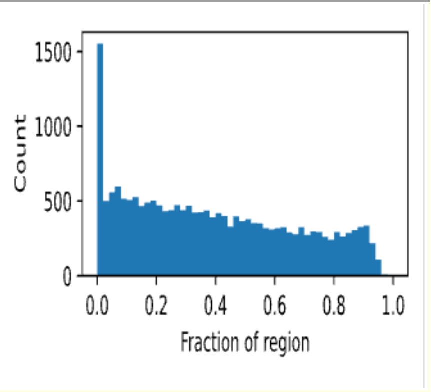 | 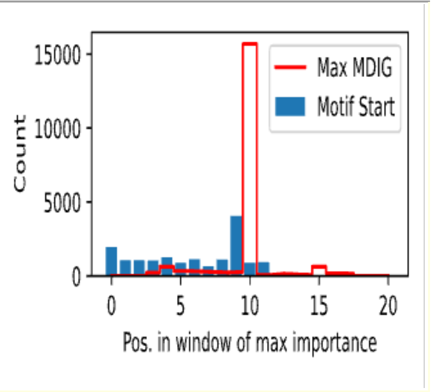 | 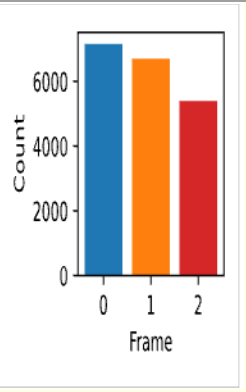 | 1.87E-22 | 8.20E-25 | 3.67 |
| 13 | ORF | mRNAs (↑ PC) | lncRNAs (↑ PC) | 190/26039 (0.7%) | 29/24881 (0.1%) | 9 | 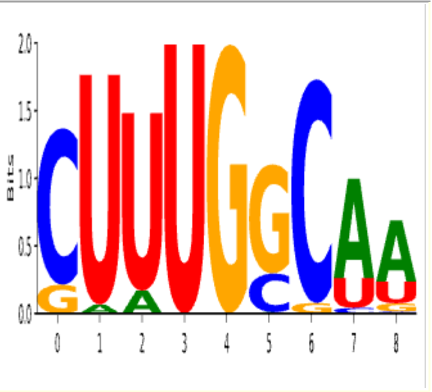 | 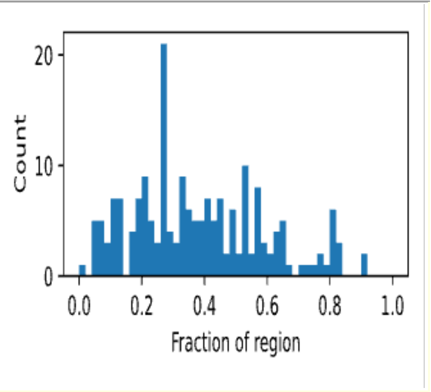 | 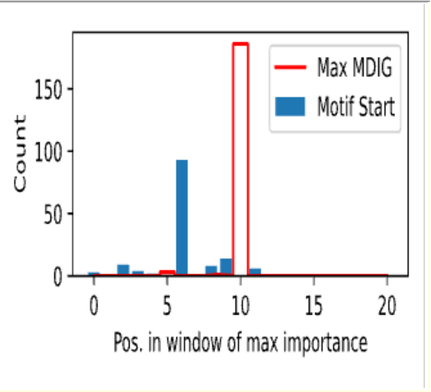 | 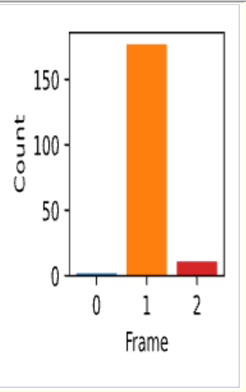 | 5.70E-04 | 2.50E-06 | 13.11 |
| 14 | ORF | mRNAs (↑ NC) | lncRNAs (↑ NC) | 14928/26039 (57.3%) | 8725/24881 (35.1%) | 10 | 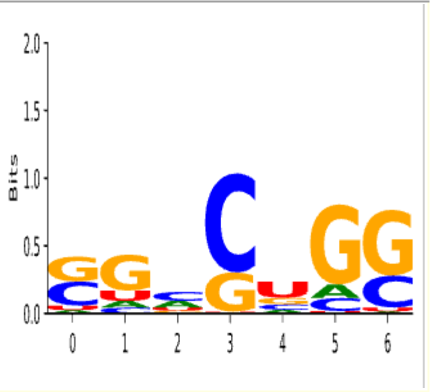 | 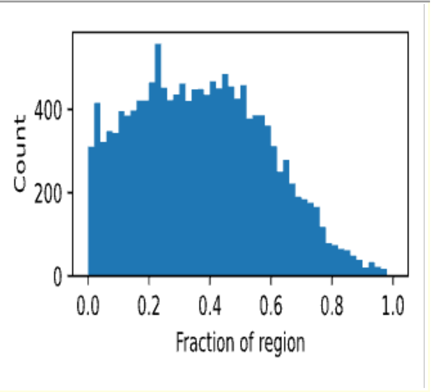 | 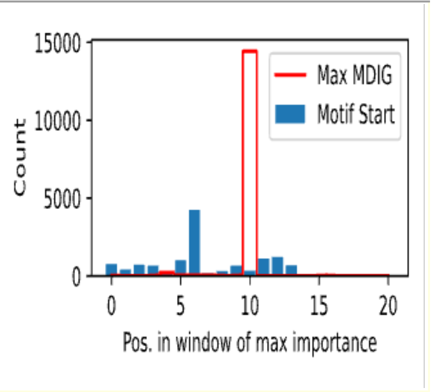 | 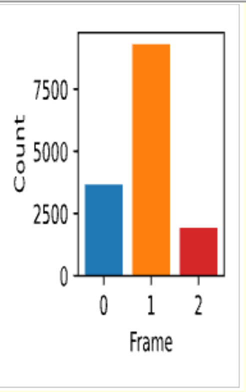 | 4.10E-53 | 1.80E-55 | 3.86 |
| 15 | ORF | lncRNAs (↑ NC) | mRNAs (↑ NC) | 6558/24881 (26.4%) | 5842/26039 (22.4%) | 8 | 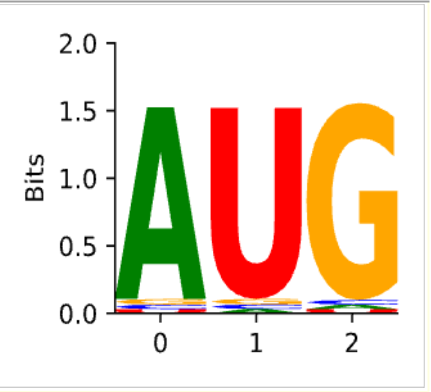 | 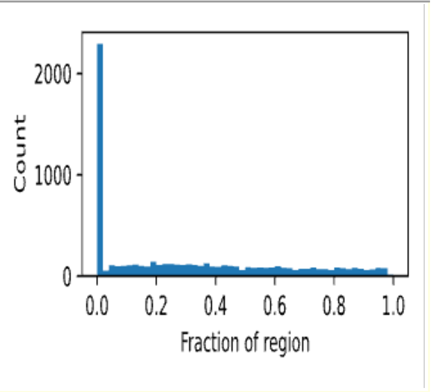 | 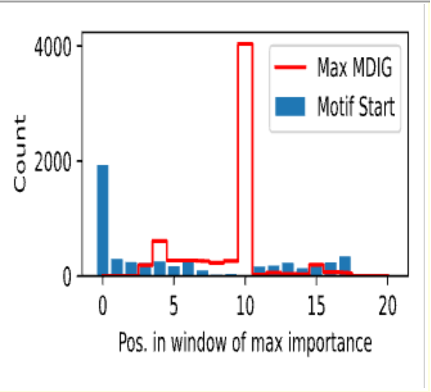 | 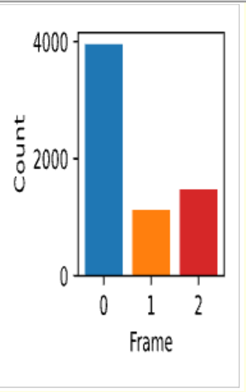 | 5.93E-04 | 2.60E-06 | 4.60 |
